# Preclinical Pediatric Molecular Analysis for Therapy Choice (MATCH)

**DOI:** 10.1101/2024.07.22.604648

**Authors:** Kaley Blankenship, Ti-Cheng Chang, Yiping Fan, Brittney Gordon, William C. Wright, Matthew Kieffer, Asa Karlstrom, Jongrye Jeon, Anand G. Patel, Jason Dapper, William V. Caufield, Burgess B. Freeman, Sara Federico, Michael R. Clay, Gang Wu, Xin Zhou, Lauren Hoffmann, Paul Geeleher, Michael A. Dyer, Justina McEvoy, Elizabeth Stewart

**Affiliations:** Department of Oncology, St. Jude Children’s Research Hospital, Memphis, Tennessee 38105, USA; Center for Applied Bioinformatics Shared Resource, St. Jude Children’s Research Hospital, Memphis, Tennessee 38105, USA; Departments of Developmental Neurobiology, St. Jude Children’s Research Hospital, Memphis, Tennessee 38105, USA; Departments of Computational Biology, St. Jude Children’s Research Hospital, Memphis, Tennessee 38105, USA; Preclinical Pharmacokinetics Shared Resource, St. Jude Children’s Research Hospital, Memphis, Tennessee 38105, USA; Department of Pathology, University of Colorado, Anschutz Aurora, CO 80045

## Abstract

Pediatric solid tumors arise from diverse tissues during development and exhibit a wide range of molecular, cellular and genetic features. This diversity, combined with the low incidence of pediatric cancer makes it increasingly difficult to personalize therapy for individual patients based on the unique features of their tumors. Therefore, well-credentialed preclinical models that capture the diversity and heterogeneity of pediatric solid tumors are essential for identifying molecular targeted therapeutics for precision medicine. Here, we report 281 orthotopic patient derived xenografts (O-PDXs) from 224 patients representing 24 different types of pediatric solid tumors. We have performed genomic characterization of the O-PDXs and compared them to their corresponding patient tumors. To demonstrate the feasibility and utility of using such a diverse collection of O-PDXs in preclinical studies, we performed a preclinical pediatric precision medicine trial based on the NCI-COG Pediatric MATCH trial enrollment criteria. We also tested molecular targeted therapy for a novel oncogenic fusion recently reported in pediatric melanoma and precision drug delivery using nano-liposomal irinotecan. Our studies demonstrate the value of large, well-credentialed preclinical models for future precision medicine in pediatric oncology using single agents, drug combinations and novel drug formulations.

**Translational Relevance:** This study demonstrates the value of utilizing fully characterized preclinical models of pediatric solid tumors to evaluate the response to precision medicine approaches. Our results demonstrate the importance of performing comprehensive preclinical testing using multiple orthotopic patient derived xenografts to validate and prioritize vulnerabilities identified through genomic or integrated analyses which can be translated into clinical trials. Importantly, this study identified combinations using nano-liposomal irinotecan in a precision drug delivery approach that may benefit pediatric solid tumor patients. In addition, all models and their associated data are made freely available to the scientific community through the Childhood Solid Tumor Network.

## Introduction

Preclinical models are increasingly important in oncology research with the expansion of precision medicine^1,2^. For example, NTRK fusion-positive cancers are sensitive to TRK inhibitors (e.g. larotrectinib) irrespective of cancer type, fusion partner or patient age^3^. Despite these remarkable advances in precision medicine, future success is dependent on our ability to identify actionable genetic mutations and biomarkers that can improve and predict clinical responses. Indeed, NTRK fusion-positive cancers are rare in children (<0.1%) so there would have been very little chance of identifying the TRK inhibitor exceptional responders in a standard all-comers phase I trial. Preclinical models that span the diversity of cancer types and heterogeneity within those diseases are invaluable for identification and validation of actionable genetic lesions and biomarkers of response to molecular targeted therapies.

Diverse and well-credentialed preclinical models are even more important in pediatric cancer because childhood cancer is rare^4^. Ideally, new therapeutic combinations should be validated and compared to conventional regimens in preclinical models before launching new clinical trials that can take 3-5 years to complete^5–9^. Moreover, scheduling of agents may differ from adults. For example, irinotecan is typically administered as a single large dose once every course of treatment in adults, whereas in children, smaller doses of irinotecan are administered over several days to extend the duration of tumor exposure and reduce the toxicity^9^. Thus, preclinical models are essential in pediatric cancer research for identifying actionable mutations or biomarkers for molecular targeted therapeutics, for evaluating the best drug regimens to move into the clinic and for optimizing drug dose, schedule and delivery method to improve outcomes while minimizing toxicity.

Previously, 67 orthotopic patient-derived xenografts (O-PDXs) representing 12 different pediatric solid tumor types were characterized and released to the international biomedical research community through the Childhood Solid Tumor Network (www.stjude.org/CSTN/)^7^. Here we report the expansion of that resource to 281 representing 24 different types of pediatric solid tumors. Some of these O-PDXs are so rare that no pre-existing cell lines or genetically engineered mouse models are available for laboratory studies. Characterization of these models includes whole-genome sequencing (WGS), whole-exome sequencing (WES), DNA methylation, and RNA sequencing (RNA-seq) of the patient tumor and matched O-PDXs. WGS and WES of the patient germline were also performed. Cellular features were validated with histopathologic analysis by a pediatric solid tumor pathologist and transmission electron microscopy.

In addition to these carefully credentialed orthotopic models, we have established an approach to preclinical testing that recapitulates clinical trials in patients (phase I, II, III)^6,10–17^. Pharmacokinetics (PK) are done to ensure that the drug exposure in mice is similar to that in patients including the schedule and number of chemotherapy courses. To reduce bias, mice are randomized to treatment groups and studies are benchmarked against a current standard of care. Previous studies have incorporated molecular targeted therapies into combination regimens that are clinically relevant^6,10–17^. We also use O-PDXs from multiple patients to represent the clinical diversity for each disease with an emphasis on the most aggressive recurrent tumors.

In 2017, the National Cancer Institute (NCI) and Children’s Oncology Group (COG) opened the pediatric version of the Molecular Analysis for Therapy Choice (MATCH) trial with 8 different study groups targeting a predefined set of gene mutations. The study is designed to add or remove study groups based on scientific rationale and/or clinical results. To enroll on pediatric MATCH, DNA and RNA is sequenced to identify genetic abnormalities that may be targeted by a drug in one of the study groups. If pediatric phase I data was available for a MATCH drug, the recommended phase II dose (RP2D) in children is used. If such data was not available, 80-100% of the adult RP2D is used. The NCI-COG Pediatric MATCH trial is a phase II study so the primary endpoint is overall response rate for each treatment group. In an interim analysis in 2019, 390 patients had undergone molecular testing and 10% of those patients were ultimately enrolled in a study group^18^. Anticipated enrollment is 1,000 patients so accrual is expected to continue over several years.

Here, we use our expanded cohort of credentialed O-PDX models of pediatric solid tumors and standardized preclinical testing paradigm to determine the feasibility of performing a preclinical pediatric MATCH trial. In total, we identified 49 O-PDXs from 39 patients with matches among our cohort using the criteria from the NCI-COG-Pediatric MATCH trial. The enrollment criteria for pediatric MATCH includes patients who failed initial therapy^18^ so we prioritized O-PDX models from patients with recurrent or refractory disease. We performed plasma and tumor pharmacokinetics for each MATCH study drug in order to determine the murine equivalent dose (MED) and the tumor penetration. Randomized preclinical phase II studies were performed for each MATCH study drug including genetic control cohorts and appropriate standard of care regimens (SOC). We extended our studies beyond the NCI-COG-Pediatric MATCH treatment groups to include emerging precision oncology agents. Taken together, our results demonstrate the feasibility of using O-PDXs for optimizing molecular targeted therapeutic regimens in standardized preclinical studies to provide scientific justification and translational relevance for precision oncology clinical trials in children.

## Results

### Pediatric Solid Tumors Represent Diverse Developmental Origins

In 2010, a biology protocol was opened called Molecular Analysis of Solid Tumors (MAST:NCT01050296) that offers every solid tumor patient at St. Jude Children’s Research Hospital the opportunity to consent to donate excess tumor tissue for orthotopic implantation into immunocompromised mice. Over the first 10 years (2010-2020), 708 patients consented to MAST and donated 795 tissue specimens. Among those 795 specimens, 331 (42%) grew at least one O-PDX subline that was passaged and cryopreserved. We have now carried out detailed characterization on 281 of those O-PDXs representing 24 different tumor types (Fig. 1A and Table S1). There were 192 (68%) O-PDXs from the primary site and 89 (32%) from metastatic sites and 111 (40%) were from recurrent tumors (Fig. 1A and Table S1). We performed whole genome sequencing (WGS) and whole exome sequencing (WES) of the patient tumor, patient germline and O-PDX. We also performed RNA-seq and DNA methylation (DNAm) profiling (Infinium MethylationEPIC BeadChip) of the patient tumor and O-PDX. Histopathologic review was performed by a pediatric solid tumor pathologist on all O-PDXs and their corresponding patient tumor. Some tumors have additional characterization including genetic clonal analysis using custom-capture and deep sequencing, single nucleus RNA-seq (snRNA-seq), ChIP-seq and proteomics/phosphoproteomics^5–8,19^. All O-PDXs are shared free of charge with no obligation to collaborate through the Childhood Solid Tumor Network (www.stjude.org/CSTN/) and associated data are also publicly available in an unrestricted data portal (https://cstn.stjude.cloud/). To date, we have filled 684 requests (>1900 vials) from 281 investigators at 124 institutions across 18 countries.

**Figure 1.**
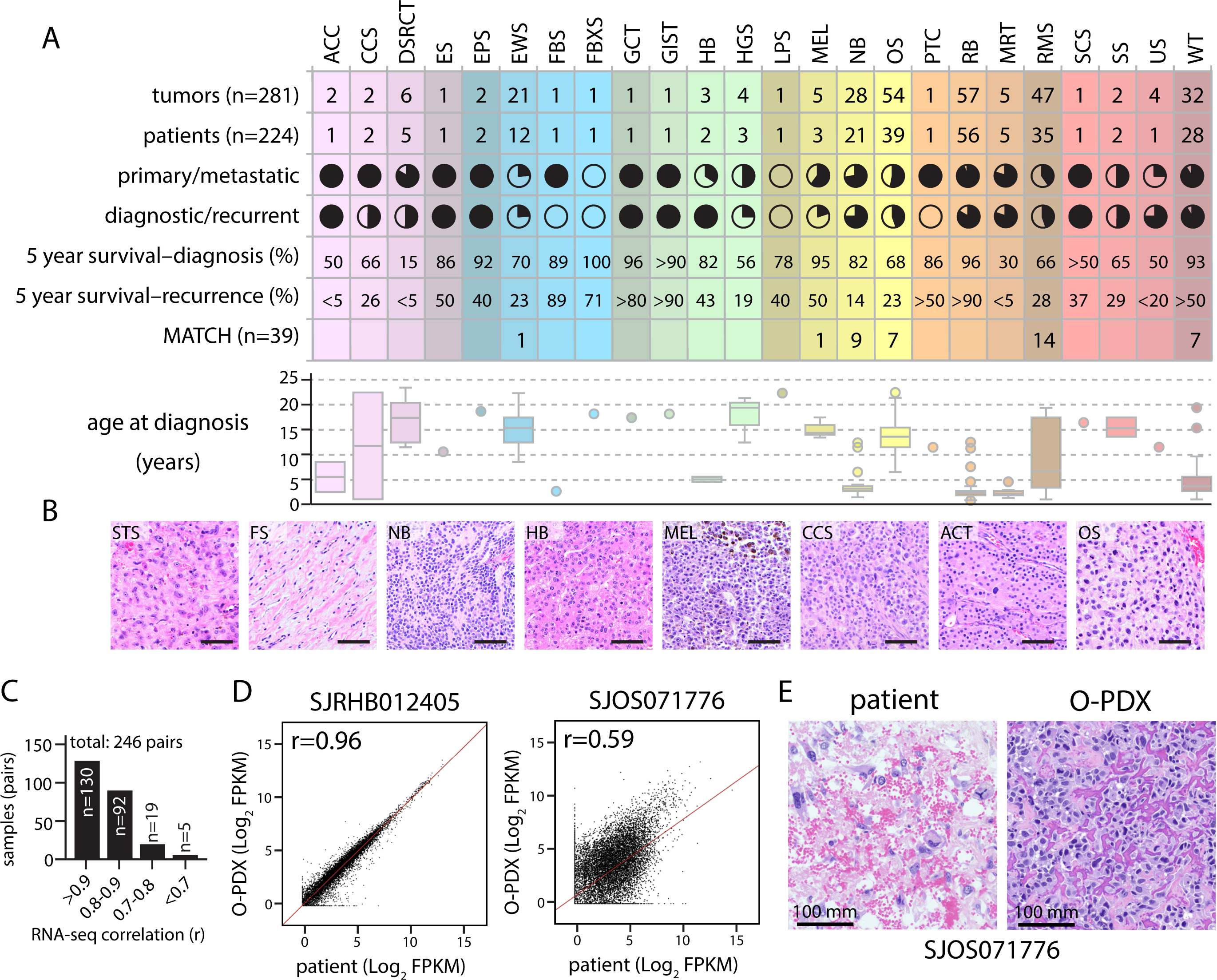
A diverse collection of credentialed pediatric solid tumor O-PDXs. **A)** Table summarizing the diversity of tumor types and clinical information. Filled portion is primary and transparent is metastatic. Filled portion is diagnostic and transparent is recurrent. **B)** Micrographs of representative hematoxylin and eosin (H&E) stained sections of the specified subset of O-PDXs. **C)** Bar plot of RNA-seq correlation of matched patient tumors and corresponding O-PDXs showing correlation ≥ 0.8 in 90% of the analyzed pairs. **D-E)** Correlation analysis of the RNA-seq data for a representative primary tumor and xenograft pair with a high (r = 0.96; SJRHB012405, rhabdomyosarcoma) and low (r = 0.59; SJOS071776, osteosarcoma) coefficient (red line). H&E micrographs are shown in (E) for the osteosarcoma pair with a lower correlation coefficient which demonstrate extensive infiltration of normal cells in the patient tumor which were lacking in the paired O-PDX. Abbreviations: ACC, adrenal cortical carcinoma; CCS, clear cell sarcoma; DSRCT, desmoplastic small round cell tumor; ES, embryonal sarcoma; EPS, epithelioid sarcoma; EWS, Ewing sarcoma; FBS, fibrosarcoma; FBXS, fibromixoid sarcoma; GCT, germ cell tumor; GIST, gastrointenstinal stromal tumor; HB, hepatoblastoma; HGS, high grade sarcoma; LPS, liposarcoma; MEL, melanoma; NB, neuroblastoma; OS, osteosarcoma; PTC, parathyroid carcinoma; RB, retinoblastoma; MRT, malignant rhabdoid tumor; RMS, rhabdomyosarcoma; SCS, spindle cell sarcoma; SS, synovial sarcoma; US, unspecified sarcoma; WT, Wilm’s tumor; FPMK, fragments per kilobase of transcript per million. Scale bars: 50 mm unless noted.

The 24 different tumor types have distinct histologic features that are consistent with their diverse developmental origins (Fig. 1B and CSTN data portal). This diversity is also represented in their transcriptomes (Fig. S1) and DNAm profiles (Fig. S2). We analyzed the correlation for the bulk RNA-seq between patient tumor and matched O-PDX. There was strong correlation with 90% of pairs having a correlation coefficient of 0.8 or greater (Fig. 1C, D and Table S1). For those with low correlation coefficients (e.g. SJOS071776) histopathologic review showed that they had extensive infiltration of normal cells in the patient tumor that were lacking in the O-PDX (Fig. 1E). Pathway analysis (Enrichr) of the top 100 most enriched genes for the SJOS071776 patient tumor relative to the corresponding O-PDX was consistent with immune cell infiltration (GO process 2023: inflammatory response, Table S1). Analysis of DNAm profiles using Euclidean distance showed that nearly all O-PDXs are closer to their paired patient tumor compared to other patient or O-PDX tumors of the same lineage (Fig. S2 and Table S1). It has been shown previously that the process of transplanting a patient tumor into an immunocompromised mouse leads to clonal selection and clonal evolution with each passage^5,7^. Therefore, the genomic landscape can change over time in O-PDXs relative to the patient tumor^5,7^. The most common recurrent genetic lesions are presented for the patient tumor and matched O-PDX on the CSTN data portal and raw sequence data files are available on the St. Jude Cloud (https://www.stjude.cloud/). To determine if the likely pathogenic somatic SNVs and indels are preserved in the O-PDX relative to the patient tumor, we used the St. Jude clinical genomics pipeline used in the genomes for kids (G4K) clinical trial (NCT02530658)^20^ for a subset (n=81) of our patient/O-PDX pairs. The St. Jude Medal Ceremony clinical genomics pipeline) (https://university.stjude.cloud/docs/pecan/pecan-pie/) assigns gold, silver or bronze medals for coding and splice-related variants in disease predisposition genes. We found that 71 SNVs and 24 indels were found in the patient tumor and O-PDX (Table S2). There were 55 SNVs and 26 indels found in the patient tumor but not the O-PDX and 84 SNVs and 18 indels found in the O-PDX but not the patient tumor (Table S2 and Fig. S2) which are likely related to clonal changes or difference in detection of mutations due to patient tumor purity^5,7^.

### Identification of NCI-COG Pediatric MATCH Mutations in O-PDXs

To determine if any of the O-PDXs had mutations that would make a patient eligible for the NCI-COG Pediatric MATCH trial, we cross referenced our genomic data to the patient eligibility criteria for each MATCH drug group. We identified 49 O-PDXs from 39 patients representing 6 different tumor types (Ewing sarcoma, melanoma, neuroblastoma, osteosarcoma, rhabdomyosarcoma, Wilms tumor) eligible for the NCI-COG-Pediatric MATCH protocol (Fig. 2). While germline results are available for 98% of the patients in our cohort, matches in the O-PDXs were calculated based on tumor alterations as is done in the clinical trial. There were genetic lesions in 13 genes (*ALK, CDK4, CCND2, CCND3, FGFR1, FGFR4, KRAS, NRAS, HRAS, PI3CA, NF1, PTEN, MAP2K1*) that matched 6 different drugs (ensartinib, erdafitinib, palbociclib, LY3023414, selumetinib, ulixertinib). The most common preclinical MATCHs in our cohort were MAPK pathway activating mutations for the selumetinib treatment group (Fig. 2) which is consistent with reported data for patients enrolled in the Pediatric MATCH trial^21^. As a positive control for precision oncology, we used an infantile fibrosarcoma (IFS) O-PDX model with *ETV6-NTRK3* fusion and generated 2 BaF3 cell lines that ectopically express druggable oncogenic fusions^22^. The BaF3^ALK^ line has an *EML4-ALK* fusion and the BaF3^NTRK^ line has an *ETV6-NTRK3* fusion. ALK and NTRK fusion oncoproteins have been effectively targeted with ALK inhibitors (ceritinib, ensartinib) and NTRK inhibitors (larotrectinib, entrectinib), respectively^3,23,24^. As done previously^22^, these lines are dependent on the oncofusion proteins and are no longer dependent on IL-3 for growth (data not shown). The BaF3^ALK^ and BaF3^NTRK^ lines were used as positive controls for all subsequent preclinical precision medicine trials.

**Figure 2.**
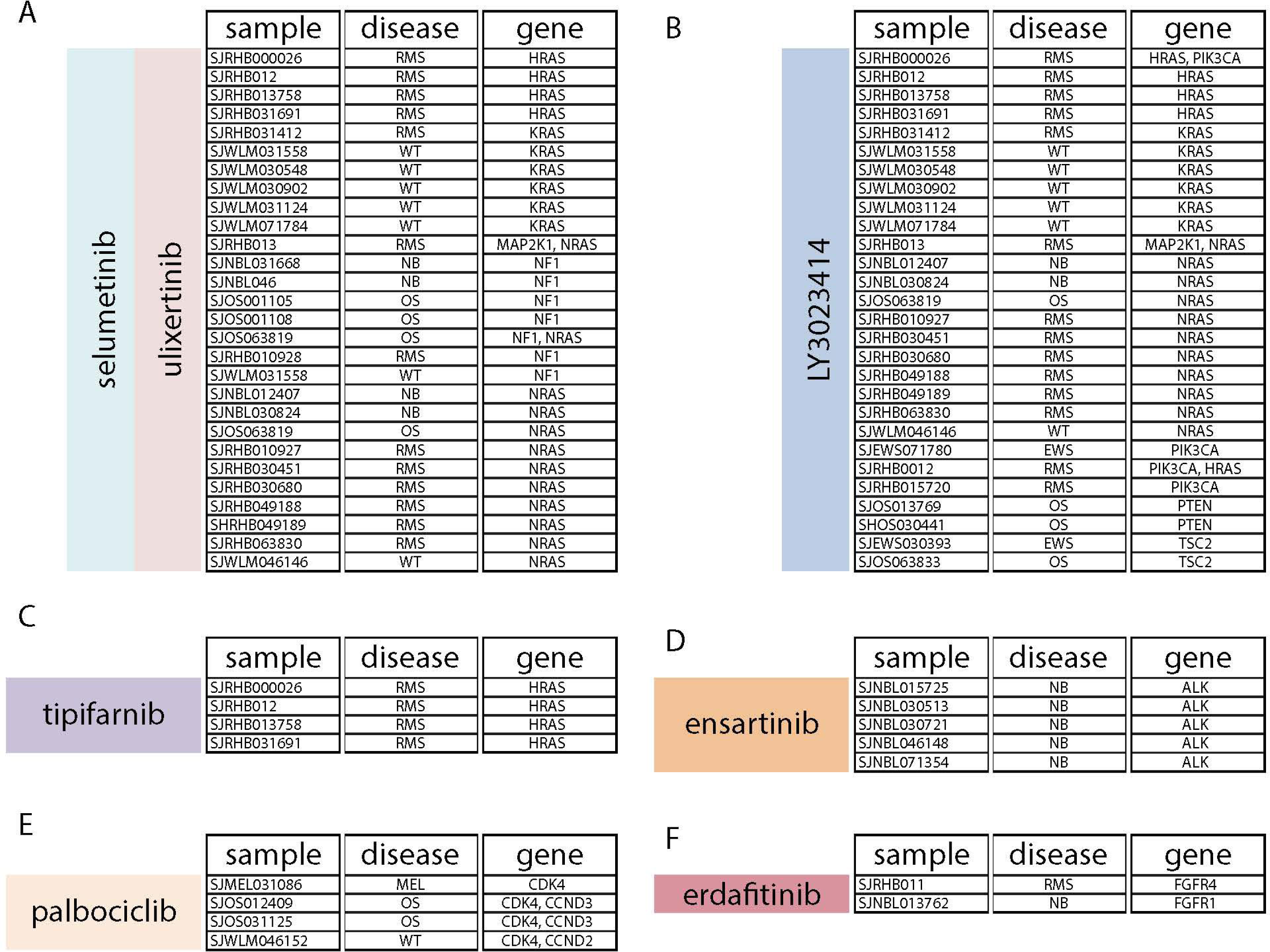
Preclinical pediatric MATCH samples. **A)** MATCH samples and gene mutations for selumetinib and ulixertinib to target MAP kinase pathway perturbations. Two samples had multiple 2 gene mutations that qualify for the protocol. **B)** MATCH samples and gene mutations for LY3023414 to target MAP kinase and PI3K pathway mutations. Three samples had multiple gene mutations that qualify for the protocol. **C)** HRAS specific samples targeted by tipifarnib. **D)** ALK specific samples targeted by ensartinib. **E)** The CDK4/6 inhibitor (palbociclib) is used for RB wild type tumors that have amplification of CDK4 and/or CCND2/3 in our cohort according to the clinical trial guidelines. **F)** FGFR specific samples targeted by erdafitinib. Abbreviations: RMS, rhabdomyosarcoma; WT, Wilm’s Tumor; NB, neuroblastoma; OS, osteosarcoma; MEL, melanoma.

### Preclinical Pharmacokinetics for MATCH Drugs

In order to accurately dose the MATCH drugs, we performed plasma pharmacokinetics (PK) in immunocompromised mice. We also performed tumor PK to estimate the tumor penetration at clinically relevant doses. The murine equivalent dose (MED) was calculated to mimic the plasma exposure in humans as done previously^5–8^. In total, we performed plasma and tumor PK for 8 drugs (selumetinib, ensartinib, larotrectinib, erdafitinib, tazemetostat, olaparib, LY3023414, palbociclib) (Fig. 3A-C and supplemental information). We also performed PK for standard of care (SOC) drugs to benchmark our preclinical efficacy studies (Fig. 3D and supplemental information). Detailed PK reports are publicly available for each drug (https://cstn.stjude.cloud/). Using our PK data, we were able to recapitulate the dose and schedule for MATCH treatment groups and relevant disease-specific SOC regimens (Fig. 3E and supplemental information). Pharmacodynamic (PD) assays were performed to demonstrate that the MATCH drugs penetrated the tumor in the orthotopic location and altered protein and/or pathway function (Fig. S3 and supplemental information).

**Figure 3.**
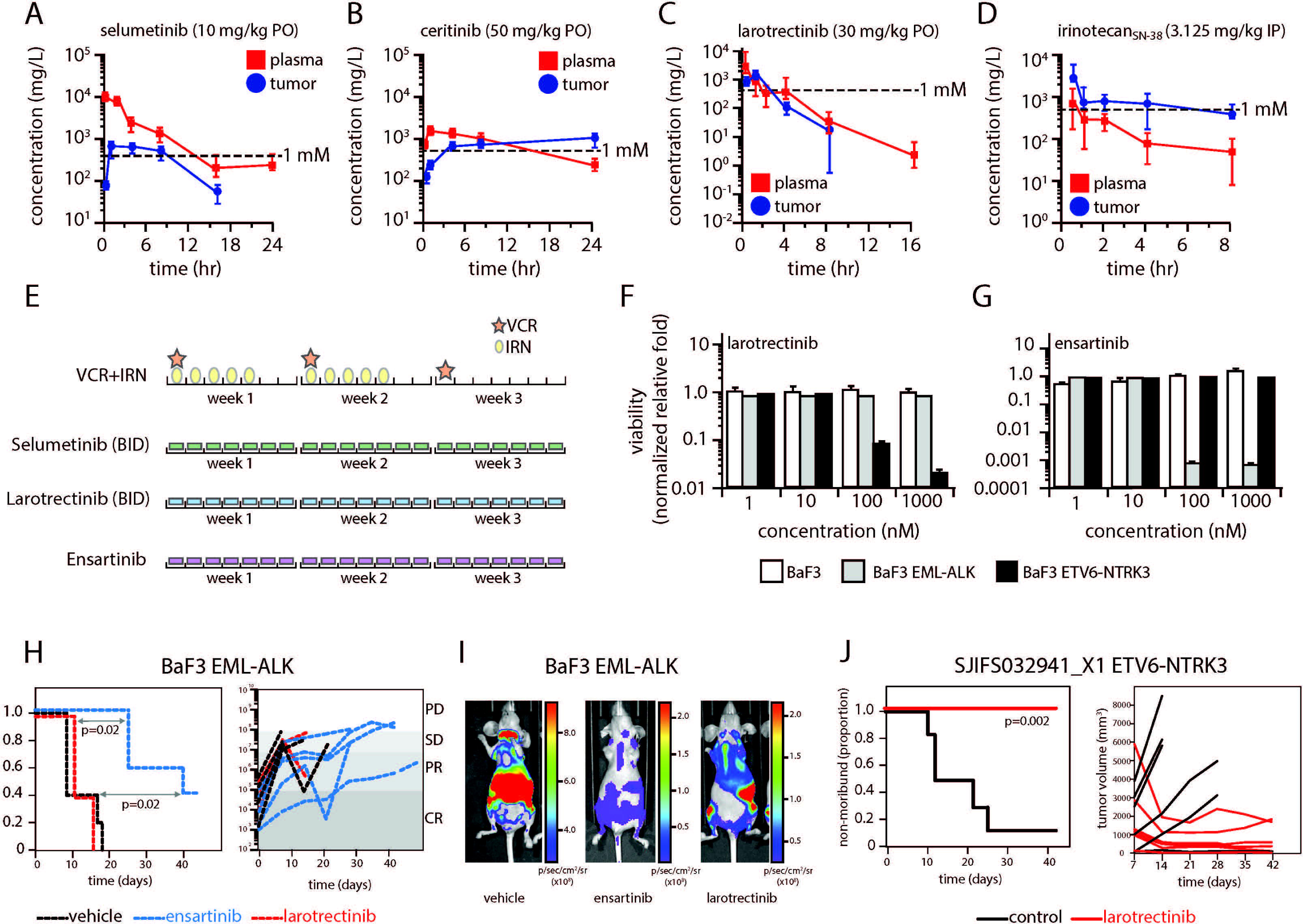
Preclinical pharmacokinetics for plasma and tumor exposure. **A-D)** Concentration time plots of plasma (red line) and tumor (blue line) levels of selumetinib, ceritinib, larotrectinib and irinotecan (SN-38) in O-PDX models. **E)** Diagram of the schedule of a representative standard of care regimen (VCR+IRN) and corresponding MATCH treatment group used in patients and in our preclinical studies. **F, G)** Barplots of mean and standard deviation for BaF3 cell line viability following exposure to larotrectinib or ensartinib. **H)** Survival curve and individual mouse level response for the disseminated BaF3 tumor model with EML-ALK oncofusion showing significant response to ensartinib. **I)** Representative images of bioluminescence for mice on the treatment regimens in (H). **J)** Survival curve and tumor volume plot for infantile sarcoma O-PDX model with ETV6-NTRK3 fusion showing significant response to treatment with larotrectinib. Abbreviations: PO, oral dosing; IP, intraperitoneal dosing; VCR, vincristine; IRN, irinotecan; BID, twice daily dosing; PD, progressive disease; SD, stable disease; PR, partial response; CR, complete response.

To demonstrate the feasibility of using molecular targeted therapeutics for targeting specific oncogenic mutations, we used our BaF3 control cell lines. *In vitro*, the BaF3^NTRK^ cells were sensitive to larotrectinib (Fig. 3F) and the BaF3^ALK^ cells had selective sensitivity to ceritinib and ensartinib (Fig. 3G and data not shown). Next, the BaF3 cell lines were labeled with luciferase and 1x10^6^ cells were injected into the tail vein for a disseminated tumor model or subcutaneously for a localized tumor model. Ensartinib or larotrectinib was administered and tumor cell response monitored by bioluminescence every week. In these models, we generally achieved more efficient targeting of BaF3^ALK^ tumors with ensartinib than BaF3^NTRK^ with larotrectinib in culture and *in vivo* (Fig. 3F-I and Table S3); however, we found the BaF3 lines to be faster growing and more disseminating compared to O-PDX models. We therefore included additional preclinical experiments using our IFS O-PDX model with *ETV6-NTRK3* fusion along with a 3T3 cell line expressing *EML4-NTRK3* fusion^25^. Both models showed favorable responses to larotrectinib compared to control (p=0.002 and p=0.009 respectively, Fig. 3J and Table S3).

### Preclinical Pediatric MATCH

Having demonstrated the feasibility of targeting oncogenic mutations in preclinical models using ALK and NTRK inhibitors, we performed preclinical phase II (P-PII) studies^5–8^ for MATCH drug groups using our O-PDX models (see Fig. 2). Importantly, models were selected with mutations and as controls we used O-PDXs that were the same tumor type but lacked the MATCH mutation (Table S4 and supplemental information). O-PDX cells were luciferase labeled as described previously and orthotopically injected into nude mice^5–8^. Animals were randomly enrolled to individual treatment groups using pre-defined enrollment criteria^5–8^. The P-PII for each drug group included placebo and SOC regimens for comparison with an average of 5 mice per treatment group. As mentioned above, the SOC regimens were disease specific and dose and schedule were matched to those used in patients (supplemental information). In total, 16 P-PII studies were performed with 172 mice. Four 3-week courses of chemotherapy were administered (17,864 individual doses) and bioluminescence was monitored weekly for a total of 1,911 images. Disease response was measure using predefined criteria for progressive disease (PD), stable disease (SD), partial response (PR) and complete response (CR) as done previously^5–8^.

Among all the preclinical phase II studies, we identified one specific single agent response. An osteosarcoma tumor (SJOS001108_X1, Table S1) with a *TSC2* variant of unknown significance (missense p.Q572H, not in the current MATCH eligibility criteria) had prolonged stable disease when treated with an inhibitor of class I PI3K isoforms, mTOR and DNA-PK (LY3023414) relative to SOC (gemcitabine + docetaxel, p=0.001) or another tumor (SJRHB012405_X1) that had a *TSC1* variant (Fig. 4A-D, Table S4). While missense alterations in TSC2 have been shown to be inactivating in many adult tumors^26,27^, this model had an accompanying *CCND2* variant (D191_E4 splice) for which the combination of these has unknown significance in pediatric cancer. Complete absence of the wild type allele for each of these variants was confirmed suggesting loss of heterozygosity. Testing of an *ALK*-mutant neuroblastoma model (SJNBL046148_X1) with single agent ALK inhibitor (ensartinib) failed to show activity (Table S4). *ALK* alterations in neuroblastoma have been shown to have a varied inhibitor response with R1275Q mutations showing sensitivity to crizotinib; however, the most recurrent mutation F1174L is only sensitive at higher doses^28^. Our model harbored the L1196M mutation which is more likely to be resistant to crizotinib yet sensitive to lorlatinib^28^ which may explain the lack of response to the ALK inhibitor ensartinib used in Pediatric MATCH. To relate *in vivo* single agent drug sensitivity to *ex vivo* data using primary cultures of our O-PDX tumors, we performed an additional P-PII study with selumetinib. In particular, rhabdomyosarcoma SJRHB012_Y is sensitive to selumetinib and rhabdomyosarcoma SJRHB011_X is resistant (Fig. 4E, F). *In vivo*, neither tumor was sensitive to selumetinib using the clinically relevant dose and schedule (Table S4). Even when combined with SOC drug regimen for rhabdomyosarcoma (RMS), the addition of selumetinib had only modest improvement in survival in the sensitive tumor (Fig. 4G, H and Table S4). The selumetinib arm of the Pediatric MATCH trial which included 7 RMS patients has recently been published which showed no objective responses^29^. Taken together, these data are consistent thus far with early results from the NCI-COG-Pediatric MATCH trial and highlight the challenges with precision oncology in pediatric cancer^2^. Specifically, exceptional responders such as the *TSC2* variant osteosarcoma tumor treated with LY3023414 are rare but have the potential to transform treatment and outcomes for children with cancer.

**Figure 4.**
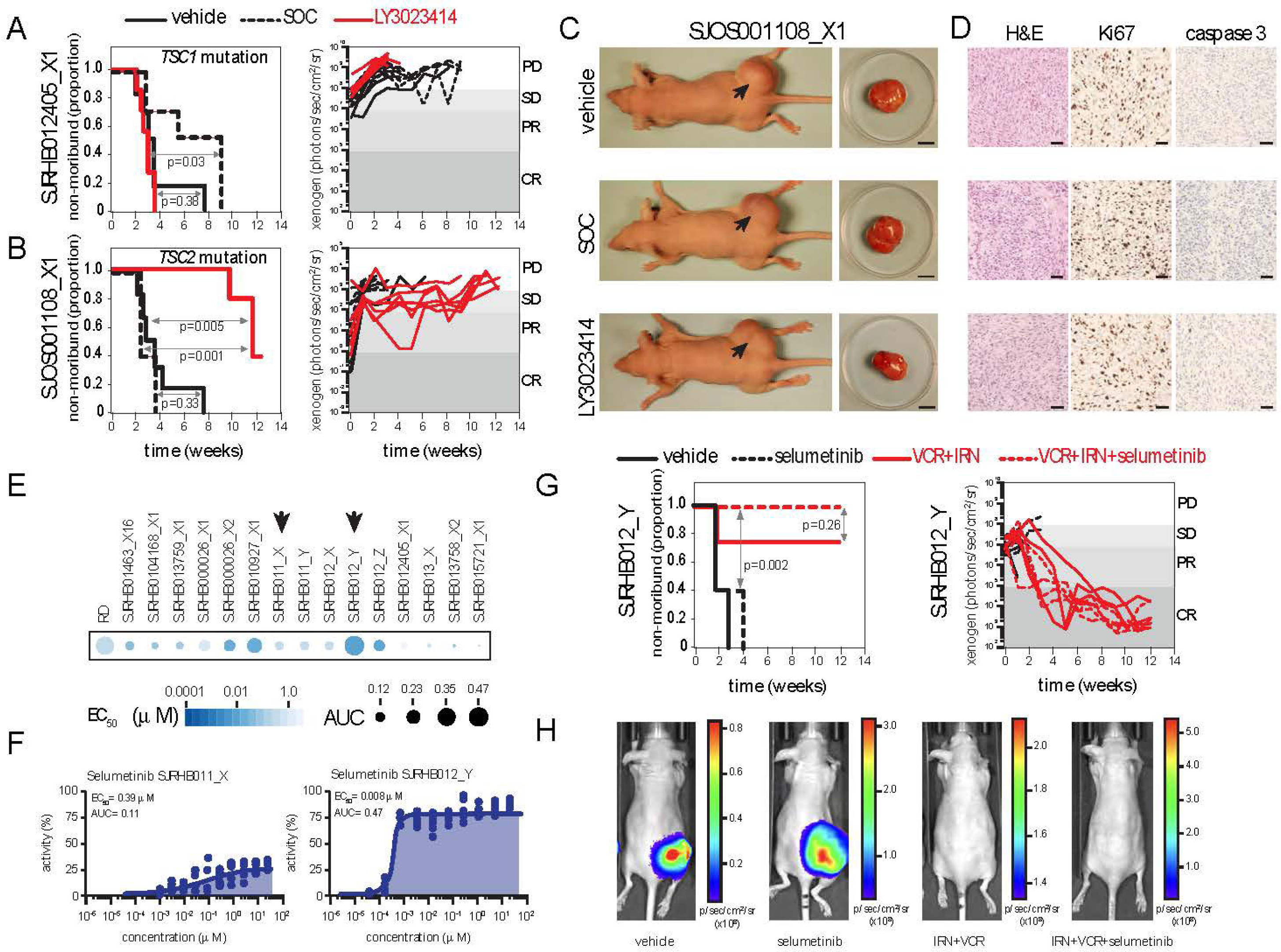
Preclinical phase II study of MATCH drugs. **A, B)** Survival curves and response of individual mice with TSC1 and TSC2 mutation variants to LY3023414 MATCH therapy, placebo and SOC control treatment regimens. **C)** Photographs of representative mice at end of study for each treatment group and their tumor. **D)** Representative histology photographs for each tumor shown in (C) for hematoxylin and eosin (H&E), Ki67 and activated caspase 3. **E, F)** *Ex vivo* drug sensitivity heatmap (E) of primary cultures of rhabdomyosarcoma xenografts. Arrows indicate representative resistant and sensitive O-PDX models used for the *in vivo* study with corresponding dose response curves (F). **G)** Survival curve and response of individual mice for vehicle control, selumetinib alone, VCR+IRN and VCR+IRN+selumetinib. **H)** Representative xenogen images for each treatment group shown in (G). Scale bars: C) 5 mm, D) 50 mm. Abbreviations: SOC, standard of care; VCR, vincristine; IRN, irinotecan; AUC, area under the curve.

### Targeting MAP3K8 in Spitzoid Melanoma

In children, the most common form of melanoma is a specific variant called spitzoid melanoma^30–31^. Previous genomic studies revealed that MAP3K8 rearrangements are the most common genetic event in spitzoid melanoma^32^. In all cases reported to date, exons 1-8 of MAP3K8 are retained and exon 9 is replaced with an alternative 3’ end^32^. The protein domain encoded by exon 9 is involved in repressing kinase activity and may also contribute to proteosomal degradation of MAP3K8^33,34^. Therefore, the rearrangements of the MAP3K8 locus in spitzoid melanomas are thought to contribute to tumorigenesis. MAP3K8 is a serine-threonine kinase that activates ERK1/2 through phosphorylation of MEK^35^ and there is evidence in diverse adult malignancies that MAP3K8 may contribute to deregulation of the MEK/ERK pathway^36–38^. We have established an O-PDX from a spitzoid melanoma patient with an in-frame fusion of MAP3K8 exon 8 and GNG2 exon 3 (SJMEL030083_X1) (Fig. 5A-F). This patient has been previously described and had a mixed response to a MEK inhibitor (trametinib)^32^. Genomic profiling confirmed that the O-PDX retained the somatic mutations in the patient tumor (Fig. 5A) including the MAP3K8-GNG2 fusion (Fig. 5B-D). Histopathologic analysis confirmed that it retained cellular features of spitzoid melanoma (Fig. 5E, F). The tumor was labeled with luciferase and injected subcutaneously in nude mice to monitor tumor growth. We noticed that there was a reduction in body weight of mice with SJMEL030083_X1 relative to mice with other O-PDXs including melanoma (SJMEL031086_X4) consistent with cancer cachexia (Fig. 5D and Table S5).

**Figure 5.**
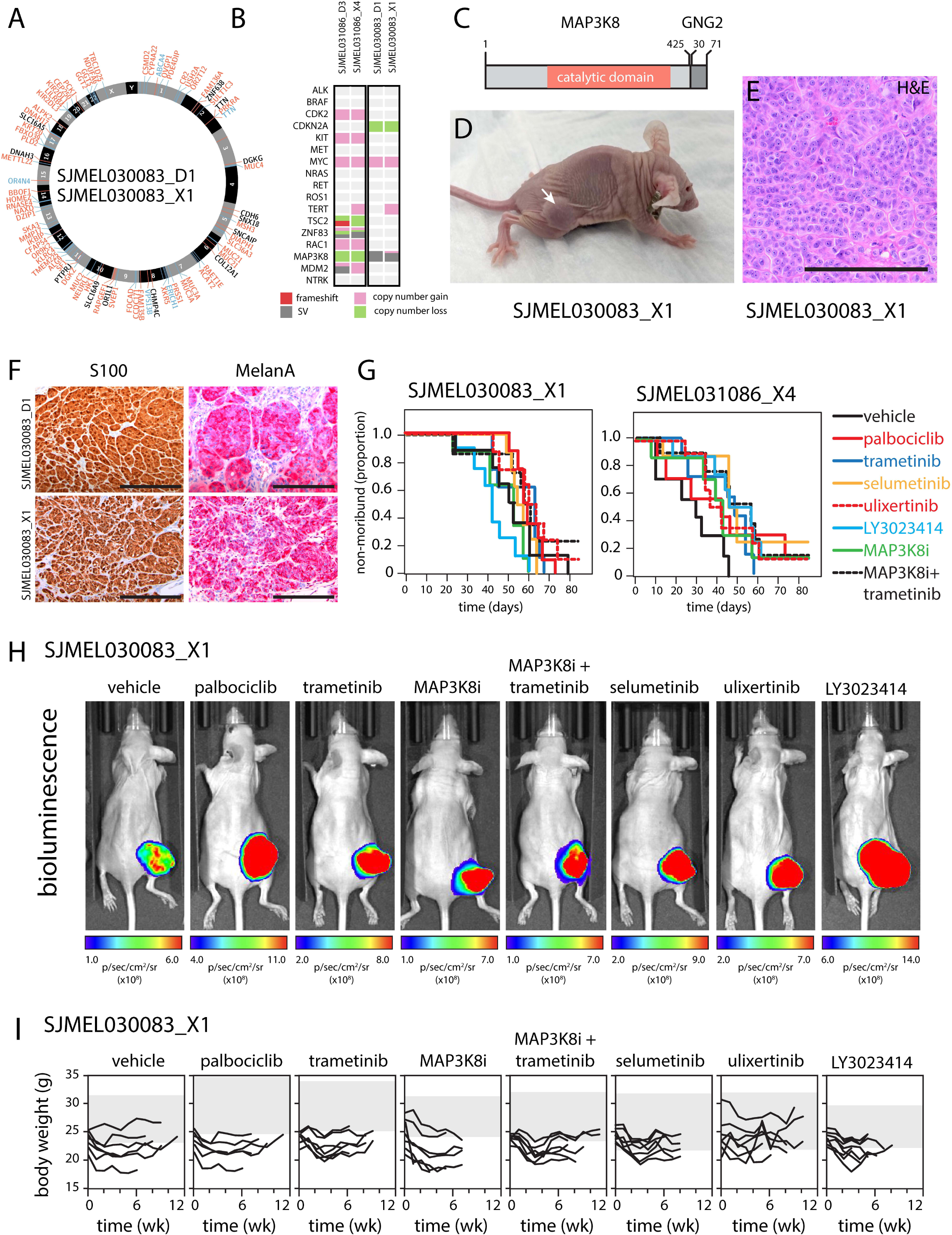
Targeting MAP3K8 in pediatric melanoma. **A)** Circos plot of mutations in SJMEL030083 patient tumor (_D1) and O-PDX (_X1). Mutations in blue are found in patient tumor but not O-PDX, red are found in O-PDX but not patient tumor and black are present in both. **B)** Heatmap of common mutations found in melanoma and other rare pediatric tumors for the two O-PDXs used in the preclinical study. **C)** Drawing of MAP3K8 rearrangement in SJMEL030083_D1 that was confirmed in the O-PDX. **D)** Photograph of a mouse with a flank tumor and weight loss due to cachexia. **E)** Micrograph of hematoxylin and eosin stained melanoma O-PDX. **F)** Micrographs of immunohistochemical stain for S100 and MelanA for the SJMEL030083 patient tumor and matched O-PDX. **G)** Survival curves for each O-PDX for each drug showing no difference in response. **H)** Representative xenogen images of mice on each treatment arm of the study shown in (G). **I)** Line graph of body weight of SJMEL030083_X1 for each treatment group over each course. The gray bar shows the body weight for the control O-PDX that does not have cachexia (SJMEL031086_X4) treated with the same therapy. Ulixertinib had partial rescue of cancer cachexia for the SJMEL030083_X1 model. Scale bars: 200 mm.

MAP3K8 inhibitors are not included in NCI-COG-Pediatric MATCH but there is scientific justification for targeting the kinase itself or the downstream pathway in spitzoid melanoma^32^. Specifically, it has been proposed that the loss of the protein domain encoded by exon 9 may lead to an overactive kinase that is more stable^32^. To test this possibility, we performed preclinical studies with MEK1/2 inhibitors (trametinib, selumetinib), an ERK1/2 inhibitor (ulixertinib), a dual PI3K/mTOR inhibitor (LY3023414) and a CDK4/6 inhibitor (palbociclib). The CDK4/6 inhibitor was included because of previous studies in adult cancer showing that this is a downstream target of MEK/ERK signaling and has efficacy in tumors with oncogenic RAS mutations^39,40^. We also tested a MAP3K8 inhibitor (PubChem:C26H25Cl2FN8) as a single agent and combined with trametinib^41^. Plasma and tumor PK was performed to determine the MED for each drug and our P-PII was carried out using the spitzoid melanoma O-PDX with the MAP3K8-GNG2 fusion (SJMEL030083_X1) as well as a control melanoma O-PDX that lacks any *MAP3K8* rearrangement (SJMEL031086_X4). Animals were randomized to the 8 different treatment arms and tumor growth and response was monitored using bioluminescence every week for 12 weeks (4 courses). Overall, 61/65 (95%) of SJMEL030083_X1 mice had progressive disease while 2/8 mice in the MAP3K8 inhibitor + trametinib group and 1/8 mice in the ulixertinib survived the entire study (Fig. 5G, H and Table S5). In the SJMEL031086_X4 model, 7/59 (12%) survived the entire study with at least one mouse remaining in each of the groups tested except for trametinib alone and untreated control (Fig. 5G, H and Table S5). While average survival days was improved in all groups compared to control, most mice still present at the end of study had significant tumor burden which grew rapidly once stopping therapy (Table S5). Review of the PK and PD data of the MAP3K8 inhibitor showed that drug was able to penetrate the tumor at low levels; however, there were no appreciable changes in relevant downstream markers (ERK, pERK) which may explain the lack of response (Fig. S4 and supplemental information). Melanoma O-PDXs are subcutaneous flank tumors so we validated our bioluminescence data with manual caliper measurements and BioVolume measurements (Table S5). We also measured body weight every week to monitor the cancer cachexia and found that ulixertinib reduced cachexic phenotype of the spitzoid melanoma O-PDX even though it did not improve outcome (Fig. 5I). These data demonstrate the feasibility of performing rapid preclinical studies for tumor types or genetic lesions that are too rare for individual clinical trials or even larger precision oncology basket trials such as NCI-COG-Pediatric MATCH.

### Precision Drug Delivery

Precision medicine in pediatric oncology is usually focused on genomics but there may be opportunities for improving individualized care through non-genomic approaches. For example, novel drug delivery mechanisms can have a significant impact for children with cancer by increasing tumor exposure while reducing systemic side effects that can hamper the delivery of efficacious therapeutic regimens that include molecular targeted agents. To determine if our O-PDX models and preclinical approach can be used to test precision drug delivery, we tested a nano-liposomal formulation of irinotecan (L-IRN)^42,43^. Clinical trials have demonstrated the efficacy of irinotecan in diverse pediatric cancers and the major dose limiting toxicity is diarrhea and neutropenia^44–51^. To overcome the toxicity of irinotecan while maintaining continued tumor exposure, children with cancer are often treated on an extended low-dose schedule (dailyx5 or dailyx5x2) rather than a single large dose as used for adult oncology patients. An alternative approach using L-IRN is promising because 95% of the drug remains encapsulated during circulation reducing systemic exposure while maintaining prolonged exposure in the tumor^50^. This lower systemic exposure led to reduced side effects in adult clinical trials without compromising efficacy^42^. In order to determine if L-IRN can replace conventional IRN in preclinical models, we performed a P-PII study using Ewing sarcoma (EWS) O-PDXs. Previously, we have shown that the combination of temozolomide (TMZ) with IRN and a PARP inhibitor (talazoparib (TAL)) leads to synthetic lethality in EWS models^6^. A recent clinical trial (BMNIRN: NCT02392793) showed that there was evidence of activity of this treatment regimen in recurrent EWS patients but there was significant dose limiting toxicity associated with the triple drug combination using conventional delivery of IRN^51^. Therefore, this is an ideal opportunity for testing precision drug delivery of L-IRN.

As a first step, we performed plasma and tumor PK for IRN and the active metabolite (SN-38) using immunocompromised mice dosed with the aqueous and liposomal formulations. The plasma levels of SN-38 drop quickly for standard IRN and L-IRN after dosing while the levels of SN-38 were maintained in the tumor for several days (>48 hrs) with a single dose of the L-IRN formulation (Fig. 6A, B). Previous studies have shown that approximately 5-10 nM SN-38 is required to achieve synthetic lethality in EWS with TMZ and TAL so this L-IRN formulation may provide sufficient continued exposure in the tumor without the side effects that limited the dose escalation of TMZ in the clinical trial^6^. To test this directly, we performed a P-PII study comparing standard IRN on a low-dose protracted schedule (dailyx5x2) and L-IRN on a weekly schedule (days 1 and 8) in combination with TMZ and TAL (Fig. 6C). To compare to our previous studies^6^, we first performed testing using the EWS ES-8 cell line implanted orthotopically. A total of 45 mice were randomized to 7 treatment groups: placebo, IRN+TMZ (SOC), TAL+IRN, TAL+IRN+TMZ, L-IRN, TAL+L-IRN, TAL+L-IRN+TMZ and treated for 4 courses of therapy (12 weeks total). Substituting L-IRN on days 1,8 for standard IRN on days 1-5 in mice receiving triple drug combinations was well tolerated and similar efficacy was maintained as all mice survived the entire study with the majority achieving a CR (Fig. 6D-F). Notably, TAL+L-IRN was significantly more effective than the SOC regimen of standard IRN+TMZ (p=0.002, Fig. 6D-F and Table S6). There was an improvement in outcome for the combination of TAL+L-IRN compared to TAL+IRN with average survival 84 and 61 days, respectively (p=0.12, Fig. 6D-F and Table S6). While the comparison of days on study was not significantly different between these groups, mice treated with L-IRN all had durable complete responses at least 3 months post therapy compared to the standard IRN group whose tumors regrew once stopping treatment (data not shown). Similar studies were performed with three EWS O-PDXs from our collection with comparable results; 34/38 (89%) of mice treated with L-IRN containing regimens survived the entire study (84 days) and there were no significant differences in survival in any of the EWS O-PDXs when L-IRN was substituted for standard IRN when combined with TAL (Fig. S5 and Table S6). With these encouraging results for L-IRN for the treatment of Ewing sarcoma, we extended our studies to rhabdomyosarcoma. We have previously shown that the combination of IRN with vincristine (VCR) and a WEE1 inhibitor (AZD1775) had improved efficacy for the treatment of RMS compared to the SOC^8^. Specifically, we evaluated 8 treatment groups including VCR+IRN, VCR+L-IRN, VCR+IRN+AZD1775 and VCR+L-IRN+AZD1775 (Table S5). We also compared administration of L-IRN on day 1 versus day1 and day 8 (Table S6). As for Ewing sarcoma, the liposomal formulation of IRN was effective for the treatment of rhabdomyosarcoma when combined with other drugs (Fig. 6G-I and Table S6). Moreover, we showed that dosing of L-IRN on days 1, 8 is superior to dosing on day 1 in combination with VCR (Fig. 6G-I). Taken together, the encouraging results from the clinical trial combined with this precision drug delivery preclinical data suggests that the combination of L-IRN with TAL and TMZ may be more effective in patients due to higher intra-tumoral concentrations and potentially fewer systemic side effects and further clinical investigation is warranted.

**Figure 6.**
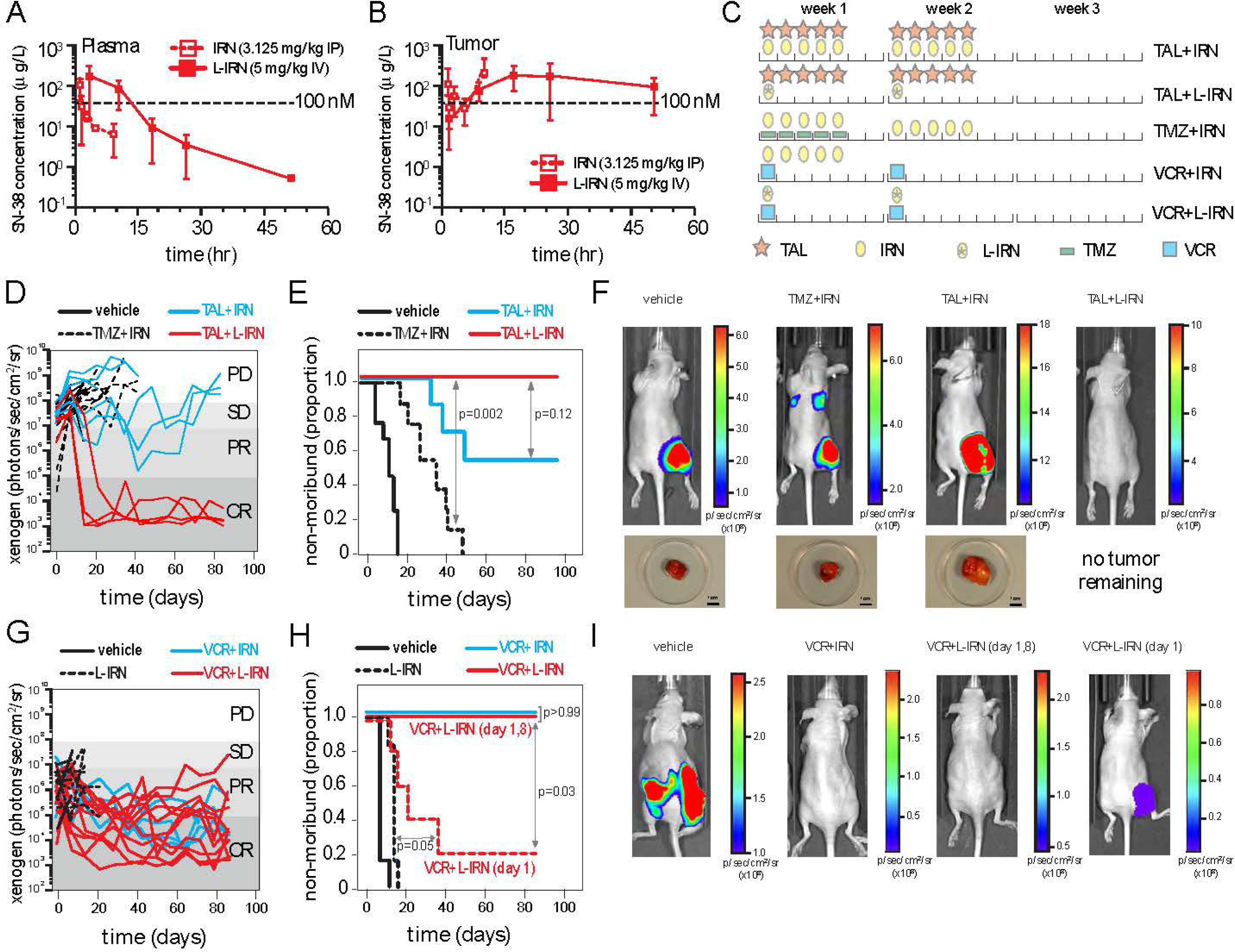
Precision medicine using a liposomal formulation of IRN. **A, B)** Concentration versus time plot of irinotecan (IRN) and SN-38 for the standard formulation (open box) and liposomal formulation (filled box) of IRN (L-IRN). Levels are maintained above 100 nM for at least 48 hours in the O-PDX tumor. **C)** Drawing of the schedules that were used in preclinical studies for administration of standard and L-IRN in patients and in our preclinical studies. **D, E)** Representative survival and individual mouse response for standard and L-IRN in Ewing sarcoma O-PDXs treated with PARP inhibitor (talazoparib) combination chemotherapy. Significant improvement in survival was seen in mice treated with PARP inhibitor + L-IRN compared to standard of care agents (temozolomide + standard IRN, p = 0.002). **F)** Representative xenogen images of mice for the EWS preclinical study shown in (D, E). **G, H)** Survival curves and individual mouse response for standard and L-IRN in RMS O-PDXs showing the improvement in outcome for L-IRN administered on days 1,8 rather than just day 1 alone when combined with vincristine (p = 0.03). Control and single agent L-IRN treated mice are removed from study early due to increased tumor burden. **I)** Representative xenogen images for each treatment group shown in (H). Scale bar: 1 cm. Abbreviations: TAL, talazoparib; IRN, irinotecan; L-IRN, liposomal formulation of irinotecan; TMZ, temozolomide; VCR, vincristine.

## Discussion

Pediatric cancer is rare and there are more than two dozen different types of pediatric solid tumors. In addition, novel molecular targeted therapeutics for oncology are often limited to subsets of patients with particular molecular and/or genetic features. While it is possible to target deregulated pathways across tumor types, often the molecular/genetic stratification further reduces the number of patients that are eligible for clinical trials to establish the efficacy of new oncology drugs. Therefore, preclinical models are increasingly important in pediatric oncology so that the efficacy of new agents can be tested and patient selection based on molecular/genetic profiles can be further refined and prioritized before launching a clinical trial. However, it is important that preclinical models encompass the diverse solid tumor types and the most aggressive refractory/recurrent disease groups that include patients most likely to benefit from new therapeutic combinations. We also recognize the importance of benchmarking novel agents to SOC to gain a better understanding of a significant response. Additionally, we have shown the importance of performing pharmacokinetic/pharmacodynamic studies to guide preclinical dosing based on clinically relevant schedule. This was most evident in testing the novel MAP3K8 inhibitor in our melanoma model with MAP3K8-GNG2 fusion which did not show a significant effect on tumor growth despite continuous twice daily dosing. Penetration of drug in the tumor was confirmed; however, drug levels were not high enough to cause appreciable change in downstream markers of this pathway (ERK, pERK). Additional testing with alternative drugs that effectively target this mutation/pathway is warranted to better understand if this is a promising therapeutic strategy.

### A Diverse, Fully-Credentialed Collection of Pediatric Solid Tumor O-PDXs

Here, we report a collection of 281 O-PDX models of 24 diverse pediatric solid tumors. The majority are from refractory/recurrent tumors and/or metastatic lesions. Detailed characterization of the O-PDXs has confirmed that they retain the molecular, cellular and genetic features of the patient tumors and those data are shared through a freely accessible website (www.stjude.org/CSTN/) and portal on the St. Jude Cloud^52^. The orthotopic location of our O-PDX models is important for three reasons. First, the microenvironment made up of non-tumor cells is different in each location. For example, retinoblastomas grow in the eye where they may encounter astrocytes on the surface of the retina^53^, neuroblastomas grow in the para-adrenal space surrounded by catecholaminergic neurons^54^ and osteosarcoma cells are surrounded by a dense collagen matrix filled with osteoblasts^55^. Second, drug pharmacokinetics may differ across tumor sites such as the eye, bone, liver, muscle and kidney. Some drugs such as topotecan have very rapid uptake in diverse organs following systemic administration while molecular targeted therapeutics such as MDM2 inhibitors have more complex patterns of biodistribution^13^. This is important for selecting the appropriate route of delivery and for providing preclinical efficacy data that are predictive of patient response. Third, quantitative imaging in preclinical studies is most relevant in orthotopic locations. This is important for monitoring the growth and response of tumors in orthotopic locations using imaging modalities that are the same as those used in patients and may include radiography, ultrasound, MRI, CT and PET^6^.

While these O-PDXs mirror many of the human components, one limitation is the lack of a functional human immune system which restricts evaluation of immunotherapy in these models as presented. Methodologies for generating humanized mouse models are constantly improving^56,57^, yet there remains a paucity of immunocompetent pediatric solid tumor models. Approaches such as co-injecting human peripheral blood mononuclear cells or hematopoietic stem cells along with tumor cells have been utilized in some pediatric models^58^ and these methods can readily be applied in our collection. Additionally, we are currently generating 3D and 2D *in vitro* model systems from our O-PDXs^59^ where human immune components can be co-cultured for short-term experiments.

### Precision Oncology Using O-PDXs

Here, we performed 32 preclinical studies using 642 mice that received 48,714 doses of chemotherapy. 3 different approaches to precision oncology were evaluated. First, we performed a preclinical MATCH study to demonstrate the feasibility of using O-PDXs for establishing scientific justification and translational relevance of MATCH based therapy. This is particularly important as the next phase of the NCI-COG-Pediatric MATCH study is being developed to include combination chemotherapy. Second, we tested an O-PDX for a newly discovered genetic lesion in MAP3K8 in a rare group of pediatric melanoma patients that are not included in the NCI-COG Pediatric MATCH trial. There are not enough patients for a disease-focused clinical trial nor is the oncogenic mutation common enough to be included in a basket trial such as NCI-COG Pediatric MATCH. Third, we extended our preclinical precision medicine approach to include precision drug delivery using L-IRN. This formulation is easier to administer in pediatric patients, has lower toxicity due to reduced systemic exposure and retains tumor exposure over several days following a single dose. Efficacy was improved with this precision drug delivery approach. Taken together, these examples demonstrate the feasibility and highlight the value of incorporating preclinical testing for precision medicine in oncology using our freely distributed collection of credentialed O-PDXs. As future precision oncology trials are planned for childhood cancer, preclinical studies may represent an important first step in trial design and drug combination selection. We envision a seamless integration of discovery from preclinical studies to the clinic and back again. That is, evidence of efficacy for an NCI-COG-Pediatric MATCH treatment group could prompt expansion and/or refinement of the clinical trial cohort and exceptional responders in the clinical trial would encourage much more extensive preclinical testing. Through such an iterative process, efficacious MATCH based therapies could rapidly be prioritized for further investigation as was done for TRK inhibitors. Our preclinical precision medicine approach also provides a valuable framework for testing combination MATCH drugs. Our data on PARP inhibitors with IRN and TMZ demonstrate the value of such an approach. PARP inhibitors have no activity as single agents in preclinical studies or in patients but the combination of PARP inhibitors with IRN and TMZ was synthetic lethal in preclinical studies^6^, had activity in patients^44^ and is now being tested in a multi-center randomized phase II trial (NCT04901702).

The O-PDXs are also useful for testing precision medicine for diseases and/or oncogenic mutations that are ultra-rare and unlikely to be tested otherwise. By freely sharing our cohort of data, we seek to better understand variants of unknown significance and recognize that harmonizing datasets across pediatric solid tumors will benefit the scientific community. Importantly, our new sequence data presented here includes patient tumor, germline and O-PDX for most tumors making it a particularly useful resource for future meta-analyses. As genomic efforts expand and increasingly rare oncogenic drivers are identified, such O-PDXs that come from those patients are essential for assessing the scientific justification and translational relevance of novel therapeutic regimens.

The most effective precision medicine approach using our O-PDXs in this study came from precision drug delivery using L-IRN. This formulation was designed to minimize systemic exposure and toxicity while maintaining tumor penetration. Our preclinical PK and efficacy data confirmed that the liposomal formulation of IRN results in sustained SN-38 levels in the tumor while the systemic SN-38 was rapidly cleared. Our data are consistent with data from clinical trials showing that there are fewer side effects of L-IRN in adult oncology patients^50^. In addition to the pharmacologic benefits of L-IRN, we were surprised to discover that there was also greater efficacy in our Ewing sarcoma O-PDXs treated with L-IRN in combination with TAL. This is relevant and important as toxicity associated with IRN and TMZ was dose limiting in a recent clinical trial^51^. Therefore, the combination of L-IRN with TMZ and TAL is now being tested in a multi-center randomized phase II trial. In this example, the precision came from a new approach to drug delivery rather than a novel molecular target but it also highlights an important challenge in the field of precision medicine. As new molecular targeted therapeutics are incorporated into combination chemotherapeutic regimens, dose limiting toxicity may be a significant barrier in patients with recurrent/refractory cancer. Thus, a balanced approach to precision medicine that optimizes the targeted agent and delivery of conventional chemotherapy may benefit patients in the long term by increasing efficacy and reducing side effects of treatment.

The authors declare no potential conflicts of interest.

---

**Figure S1.**
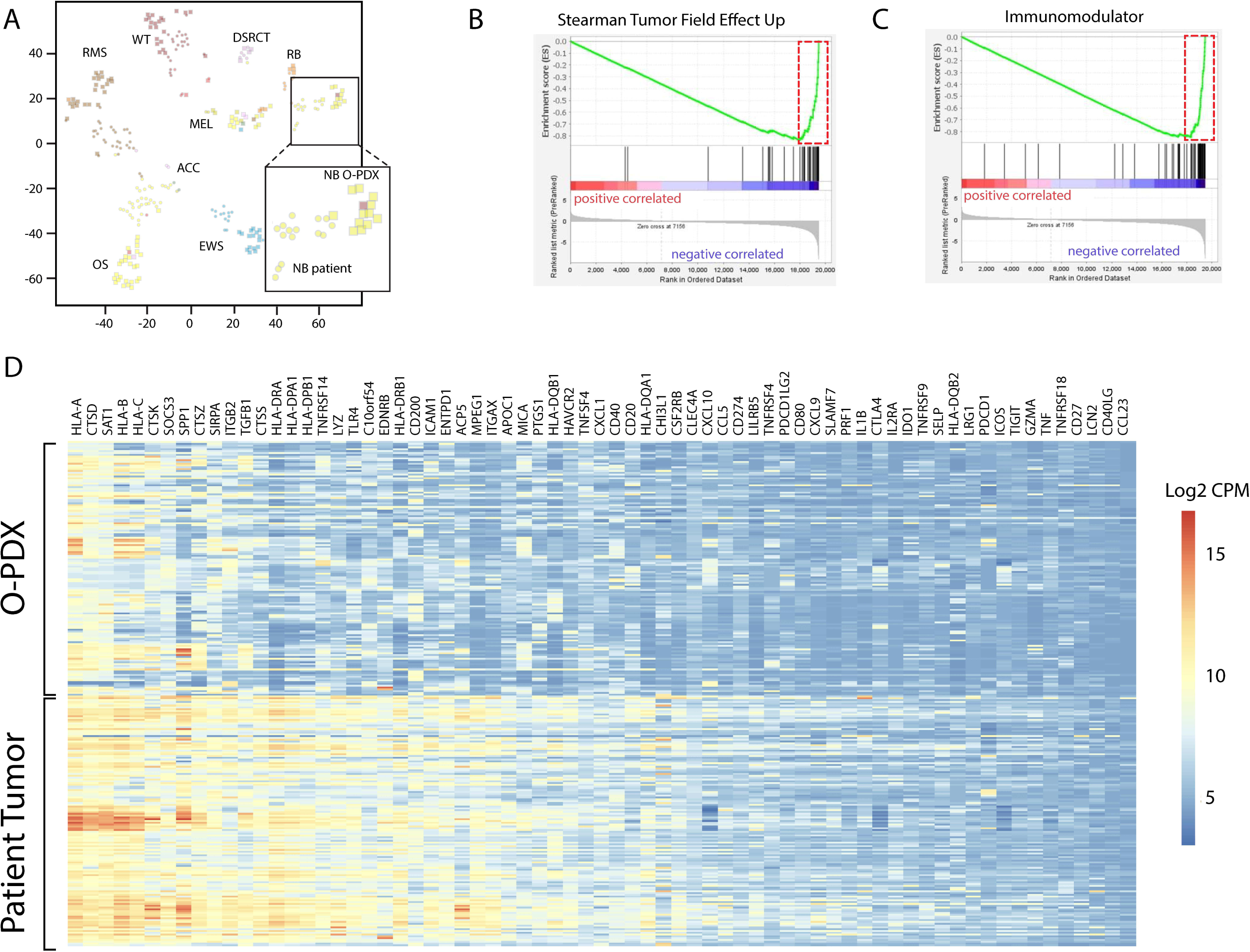
Human tumor microenvironment contributes to differences between patient tumor and O-PDXs. **A)** tSNE plot of RNA-seq showing separation of patient tumors (squares) and matched O-PDXs (circles). Regression and rank statistics (t-stat) was used to identify the genes that contribute to differences in gene expression between O-PDX and matched patient tumor. Those genes were then used for gene set enrichment analysis **(B, C)** with gene sets that represent cells in the tumor microenvironment. The leading edge genes (dashed red box) that had the strongest negative correlation with O-PDX (downregulated) were then used to generate the heatmap shown in **(D)**. Log2 fold change of counts per million reads are presented in the heatmap.

**Figure S2.**
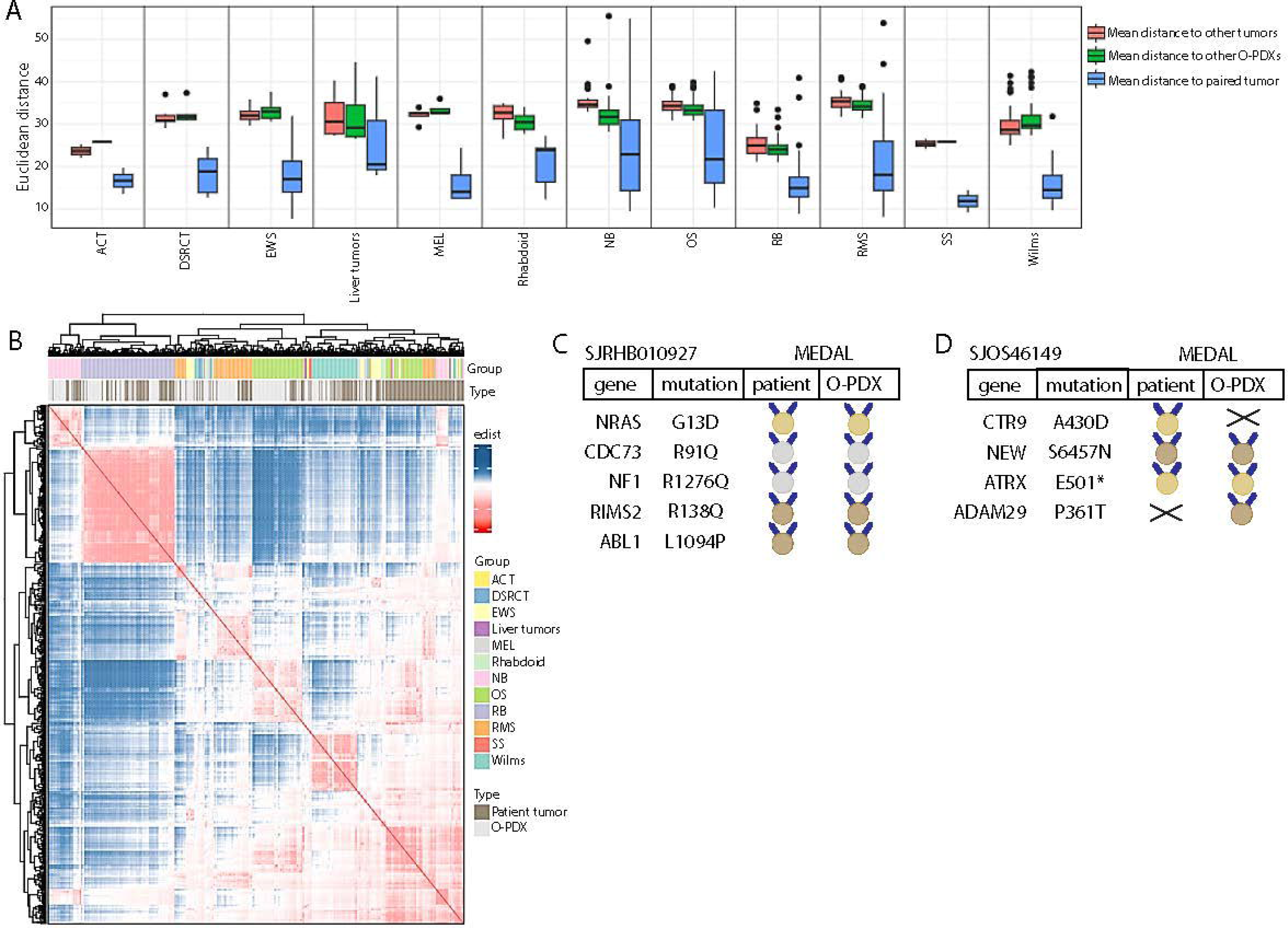
DNA methylation and clinical genomic characterization of matched patient tumor and O-PDXs. **A)** Box plot of calculated Euclidean distance from DNA methylation profiling showing O-PDXs are closer to their paired patient tumors (blue) compared to other patient tumors (red) or other O-PDXs (green) with the same noted tumor type. **B)** Heatmap of Euclidean distance calculated between patient and O-PDX tumors using all CpG probes showing clustering of samples with the same lineage. Tumor groups are color coded by histology and sample type is noted (patient tumors, dark brown; O-PDXs light brown). **C-D)** Medal ceremony results using the St. Jude Clinical Genomics pipeline showing representative patient and O-PDX comparisons. The SJRHB010927 (rhabdomyosarcoma) sample shown in (C) has preservation of all shown gold, silver, and bronze medals for the noted mutations in the O-PDX whereas the SJOS46149 (osteosarcoma) sample shown in (D) has partial preservation of medals with CTR9 and ADAM29 mutations lost and gained in the O-PDX respectively. Abbreviations: ACT, adrenal cortical tumor; DSRCT, desmoplastic small round cell tumor; EWS, Ewing sarcoma; MEL, melanoma; Rhabdoid, malignant rhabdoid tumor; NB, neuroblastoma; OS, osteosarcoma; RB, retinoblastoma; RMS, rhabdomyosarcoma; SS, synovial sarcoma; Wilms, Wilm’s tumor; edist, Euclidean distance.

**Figure S3.**
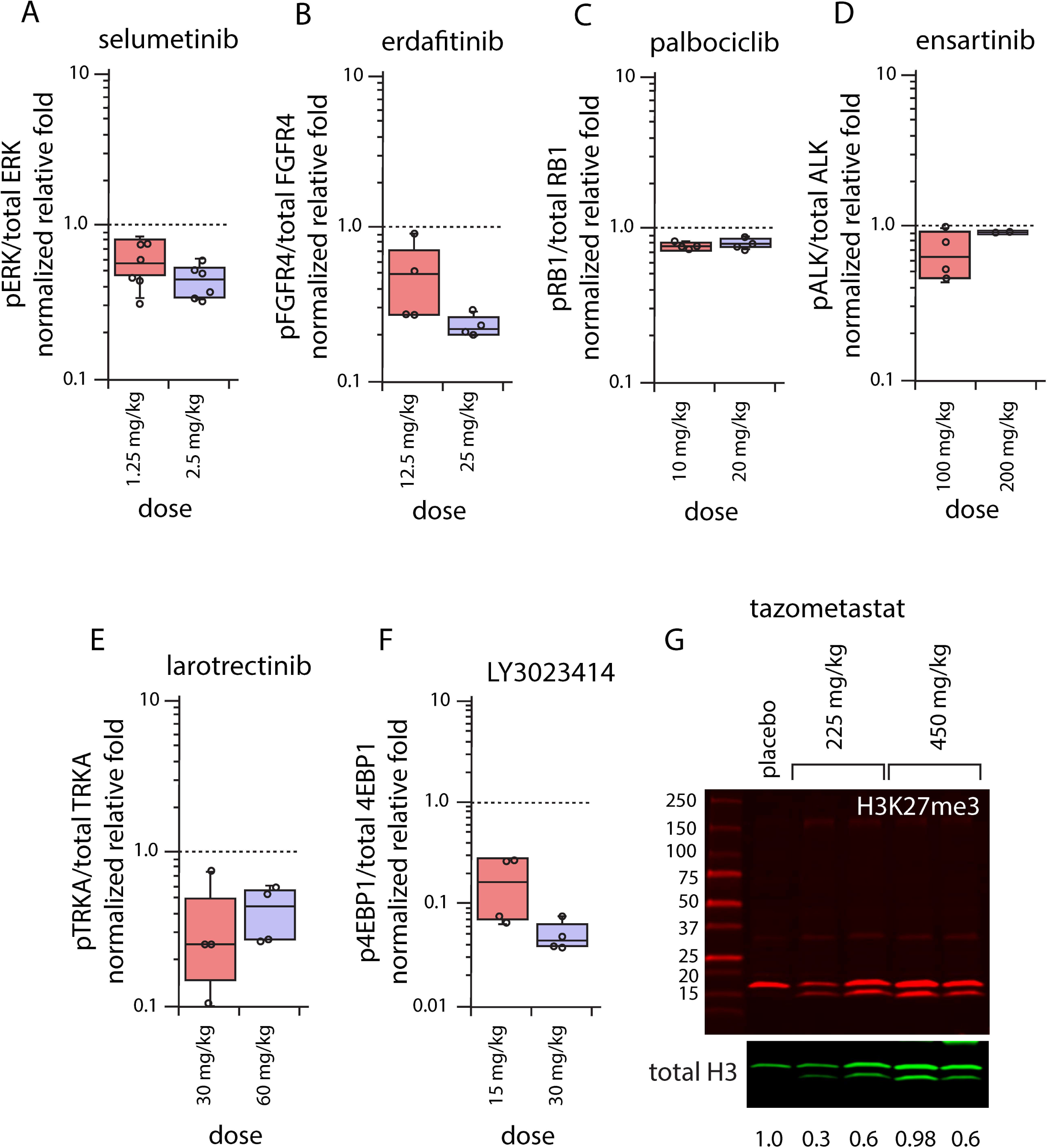
Pharmacodynamic studies of MATCH therapy. **A-F)** Box plots of phosphoprotein over total protein normalized relative fold for O-PDX tumors harvested from mice 4 hours after treatment with the noted therapy. Two doses were used for these PD experiments. Each datapoint is shown as a dot on top of the bars. **G)** Immunoblot of H3K27me3 and total H3 for samples treated with tazemetostat at indicated doses or placebo control. The relative reduction in the H3K27me3/total H3 levels are shown below each band.

**Figure S4.**
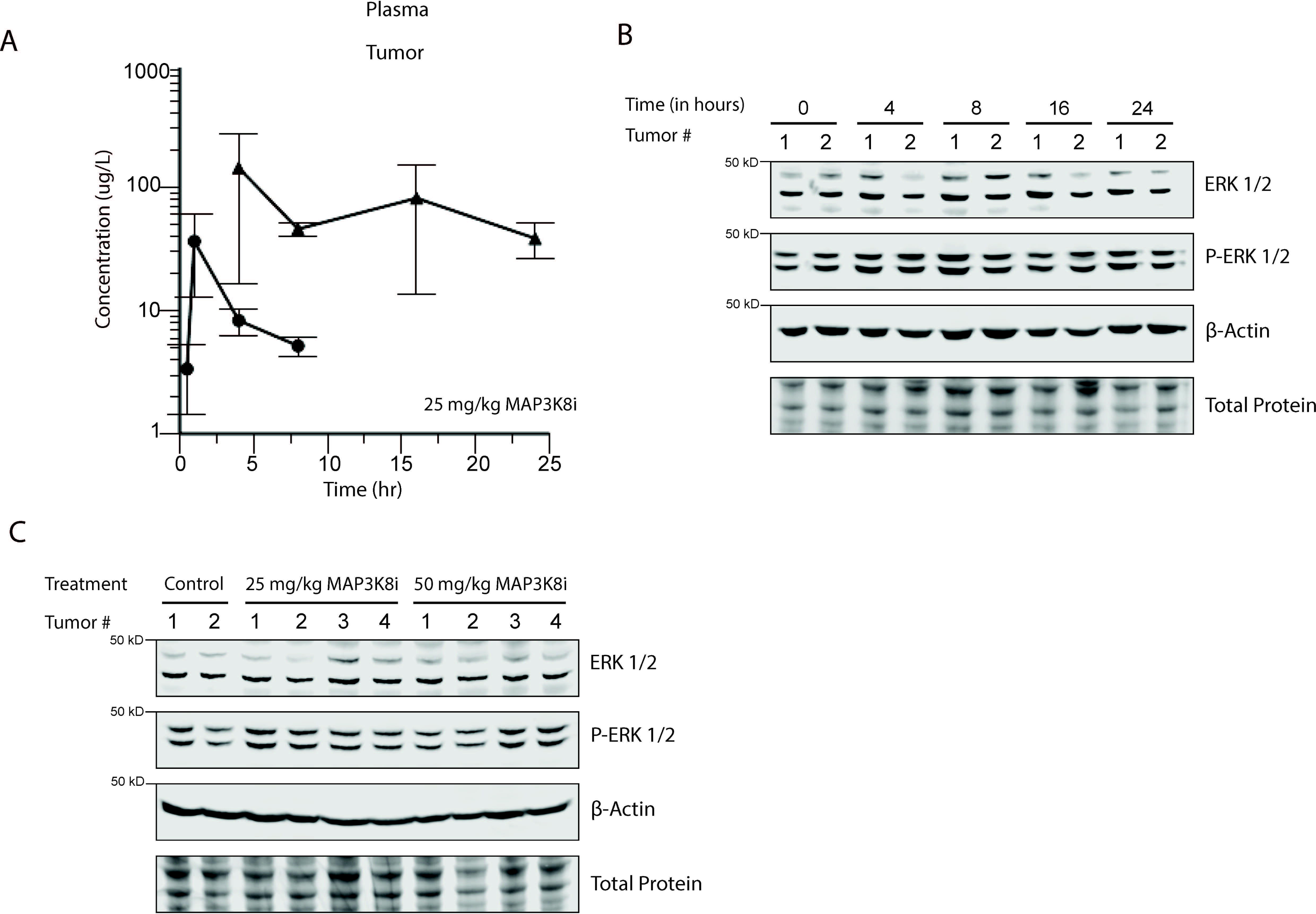
Pharmacokinetic and pharmacodynamic studies of MAP3K8 inhibitor. **A)** Concentration versus time plot of plasma (circle line) and tumor (triangle line) levels after a single dose (25 mg/kg) of a MAP3K8 inhibitor (MAP3Ki, PubChem:C26H25Cl2FN8). **B)** Western blot analysis of control or treated tumors from (A) at 0, 4, 8, 16, and 24 hours following treatment with MAP3K8i. **C)** Western blot analysis of control and treated tumors following treatment with 25 or 50 mg/kg MAP3K8i at 0, 4, 8, 16, and 24 hours post treatment showing no discernable change between groups. Abbreviations: P-ERK, phospho-ERK.

**Figure S5.**
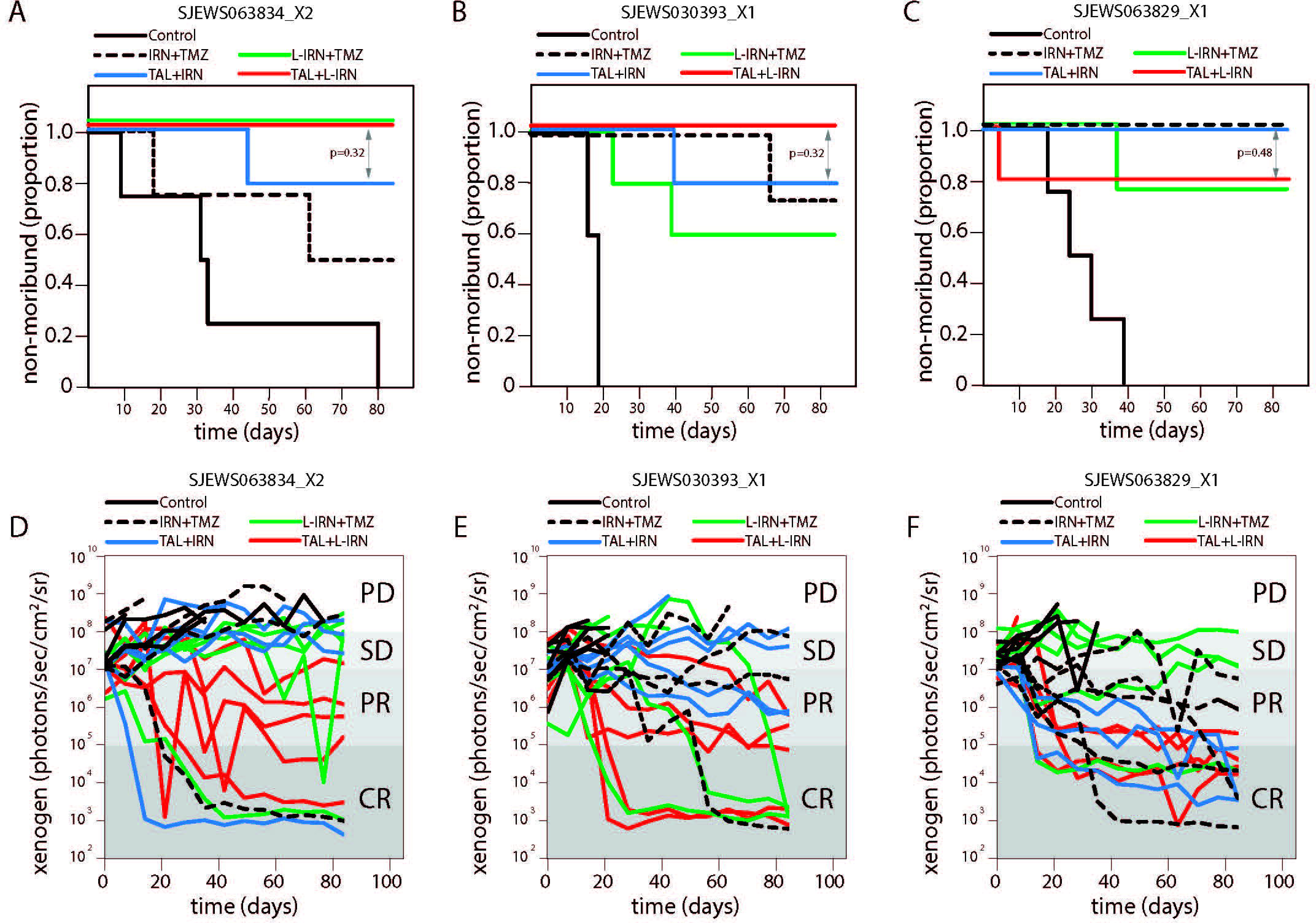
Preclinical studies using liposomal formulation of IRN in Ewing Sarcoma. **A-C)** Survival curves for standard and L-IRN in Ewing sarcoma O-PDXs (SJEWS063834_X2, SJEWS030393_X1, SJEWS063829_X1) treated with PARP inhibitor (talazoparib) combination chemotherapy. p values for each O-PDX comparing standard (blue) and liposomal IRN (red) with talazoparib are shown. **D-F)** Line graph of individual mouse response using xenogen for the O-PDXs from (A-C) for each of the treatment groups shown. Abbreviations: TAL, talazoparib; IRN, irinotecan; L-IRN, liposomal formulation of irinotecan; TMZ, temozolomide.

## ACKNOWLEDGEMENTS

This work was supported, in part, by Cancer Center Support (CA21765) from the NCI, grants to E.A.S. from National Comprehensive Cancer Network and St. Baldrick’s Foundation with generous support from the Invictus Fund, grants to M.A.D from the NIH (CA219686) and ALSAC. E.A.S. was also supported by the National Pediatric Cancer Foundation. M.A.D. was also supported by a grant from Alex’s Lemonade Stand Foundation for Childhood Cancer, the Tully Family Foundation, the Peterson Foundation, and by the Howard Hughes Medical Institute. The 3T3-NTRK cell line was received from Filemon Dela Cruz at Memorial Sloan Kettering Cancer Center. Preclinical imaging studies were performed with assistance from the Center for In Vivo Imaging and Therapeutics; electron microscopy was performed with assistance from the Cell and Tissue Imaging Shared Resource; pharmacokinetics was performed with assistance from the Preclinical Pharmacokinetics Shared Resource; and histopathologic analysis was performed with assistance from the Veterinary Pathology Shared Resource at St. Jude. Computational assistance was also provided by Honjian Jin and Patrick Schreiner from the Center for Applied Bioinformatics at St. Jude. Consultation for the use of MAP3K8 inhibitor was provided by Zoran Rankovich from Chemical Biology and Therapeutics. Liposomal irinotecan (Onivyde) was provided by Ipsen. BioVolume equipment was provided by Fuel3D with measurement assistance from Amanda George, Krista Millikin, and Chandra Savage (Research Services, St. Jude).

