## Supplementary material for "Preclinical Pediatric Molecular Analysis for Therapy Choice (MATCH)": Materials and Methods

### Patient Consent and MAST Protocol

Excess, deidentified tumor material was collected from solid tumor patients at St. Jude Children's Research Hospital in agreement with local institutional ethical regulations and institutional review board approval. Patient consent for tissue acquisition was obtained under the guidelines of the MAST protocol (NCT01050296).

### Animals

Athymic nude immunodeficient mice were purchased from Charles River (strain code 553). NSG mice were purchased from Jackson Laboratories (strain code 005557). This study was carried out in strict accordance with the recommendations in the Guide to Care and Use of Laboratory Animals of the National Institute of Health. The protocol was approved by the Institutional Animal Care and Use Committee at St. Jude Children's Research Hospital. All efforts were made to minimize suffering. All mice were housed in accordance with approved IACUC protocols. Animals were housed on a 12-12 light cycle (light on 6am off 6pm) and provided food and water *ad libitum*.

### Primary Patient Tissue Processing

Primary tumor tissue was processed for implantation within 2 hours of surgical resection in the majority of cases. Primary tumor was enzymatically dissociated into a single cell suspension and injected into an anatomically correct location for the disease type when possible. If the initial tumor sample was too small for dissociation, tumor tissue was implanted in the flank location. Initial implantation was primarily done into recipient NSG female mice with the exception of retinoblastoma which was implanted into nude mice. After engraftment and sufficient tumor growth, the tumor was harvested and passaged into nude mice using the same dissociation and implantation techniques.

### Orthotopic Injections

*Bone Marrow Injections*
To minimize distress and movement during the procedure, isoflurane gas anesthetic is used. The mouse is placed in the supine position on a nose cone prior to injection. The skin of the knee joint is prepped with alternating iodine scrubs and 70% isopropyl alcohol wipes. The prepped leg is flexed at the knee joint and secured to the work surface. Their femurs are palpated until the femoral condyles become visible**.** A 25gauge needle on a 50 µL Hamilton glass syringe (Hamilton Cat# 80920) is held at a 45 degree angle to the mouse, and the needle tip is inserted mid-femur via the femoral intercondylar notch, while retracting the patella and patella tendon medially to avoid ligament damage. The needle is advanced down to the femoral head, which is approximately 5-10 mm depending on the size of the mouse.

*Intramuscular injection*

The mouse is restrained gently but firmly by the scruff method. The rear foot nearest to the investigator is secured beneath the little finger and lower thumb. The area to be injected is swabbed with 70% ethanol. The needle is inserted, bevel up, into the caudal thigh at a 45 degree angle and cell suspension slowly injected into the muscle while avoiding injury to the sciatic nerve.

*Intravitreal injections*
Nude mice are given general anesthesia via isoflurane inhalation continuously at 1-3% concentration with an oxygen flow rate of 2 Liter/minute. The mouse is placed under the microscope where the eye is proptosed and a small incision using a 30 gauge needle is created between the sclera and the cornea. Using a 5 µL glass Hamilton syringe (Hamilton Company, 7633-01, 65RN) with a 33 gauge small hub needle, cells are injected into the vitreous of the eye.

*Ultrasound guided para-adrenal injections*
All ultrasound procedures are performed using the VEVO 2100 high frequency ultrasound equipped with a MS-550S transducer running at 40 MHz. Recipient immunocompromised mice are given general anesthesia via isoflurane inhalation continuously at 1-3% concentration with an oxygen flow rate of 2 Liter/minute. Anesthetized recipients are placed laterally on the imaging bed with left flank facing upward. In order to provide a channel for delivery of the implant, a 22 gauge catheter (BD Worldwide, Cat# 381423) is gently inserted through the skin and back muscle into the para-adrenal region and the hub is removed. A chilled 50 µLHamilton glass syringe (Hamilton Cat# 80920) fitted with a 27 gauge 1.5 inch needle is loaded with 10 µl of suspension and guided stereotactically through the catheter and positioned between the kidney and adrenal gland using ultrasound. The fluid is then injected into the region and the needle is left in place for 30 seconds to permit the matrigel component to set. The needle is then slowly removed, followed by gentle removal of the catheter. The mouse is placed in a clean cage on a warmer to recover from anesthesia.

**Flank Implantation**

Recipient immunocompromised mice are given general anesthesia via isoflorane inhalation continuously at 1-3% concentration with an oxygen flow rate of 2 Liter/minute. The mouse is placed ventral side down with a nose-cone to provide continued anesthesia. The area from mid-spine to the tail base is cleaned with 70% ethanol. A small horizontal 5mm incision in the flank area is made using sterile small surgical scissors. The tip of the sterile scissors is inserted into the incision, directly over the flank, and the scissors are opened to introduce a pocket in the subcutaneous space. One individual piece of tumor tissue is inserted into the pocket using sterile forceps. One drop of 100X penicillin/streptomycin solution is inserted into the opening over the tissue piece. The incision is closed with Vetbond tissue adhesive (3M Cat# 1469SB). The overlying skin is held together for 3-5 seconds with forceps to allow adequate time for drying.

**Enzymatic Tumor Dissociation for Initial Implantation or Passage**

*Neuroblastoma*
Tumor was minced with sterile scalpels and rinsed in phosphate buffered saline without calcium or magnesium (PBS-minus solution). Tumor suspension was transferred to a 50 mL conical tube and filled with PBS-minus solution. Dissociation was done by adding 600 µl of trypsin (10 mg/ml, Sigma Cat#T9935) and tube placed in 37 degree water bath for 10 minutes. Dissociation was stopped by

adding 60µL of Soybean Trypsin Inhibitor (10 mg/ml, Sigma Cat#T6522). Deoxyribonuclease I (2 mg/ml, Sigma Cat#D4513) and magnesium chloride (1 M) were added in equal amounts of 60 µL increments until tumor fragments easily settle at the bottom of the tube. Tumor suspension is filtered with a 40 micron cell strainer and then centrifuged at 500g (g=RCF) for 5 minutes. Supernatant was discarded and 10 ml of red blood cell lysis solution (5 Prime Cat#2301310) added and allowed to incubate at room temperature for 10 minutes. Solution of phosphate buffered saline without calcium or magnesium (PBS-minus, Lonza Cat#17-516F) /10% Fetal Bovine Serum (FBS, Biowest Cat#SO1520) was added to fill 50 ml conical tube and cell suspension centrifuged at 500g (g=RCF) for 5 minutes. Supernatant was discarded and cell pellet was resuspended in PBS-minus/10%FBS for counting. Cells were then resuspended in Matrigel™ basement membrane matrix (BD Biosicences cat. 354234) at a concentration of 2 x $10^5$ cells per 10 µL and placed on ice for injection.

*Soft tissue sarcoma (including RMS, high grade sarcoma, NR-STS, Ewing sarcoma)* Tumor was placed through a tumor press and then rinsed with Dulbecco's modified Eagle's medium (DMEM) (Lonza catalog no. 12- 604F). The tumor suspension was transferred to a 50 ml conical tube and filled with DMEM. Dissociation was done by adding 600 µL of trypsin (10 mg/ml; Sigma catalog no. T9935) and 50 mg of type II collagenase (275 U/mg; Worthington Biochemical catalog no. 4177), and then tube was placed in a 37C water bath for 1 hour. Dissociation was stopped by adding 600 µL of soybean trypsin inhibitor (10 mg/ml; Sigma catalog no. T6522). Deoxyribonuclease I (2 mg/ml; Sigma catalog no. D4513) and magnesium chloride (1 M) were added in equal amounts in 60 µL increments until tumor fragments easily settled at the bottom of the tube. The tumor suspension was filtered with a 40 micron cell strainer and centrifuged at 500g (g=RCF) for 5 min. The supernatant was discarded, and 10 ml of red blood cell lysis solution (5 PRIME catalog no. 2301310) was added and allowed to incubate at room temperature for 10 min. A solution of PBS without calcium or magnesium (PBS-minus; Lonza catalog no. 17-516F)/10% fetal bovine serum (FBS) (Biowest catalog no. SO1520) was added to fill a 50 ml conical tube, and the cell suspension was centrifuged at 500g for 5 min. The supernatant was discarded, and the cell pellet was resuspended in PBS- minus/10% FBS for counting. Cells were then resuspended in matrigel (BD Worldwide catalog no. 354234) at a concentration of 1 x $10^6$ per 100 µL and placed on ice for injection.

*Osteosarcoma*

The tumor was placed through a tumor press and then rinsed with PBS without calcium or magnesium (PBS-minus; Lonza catalog no. 17-516F). The tumor suspension was transferred to 100 ml screw cap glass bottle and filled with PBS-minus to the 100 ml mark. Dissociation was done by adding 600 µL of trypsin (10 mg/ml; Sigma catalog no. T9935) and 200 mg of type II collagenase (275 U/mg; Worthington Biochemical catalog no. 4177), and placed in a warm 37 degree water bath for 90 min, agitating with magnetic bead at ~200 rpm aiming to have all tissue circulating and lifted off the bottom of the glass bottle. Dissociation was stopped by adding 600 µL of soybean trypsin inhibitor (10 mg/ml; Sigma catalog no. T6522). Deoxyribonuclease I (2 mg/ml; Sigma catalog no. D4513) and magnesium chloride (1 M) were added in equal amounts in 60 µL increments until tumor fragments easily settled at the bottom of the tube. The tumor suspension was filtered with a 70 micron cell strainer and centrifuged at 500 g (g=RCF) for 5 min. The supernatant was discarded, and 10 ml of red blood cell lysis solution (5 PRIME catalog no. 2301310) was added and allowed to incubate at room temperature for 10 min. A solution of PBS-minus/10% fetal bovine serum (FBS) (Biowest catalog no. SO1520) was added to fill a 50 ml conical tube, and the cell suspension was centrifuged at 500 g for 5 min. The supernatant was discarded, and the cell pellet was resuspended in PBS- minus/10% FBS for counting. Cells were then resuspended in Matrigel™ basement membrane matrix (BD Biosciences cat. 354234) at a concentration of 1 x $10^6$ cells per 10 µL and placed on ice for injection.

*Retinoblastoma*

Tumor was minced with sterile scalpels and rinsed in RPMI (Lonza catalog no 12-167F). Tumor suspension was transferred to a 50 mL conical tube and filled with RPMI. Dissociation was done by adding 600 µL of trypsin (10 mg/ml, Sigma Cat#T9935) and tube placed in 37 degree water bath for 10 minutes. Dissociation was stopped by adding 600 µL of Soybean Trypsin Inhibitor (10 mg/ml, Sigma Cat#T6522). Deoxyribonuclease I (2 mg/ml, Sigma Cat#D4513) and magnesium chloride (1 M) were added in equal amounts of 60 µL increments until tumor fragments easily settle at the bottom of the tube. Tumor suspension is filtered with a 40 micron cell strainer and then centrifuged at 500g (G=RCF) for 5 minutes. Supernatant was discarded and 10ml of red blood cell lysis solution (5 Prime Cat#2301310) added and allowed to incubate at room temperature for 10 minutes. Solution of phosphate buffered saline without calcium or magnesium (PBS-minus, Lonza Cat#17-516F) /10% Fetal Bovine Serum (FBS, Biowest Cat#SO1520) was added to fill 50 ml conical tube and cell suspension centrifuged at 450g (G=RCF) for 5 minutes. Supernatant was discarded and cell pellet was resuspended in PBS-minus/10%FBS for counting. Cells were then resuspended in RPMI at a concentration of 1 x $10^5$ cells per 5 µL for injection.

**Tumor Cryopreservation**

After dissociation, tumor cells that were not utilized for passaging or high throughput screening were cryopreserved for banking and later usage. Cells were counted and resuspended in chilled FBS/10% DMSO at a concentration of 6 x $10^6$ cells per 1 ml per tube. Cryo tubes were placed in styrofoam containers and frozen at -$80^o$ C for 3 days and then transferred to liquid nitrogen for long term storage.

**Whole Genome Sequencing**

Whole genome sequencing and library construction was performed as described previously [1-2] with the following modifications; 250-500 ng of genomic DNA was input for library construction using Illumina compatible adapters, and 4-6 cycles of amplification was performed with Kapa HiFi Hotstart ReadyMix (KAPA Biosystems).

**Whole Exome Sequencing**

Whole exome sequencing was conducted using the SeqCap EZ HGSC VCRome (Roche) according to manufactures instructions.

**RNA-Sequencing**

RNA-Seq was performed using the TruSeq Stranded Total RNA Library Prep Kit (Illumina) with 250 ng-1 ug of total RNA as input.

**Transcriptome Sequencing Analysis**

The paired end sequencing reads were subjected to mouse read cleansing with "bbsplit" (https://sourceforge.net/projects/bbmap/) if the sample was derived from xenografts. The adapters in sequencing reads were trimmed with "trim_galore" (v0.4.4, https://www.bioinformatics.babraham.ac.uk/projects/trim_galore/, -q 20 –phred 33 -- paired). The trimmed sequencing reads were mapped with STAR [3] to human genome GRCh38. The expected gene counts calculated using RSEM [4] for each sample were compiled to one gene count matrix. Only genes annotated as level 1 or 2 by GENCODE (v31) were kept in the downstream analysis. In addition, only genes with count per million (CPM) more than 0.5 in at least one sample were kept. The normalization factor for each sample was calculated using "calcNormFactors" in the "edgeR" package (v3.26.8) [5], and gene expression values were transformed and normalized using voom [6] in the "limma" package (v3.40.6) [7] in R. For t-sne plot, the variance of count data was first stabilized using varianceStabilizingTransformation function in R DESeq2 package [8], the transformed data was then batch corrected using removeBatchEffect function in R limma package [7]. The batch effect was mainly due to different sequencing dates or protocols. Finally, the top 1000 most variable genes as defined by median absolute deviation were chosen to run tSNE with perplexity value of 10 ("Rtsne" package, version 0.15, https://github.com/jkrijthe/Rtsne).

**Methylation array analysis**

Genome-wide DNA methylation array was performed with Infinium MethylationEPIC BeadChip (Illumina, CA, USA) targeting 850,000 CpG sites in accordance with the manufacturer's instructions. Raw data files generated by the iScan array scanner were read and preprocessed by using "minfi" Bioconductor package. Unsupervised hierarchical clustering (Euclidean Distance and "Ward.D2" linkage) of DNA methylation profiling was performed based on the 5000 most variable methylation probes across all samples (or subset of samples), which were selected by variance of the beta values. For dimensionality reduction and visualization, PCA was performed in the initial steps using top 5000 most variable probes and first 50 dimensions were retained to run tSNE with perplexity values in the range (5-20) and 5000 iterations ("Rtsne" package, version 0.15, https://github.com/jkrijthe/Rtsne). Copy number variation (CNV) analysis from methylation array data was performed using the "Conumee" package (version 1.16.0). Most differentially methylated regions were detected with DMRcate (version 1.18.0, PMID: 25972926).

*Quantification of similarity between tumor-PDX pairs*

Raw methylation data from Illumina 850k arrays were processed using the Minfi package [24478339]. The SWAN method of normalization [22703947] was used to produced beta values, and the top 10,000 probes with the highest standard deviation were kept for downstream analysis. To generate UMAP plots, the UMAP package (https://github.com/tkonopka/umap) was used with default parameters. To explore how similar a given PDX sample is to its matched tumor, other tumors of the same lineage, or other xenografts of the same lineage, the Euclidean distance was calculated as a similarity metric. Within a given lineage, the mean Euclidean distance between all probes of a given PDX sample to all probes of its matched tumor sample, all probes of other tumors, and all probes of other xenografts was calculated. The means of these values across all PDXs for a given lineage were then used to summarize how similar that lineage's PDX samples were to the respective matched tumors, other tumors, and other xenografts.

**Mutation Detection of Normal-tumor Paired Samples**

The paired end sequencing reads were mapped with bwa [9]. The in-house somatic mutation detection procedure was described previously [10]. In addition, we also used an ensemble approach to call somatic mutations (SNV/indels) with multiple published tools, including Mutect2 (v4.1.2.0) [11], SomaticSniper (v1.0.5.0) [12], VarScan2 (v2.4.3) [13], MuSE (v1.0rc) [14] and Strelka2 (v2.9.10) [15]. The consensus calls by at least two callers were considered as confident mutations. The consensus call sets were further manually reviewed for the read depth, mapping quality, and strand bias to remove additional artifacts. The variant annotation was performed using Annovar [16]. Circos plots were generated using Circos (v0.69, PMID: 19541911). Somatic copy number alternations (SCNA) were determined by CONSERTING [17] and CNVkit (v0.9.6) [18]. For somatic structural variants, three SV callers were implemented in the workflow for SV calling, including Delly (v0.8.2) [19], Manta (v1.5.0) [20], and Gridss (v2.5.0)[21]. The SV calls passing the default quality filters of each caller were merged using SURVIVOR [22] and genotyped by SVtyper [23]. The intersected call sets were manually reviewed for the supporting soft-clipped and discordant read counts at both ends of a putative SV site.

**Mutation Detection of Tumor-only Samples**

For samples without matched normal samples, variants were determined by Mutect2 using the tumor-only calling mode. Sequencing artifacts were filtered following GATK best practice [24]. Multiple filtering steps were applied to exclude potential calling artifacts. The variants passing the filtering steps fulfilled the following criteria: coverage depth of the variant > 10, variant allele frequency >0.01, alternative allele count >=4, allele population frequency in public databases <0.01 (gnomadAD, 1000 genomes, ExAC and Exome Sequencing Projects), mappability > 0.7, not co-localized with repeat elements and GC percentage between 0.4 and 0.6. The same set of SV/SCNA callers were applied for identification of CNA and SV. The lesions detected in candidate genes were reviewed manually.

**Medal Ceremony**

To determine if the likely pathogenic somatic SNVs and indels are preserved in the O-PDX relative to the patient tumor, we used the St. Jude clinical genomics pipeline used in the genomes for kids (G4K) clinical trial (NCT02530658) as previously published [25] for a subset (n=81) of our patient/O-PDX pairs. The St. Jude Medal Ceremony clinical genomics pipeline) (https://university.stjude.cloud/docs/pecan/pecan-pie/) assigns gold, silver or bronze medals for coding and splice-related variants in disease predisposition genes.

**MATCH Drug Selection**

Eligibility criteria was reviewed for each MATCH drug according to the clinical protocol (NCT03155620). Patient and PDX samples were considered a “match” if the sample met the mutation criteria specified for each drug in the clinical protocol.

**In Vitro Screening of Xenografts**

Primary cultures of PDX models were plated for drug screening as previously described [2] and drug sensitivity and EC50 values for the models presented were obtained from the St. Jude PDX Drug

Screening Portal (https://braid.stjude.org/masttour/).

### Pharmacokinetics

*Ensartinib*

The first ensartinib PK study was a survival plasma PK evaluation using non-tumor bearing athymic nude mice. Ensartinib was suspended in 1% hydroxyethylcellulose (MW 720,000), 0.25% Tween 80, and ~0.05% simethicone at 2.5 mg/mL and administered as a 10 mL/kg oral gavage for a 25 mg/kg dose. A batch sampling design was implemented where 3 samples were collected per mouse. Mice were divided into 3 groups for sample collection. Mice from group 1 were sampled at 0.125, 1, and 16 hr post-dose. Mice from group 2 were sampled at 0.25, 2, and 24 hr, and mice from group 3 were sampled at 0.5, 4, and 8 hr post-dose. Blood samples (~ 50 µL) were collected by retroorbital eye bleed technique using Minivette POCT 50 µL capillary devices containing K3EDTA (Sarstedt AG, Germany). Terminal samples at the last time point were collected by cardiac puncture using a 1 mL syringe, and the blood placed in a Sarstedt Microvette K3EDTA 500 µL tube.

In the second ensartinib PK study, the plasma and tumor PK were evaluated after a single oral dose of the ensartinib 25 mg/kg suspension. Female athymic nude mice bearing neuroblastoma orthotopic xenografts in the adrenal capsule were sacrificed using an IACUC-approved method at 0.125, 1, 4, 8, 16 hr post-dose (3 mice per timepoint). Blood was collected by cardiac puncture, after which the carcass was perfused with PBS, the tumor extracted, rinsed, and placed in a microcentrifuge tube. In all instances, blood samples were immediately centrifuged to plasma. Plasma and tumor samples were temporarily placed on dry ice until transfer to a deep freezer, and samples were stored at - 80 °C until analysis. Additionally, ensartinib fraction unbound in mouse and human plasma (Fu,p,m and Fu,p,h), and patient derived rhabdomyosarcoma tumor homogenate (Fu,t) was determined using rapid equilibrium dialysis (RED, Pierce Biotechnology, ThermoFisher Scientific, Waltham, MA). Briefly, blank mouse plasma and tumor homogenates, diluted with PBS, were spiked with compounds in triplicate achieving final concentrations of 10 µM, placed in donor wells of RED apparatus, and permitted to equilibrate for 4-6 hours at 37 °C. Compounds were assayed in donor and receiver well samples using LC-MS, with the fraction unbound calculated as the ratio of concentration in receiver to donor adjusted for any dilution [26].

Frozen tumor samples were weighed in tared 15 mL Lysing Matrix D tubes (MP Biomedical, Santa Ana, CA) and diluted with a 5:1 volume of ultra-pure water. The tumor samples were then homogenized with a FastPrep-24 system (MP Biomedicals, Santa Ana, CA). The homogenization consisted of four 6.0 M/S vibratory cycles of 1 min each on the FastPrep-24 system. To prevent over-heating due to friction, samples were placed on wet ice for 5 min between each cycle. The homogenates were then stored at - 80°C until analysis.

Plasma and tumor samples were analyzed for ensartinib (Selleckchem, Lot # S823001, purity 99.4%) with a qualified liquid chromatography – tandem mass spectrometry (LC- MS/MS) assay. Plasma calibrators and quality controls were spiked with solutions, corrected for salt content, prepared in acetonitrile. Plasma and tumor homogenate samples, 25 µL each, were protein precipitated with 100 µL of 25 ng/mL tazemetostat (ADOOQ, Lot # L12712B002, purity 99.7%) in acetonitrile as an internal standard. A 2 µL aliquot of the extracted supernatant was injected onto a Shimadzu LC-20ADXR high performance liquid chromatography system via a LEAP CTC PAL autosampler.

The LC separation was performed using a Phenomenex Kinetex EVO (2.6 µm C18 100Å, 50 x 2.1 mm) column maintained at 50 °C with gradient elution at a flow rate of 0.5 mL/min. The binary mobile phase consisted of water-acetonitrile-200 mM ammonium acetate pH 6.0 (90:10:10 v/v/v) in reservoir A and acetonitrile-water-200 mM ammonium acetate pH 6.0 (90:10:10 v/v/v) in reservoir B. The initial mobile phase consisted of 10% B with a linear increase to 100% B in 4 min. The column was then rinsed for 1 min at 100% B and then equilibrated at the initial conditions for 2.0 min for a total run time of 7 min. Under these conditions, the analyte and IS eluted at 2.01 and 2.28 min, respectively. Analyte and IS were detected with tandem mass spectrometry using a SCIEX API 5500 Q-TRAP in the positive ESI mode with the following mass transitions were monitored: ensartinib 561.16 -> 257.10, and tazemetostat 573.30 -> 486.30.

The method qualification and bioanalytical runs all passed acceptance criteria for non- GLP assay performance. A linear model (1/X2 weighting) fit the calibrators across the 1 to 500 ng/mL range, with a correlation coefficient (R) of 0.9995. The lower limit of quantitation (LLOQ), defined as a peak area signal-to-noise ratio of 5 or greater verses a matrix blank with IS, was 1 ng/mL. Sample dilution integrity was confirmed. For the plasma matrix, the intra-run precision and accuracy was ≤ 4.51% CV and 94.0% to 99.9%, respectively.

The resultant ensartinib concentration-time (Ct) data were grouped by study, matrix, and time point, and manual imputation of data below the lower limit of quantitation (BLOQ) was as follows: IF at any time point ≥ 2/3rds of the Ct results were above the LLOQ, the BLOQ data were replaced with a value of ½ LLOQ, ELSE the entire time point's data were treated as missing.

Then, using Phoenix WinNonlin 6.4 (Certara USA, Inc., Princeton, NJ), Ct data summary statistics were generated, and the ensartinib arithmetic mean Ct data for 1) each study and matrix, and for 2) plasma as an aggregate across studies (Study = Aggregate), was subjected to noncompartmental pharmacokinetic analysis (NCA).

The extravascular (Model 202) was applied, and area under the Ct curve (AUC) values were estimated using the "linear up log down" trapezoidal rule. The terminal phase was defined as the three time points at the end of the Ct profile, and the elimination rate constant (Ke) was estimated using an unweighted loglinear regression of the terminal phase. The terminal elimination half-life (T1/2) was estimated as 0.693/Ke, and the AUC from time 0 to infinity (AUCinf) was estimated as the AUC to the last time point (AUClast) + predicted Clast/Ke.

Other NCA parameters estimated included the observed maximum concentration (Cmax), time of Cmax (Tmax), concentration at the last observed time point (Clast), time of Clast (Tlast), apparent oral clearance (CL/F = Dose/AUCinf), and apparent terminal volume of distribution (Vz/F). The apparent partition coefficient of ensartinib from the plasma to the tissue of interest (Kp,tissue) was estimated as the ratio of the AUCinf, tissue to AUCinf plasma when available.

To estimate a clinically relevant dosage (CRD) for mice, the resultant mouse plasma unbound AUCinf was compared with the reported human unbound plasma PK value at the recommended phase 2 dose (RP2D) of ensartinib of 225 mg PO QD [27]. All inferences were made under the assumption of timeindependent, linear and dose- proportional PK in mice and humans.

*Latrotrectinib*

The first larotrectinib PK study was a survival plasma PK evaluation using non-tumor bearing athymic nude mice. Larotrectinib was suspended in 1% hydroxyethylcellulose (MW 720,000), 0.25% Tween 80, and ~0.05% simethicone at 3 mg/mL and administered as a 10 mL/kg oral gavage for a 30 mg/kg dose. A batch sampling design was implemented where 3 samples were collected per mouse. Mice were divided into 3 groups for sample collection. Mice from group 1 were sampled at 0.125, 1, and 16 hr post-dose. Mice from group 2 were sampled at 0.25, 2, and 24 hr, and mice from group 3 were sampled at 0.5, 4, and 8 hr post-dose. Blood samples (~ 50 µL) were collected by retro-orbital eye bleed technique using Minivette POCT 50 µL capillary devices containing K3EDTA (Sarstedt AG, Germany). Terminal samples at the last time point were collected by cardiac puncture using a 1 mL syringe, and the blood placed in a Sarstedt Microvette K3EDTA 500 µL tube.

In the second larotrectinib PK study, the plasma and tumor PK were evaluated after a single oral dose of the larotrectinib 30 mg/kg suspension. Female athymic nude mice bearing rhabdomyosarcoma orthotopic xenografts in the quadriceps were sacrificed using an IACUC-approved method at 0.125, 1, 4, 8, 16 hr post-dose (3 mice per timepoint). Blood was collected by cardiac puncture, after which the carcass was perfused with PBS, the tumor extracted, rinsed, and placed in a microcentrifuge tube.

In all instances, blood samples were immediately centrifuged to plasma. Plasma and tumor samples were temporarily placed on dry ice until transfer to a deep freezer, and samples were stored at -80 °C until analysis. Additionally, larotrectinib fraction unbound in mouse and human plasma (Fu,p,m and Fu,p,h), and patient derived rhabdomyosarcoma tumor homogenate (Fu,t) was determined using rapid equilibrium dialysis (RED, Pierce Biotechnology, ThermoFisher Scientific, Waltham, MA). Briefly, blank mouse plasma and tumor homogenates, diluted with PBS, were spiked with compounds in triplicate achieving final concentrations of 10 µM, placed in donor wells of RED apparatus, and permitted to equilibrate for 4-6 hours at 37 °C. Compounds were assayed in donor and receiver well samples using LC-MS, with the fraction unbound calculated as the ratio of concentration in receiver to donor adjusted for any dilution [26]. These experiments were conducted fully by SJCRH Chemical Biology and Therapeutics (CBT) Analytical Technologies Center (ATC) personnel under the direction of Lei Yang.

Frozen tumor samples were weighed in tared 15 mL Lysing Matrix D tubes (MP Biomedical, Santa Ana, CA) and diluted with a 5:1 volume of ultra-pure water. The tumor samples were then homogenized with a FastPrep-24 system (MP Biomedicals, Santa Ana, CA). The homogenization consisted of four 6.0 M/S vibratory cycles of 1 min each on the FastPrep-24 system. To prevent over-heating due to friction, samples were placed on wet ice for 5 min between each cycle. The homogenates were then stored at - 80°C until analysis.

Plasma and tumor samples were analyzed for larotrectinib (Abmole, Lot # NA, purity 99.7%) with a qualified liquid chromatography – tandem mass spectrometry (LC- MS/MS) assay. Plasma calibrators and quality controls were spiked with solutions, corrected for salt content, prepared in acetonitrile. Plasma and tumor homogenate samples, 25 µL each, were protein precipitated with 100 µL of 100 ng/mL LY3023414 (ADOOQ, Lot # L16126B001, purity 96.6%) in acetonitrile as an internal standard. A 1.5 µL aliquot of the extracted supernatant was injected onto a Shimadzu LC-20ADXR high performance liquid chromatography system via a LEAP CTC PAL autosampler.

The LC separation was performed using a Phenomenex Kinetex EVO (2.6 µm C18 100Å, 50 x 2.1 mm) column maintained at 50 °C with gradient elution at a flow rate of 0.5 mL/min. The binary mobile phase consisted of water-acetonitrile-200 mM ammonium acetate pH 6.0 (90:10:10 v/v/v) in reservoir A and acetonitrile-water-200 mM ammonium acetate pH 6.0 (90:10:10 v/v/v) in reservoir B. The initial mobile phase consisted of 10% B with a linear increase to 100% B in 4 min. The column was then rinsed for 1 min at 100% B and then equilibrated at the initial conditions for 2.0 min for a total run time of 7 min. Under these conditions, the analyte and IS eluted at 1.58 and 1.40 min, respectively. Analyte and IS were detected with tandem mass spectrometry using a SCIEX API 5500 Q-TRAP in the positive ESI mode with the following mass transitions were monitored: larotrectinib 429.19 -> 342.00, and LY3023414 407.20 -> 319.10

The method qualification and bioanalytical runs all passed acceptance criteria for non- GLP assay performance. A linear model (1/X2 weighting) fit the calibrators across the 1 to 500 ng/mL range, with a correlation coefficient (R) of 0.9993. The lower limit of quantitation (LLOQ), defined as a peak area signal- to-noise ratio of 5 or greater verses a matrix blank with IS, was 1 ng/mL. Sample dilution integrity was confirmed. For the plasma matrix, the intra-run precision and accuracy was ≤ 6.23% CV and 95.3% to 97.6%, respectively.

The resultant larotrectinib concentration-time (Ct) data were grouped by study, matrix, and time point, and manual imputation of data below the lower limit of quantitation (BLOQ) was as follows: IF at any time point ≥ 2/3rds of the Ct results were above the LLOQ, the BLOQ data were replaced with a value of ½ LLOQ, ELSE the entire time point's data were treated as missing.

Then, using Phoenix WinNonlin 6.4 (Certara USA, Inc., Princeton, NJ), Ct data summary statistics were generated, and the larotrectinib arithmetic mean Ct data for 1) each study and matrix, and for 2) plasma as an aggregate across studies (Study = Aggregate), was subjected to noncompartmental pharmacokinetic analysis (NCA).

The extravascular (Model 202) was applied, and area under the Ct curve (AUC) values were estimated using the "linear up log down" trapezoidal rule. The terminal phase was defined as the three time points at the end of the Ct profile, and the elimination rate constant (Ke) was estimated using an unweighted loglinear regression of the terminal phase. The terminal elimination half-life (T1/2) was estimated as 0.693/Ke, and the AUC from time 0 to infinity (AUCinf) was estimated as the AUC to the last time point (AUClast) + predicted Clast/Ke.

Other NCA parameters estimated included the observed maximum concentration (Cmax), time of Cmax (Tmax), concentration at the last observed time point (Clast), time of Clast (Tlast), apparent oral clearance (CL/F = Dose/AUCinf), and apparent terminal volume of distribution (Vz/F). The apparent partition coefficient of larotrectinib from the plasma to the tissue of interest (Kp,tissue) was estimated as the ratio of the AUCinf, tissue to AUCinf plasma when available.

To estimate a clinically relevant dosage (CRD) for mice, the resultant mouse plasma unbound AUCinf was compared with the reported pediatric unbound plasma PK value at the recommended phase 2 dose (RP2D) of larotrectinib of 100 mg/m2 PO BID [28]. All inferences were made under the assumption of time independent, linear and dose- proportional PK in mice and humans.

*LY3023414*

The first LY3023414 PK study was a survival plasma PK evaluation using non-tumor bearing athymic nude mice. LY3023414 was suspended in 1% hydroxyethylcellulose (MW 720,000), 0.25% Tween 80, and ~0.05% simethicone at 1.5 mg/mL and administered as a 10 mL/kg oral gavage for a 15 mg/kg dose. A batch sampling design was implemented where 3 samples were collected per mouse. Mice were divided into 3 groups for sample collection. Mice from group 1 were sampled at 0.125, 1, and 16 hr post-dose. Mice from group 2 were sampled at 0.25, 2, and 24 hr, and mice from group 3 were sampled at 0.5, 4, and 8 hr post-dose. Blood samples (~ 50 µL) were collected by retro-orbital eye bleed technique using Minivette POCT 50 µL capillary devices containing K3EDTA (Sarstedt AG, Germany). Terminal samples at the last time point were collected by cardiac puncture using a 1 mL syringe, and the blood placed in a Sarstedt Microvette K3EDTA 500 µL tube.

In the second LY3023414 PK study, the plasma and tumor PK were evaluated after a single oral dose of the LY3023414 15 mg/kg suspension. Female athymic nude mice bearing rhabdomyosarcoma orthotopic xenografts in the quadriceps were sacrificed using an IACUC-approved method at 0.125, 1, 4, 8, 16 hr post-dose (3 mice per timepoint). Blood was collected by cardiac puncture, after which the carcass was perfused with PBS, the tumor extracted, rinsed, and placed in a microcentrifuge tube.

In all instances, blood samples were immediately centrifuged to plasma. Plasma and tumor samples were temporarily placed on dry ice until transfer to a deep freezer, and samples were stored at -80 °C until analysis.

Additionally, LY3023414 fraction unbound in mouse and human plasma (Fu,p,m and Fu,p,h), and patient derived rhabdomyosarcoma tumor homogenate (Fu,t) was determined using rapid equilibrium dialysis (RED, Pierce Biotechnology, ThermoFisher Scientific, Waltham, MA). Briefly, blank mouse plasma and tumor homogenates, diluted with PBS, were spiked with compounds in triplicate achieving final concentrations of 10 µM, placed in donor wells of RED apparatus, and permitted to equilibrate for 4-6 hours at 37 °C. Compounds were assayed in donor and receiver well samples using LC-MS, with the fraction unbound calculated as the ratio of concentration in receiver to donor adjusted for any dilution [26].

Frozen tumor samples were weighed in tared 15 mL Lysing Matrix D tubes (MP Biomedical, Santa Ana, CA) and diluted with a 5:1 volume of ultra-pure water. The tumor samples were then homogenized with a FastPrep-24 system (MP Biomedicals, Santa Ana, CA). The homogenization consisted of four 6.0 M/S vibratory cycles of 1 min each on the FastPrep-24 system. To prevent over-heating due to friction, samples were placed on wet ice for 5 min between each cycle. The homogenates were then stored at - 80°C until analysis.

Plasma and tumor samples were analyzed for LY3023414 (ADOOQ, Lot # L16126B001, purity 96.6%) with a qualified liquid chromatography – tandem mass spectrometry (LC- MS/MS) assay. Plasma calibrators and quality controls were spiked with solutions, corrected for salt content, prepared in acetonitrile. Plasma and tumor homogenate samples, 25 µL each, were protein precipitated with 100 µL of 100 ng/mL selumetinib (Abmole, Lot # NA, purity 100%) in acetonitrile as an internal standard. A 3 µL aliquot of the extracted supernatant was injected onto a Shimadzu LC-20ADXR high performance liquid chromatography system via a LEAP CTC PAL autosampler. The LC separation was performed using a Phenomenex Kinetex EVO (2.6 µm C18 100Å, 50 x 2.1 mm) column

maintained at 50 °C with gradient elution at a flow rate of 0.5 mL/min. The binary mobile phase consisted of water-acetonitrile-200 mM ammonium acetate pH 6.0 (90:10:10 v/v/v) in reservoir A and acetonitrile-water-200 mM ammonium acetate pH 6.0 (90:10:10 v/v/v) in reservoir B. The initial mobile phase consisted of 10% B with a linear increase to 100% B in 4 min. The column was then rinsed for 1 min at 100% B and then equilibrated at the initial conditions for 2.0 min for a total run time of 7 min. Under these conditions, the analyte and IS eluted at 1.37 and 1.65 min, respectively. Analyte and IS were detected with tandem mass spectrometry using a SCIEX API 5500 Q-TRAP in the positive ESI mode with the following mass transitions were monitored:
407.20 -> 319.10 for LY3023414 and 457.00 -> 395.00 for selumetinib.

The method qualification and bioanalytical runs all passed acceptance criteria for non- GLP assay performance. A linear model (1/X2 weighting) fit the calibrators across the 1 to 500 ng/mL range, with a correlation coefficient (R) of 0.9995. The lower limit of quantitation (LLOQ), defined as a peak area signal- to-noise ratio of 5 or greater verses a matrix blank with IS, was 1 ng/mL. Sample dilution integrity was confirmed. For the plasma matrix, the intra-run precision and accuracy was ≤ 4.18% CV and 96.8% to 104.0%, respectively.

In the SPPK study, to compare sampling techniques at termination, a retro-orbital bleed and cardiac puncture was obtained from each mouse. All observations were used in the summary and PK analyses.

The resultant LY3023414 concentration-time (Ct) data were grouped by study, matrix, and time point, and manual imputation of data below the lower limit of quantitation (BLOQ) was as follows: IF at any time point ≥ 2/3rds of the Ct results were above the LLOQ, the BLOQ data were replaced with a value of ½ LLOQ, ELSE the entire time point's data were treated as missing.

Then, using Phoenix WinNonlin 6.4 (Certara USA, Inc., Princeton, NJ), Ct data summary statistics were generated, and the LY3023414 arithmetic mean Ct data for 1) each study and matrix, and for 2) plasma as an aggregate across studies (Study = Aggregate), was subjected to noncompartmental pharmacokinetic analysis (NCA).

The extravascular (Model 202) was applied, and area under the Ct curve (AUC) values were estimated using the "linear up log down" trapezoidal rule. The terminal phase was defined as the three time points at the end of the Ct profile, and the elimination rate constant (Ke) was estimated using an unweighted loglinear regression of the terminal phase. The terminal elimination half-life (T1/2) was estimated as 0.693/Ke, and the AUC from time 0 to infinity (AUCinf) was estimated as the AUC to the last time point (AUClast) + predicted Clast/Ke.

Other NCA parameters estimated included the observed maximum concentration (Cmax), time of Cmax (Tmax), concentration at the last observed time point (Clast), time of Clast (Tlast), apparent oral clearance (CL/F = Dose/AUCinf), and apparent terminal volume of distribution (Vz/F). The apparent partition coefficient of LY3023414 from the plasma to the tissue of interest (Kp,tissue) was estimated as the ratio of the AUCinf, tissue to AUCinf plasma when available.

To estimate a clinically relevant dosage (CRD) for mice, the resultant mouse plasma unbound AUCinf was compared with the reported pediatric unbound plasma PK value at the recommended phase 2 dose (RP2D) of LY3023414 of 200 mg PO BID [29, 30]. All inferences were made under the assumption of timeindependent, linear and dose- proportional PK in mice and humans.

*Selumetinib*

The first selumetinib PK study was a survival plasma PK evaluation using non-tumor bearing athymic nude mice. Selumetinib sulfate was suspended in 1% hydroxyethylcellulose (MW 720,000), 0.25% Tween 80, and ~0.05% simethicone at 1 mg/mL free base equivalents and administered as a 10 mL/kg oral gavage for a 10 mg/kg dose. A batch sampling design was implemented where 3 samples were collected per mouse. Mice were divided into 3 groups for sample collection. Mice from group 1 were sampled at 0.125, 1, and 16 hr post-dose. Mice from group 2 were sampled at 0.25, 2, and 24 hr, and mice from group 3 were sampled at 0.5, 4, and 8 hr post-dose. Blood samples (~ 50 µL) were collected by retro-orbital eye bleed technique using Minivette POCT 50 µL capillary devices containing K3EDTA (Sarstedt AG, Germany). Terminal samples at the last time point were collected by cardiac puncture using a 1 mL syringe, and the blood placed in a Sarstedt Microvette K3EDTA 500 µL tube.

In the second selumetinib PK study, the plasma and tumor PK were evaluated after a single oral dose of the selumetinib sulfate 10 mg/kg suspension. Female athymic nude mice bearing rhabdomyosarcoma orthotopic xenografts in the quadriceps were sacrificed using an IACUC-approved method at 0.125, 1, 4, 8, 16 hr post-dose (3 mice per timepoint). Blood was collected by cardiac puncture, after which the carcass was perfused with PBS, the tumor extracted, rinsed, and placed in a microcentrifuge tube. In all instances, blood samples were immediately centrifuged to plasma. Plasma and tumor samples were temporarily placed on dry ice until transfer to a deep freezer, and samples were stored at -80 °C until analysis.

Additionally, selumetinib fraction unbound in mouse and human plasma (Fu,p,m and Fu,p,h), and patient derived rhabdomyosarcoma tumor homogenate (Fu,t) was determined using rapid equilibrium dialysis (RED, Pierce Biotechnology, ThermoFisher Scientific, Waltham, MA). Briefly, blank mouse plasma and tumor homogenates, diluted with PBS, were spiked with compounds in triplicate achieving final concentrations of 10 µM, placed in donor wells of RED apparatus, and permitted to equilibrate for 4-6 hours at 37 °C. Compounds were assayed in donor and receiver well samples using LC-MS, with the fraction unbound calculated as the ratio of concentration in receiver to donor adjusted for any dilution [26]. These experiments were conducted fully by SJCRH Chemical Biology and Therapeutics (CBT) Analytical Technologies Center (ATC) personnel under the direction of Lei Yang.

Frozen tumor samples were weighed in tared 15 mL Lysing Matrix D tubes (MP Biomedical, Santa Ana, CA) and diluted with a 5:1 volume of ultra-pure water. The tumor samples were then homogenized with a FastPrep-24 system (MP Biomedicals, Santa Ana, CA). The homogenization consisted of four 6.0 M/S vibratory cycles of 1 min each on the FastPrep-24 system. To prevent over-heating due to friction, samples were placed on wet ice for 5 min between each cycle. The homogenates were then stored at - 80°C until analysis.

Plasma and tumor samples were analyzed for selumetinib (sulfate salt, Abmole, Lot # NA, purity 100%) with a qualified liquid chromatography – tandem mass spectrometry (LC-MS/MS) assay. Plasma calibrators and quality controls were spiked with solutions, corrected for salt content, prepared in acetonitrile. Plasma and tumor homogenate samples, 25 µL each, were protein precipitated with 100 µL of 7.25 ng/mL LY3023414 (ADOOQ, Lot # L16126B001, purity 96.6%) in acetonitrile as an internal standard. A 3 µL aliquot of the extracted supernatant was injected onto a Shimadzu LC-

20ADXR high performance liquid chromatography system via a LEAP CTC PAL autosampler.

The LC separation was performed using a Phenomenex Kinetex EVO (2.6 µm C18 100Å, 50 x 2.1 mm) column maintained at 50 °C with gradient elution at a flow rate of 0.5 mL/min. The binary mobile phase consisted of water-acetonitrile-200 mM ammonium acetate pH 6.0 (90:10:10 v/v/v) in reservoir A and acetonitrile-water-200 mM ammonium acetate pH 6.0 (90:10:10 v/v/v) in reservoir B. The initial mobile phase consisted of 10% B with a linear increase to 100% B in 4 min. The column was then rinsed for 1 min at 100% B and then equilibrated at the initial conditions for 2.0 min for a total run time of 7 min. Under these conditions, the analyte and IS eluted at 1.64 and 1.36 min, respectively. Analyte and IS were detected with tandem mass spectrometry using a SCIEX API 5500 Q-TRAP in the positive ESI mode with the following mass transitions were monitored:
457.00 -> 395.00 for selumetinib and 407.20 -> 319.10 for LY3023414.

The method qualification and bioanalytical runs all passed acceptance criteria for non- GLP assay performance. A linear model (1/X2 weighting) fit the calibrators across the 1 to 500 ng/mL range, with a correlation coefficient (R) of 0.9993. The lower limit of quantitation (LLOQ), defined as a peak area signal- to-noise ratio of 5 or greater verses a matrix blank with IS, was 1 ng/mL. Sample dilution integrity was confirmed. For the plasma matrix, the intra-run precision and accuracy was ≤ 6.23% CV and 95.3% to 97.6%, respectively.

The resultant selumetinib concentration-time (Ct) data were grouped by study, matrix, and time point, and manual imputation of data below the lower limit of quantitation (BLOQ) was as follows: IF at any time point ≥ 2/3rds of the Ct results were above the LLOQ, the BLOQ data were replaced with a value of ½ LLOQ, ELSE the entire time point's data were treated as missing.

Then, using Phoenix WinNonlin 6.4 (Certara USA, Inc., Princeton, NJ), Ct data summary statistics were generated, and the selumetinib arithmetic mean Ct data for 1) each study and matrix, and for 2) plasma as an aggregate across studies (Study = Aggregate), was subjected to noncompartmental pharmacokinetic analysis (NCA).

The extravascular (Model 202) was applied, and area under the Ct curve (AUC) values were estimated using the "linear up log down" trapezoidal rule. The terminal phase was defined as the three time points at the end of the Ct profile, and the elimination rate constant (Ke) was estimated using an unweighted loglinear regression of the terminal phase. The terminal elimination half-life (T1/2) was estimated as 0.693/Ke, and the AUC from time 0 to infinity (AUCinf) was estimated as the AUC to the last time point (AUClast) + predicted Clast/Ke. Other NCA parameters estimated included the observed maximum concentration (Cmax), time of Cmax (Tmax), concentration at the last observed time point (Clast), time of Clast (Tlast), apparent oral clearance (CL/F = Dose/AUCinf), and apparent terminal volume of distribution (Vz/F). The apparent partition coefficient of selumetinib from the plasma to the tissue of interest (Kp,tissue) was estimated as the ratio of the AUCinf, tissue to AUCinf plasma when available.

To estimate a clinically relevant dosage (CRD) for mice, the resultant mouse plasma unbound AUCinf was compared with the reported pediatric unbound plasma PK value at the recommended phase 2 dose (RP2D) of selumetinib of 25 mg/m2 PO BID [31]. All inferences were made under the assumption of timeindependent, linear and dose- proportional PK in mice and humans.

*Erdafitinib*

The first erdafitinib PK study was a survival plasma PK evaluation using non-tumor bearing athymic nude mice. Erdafitinib 12.5 mg/kg was first dissolved in a small amount of DMSO and formulated in 20% hydroxy-propyl beta cyclodextrin (HPBCD) at 1.25 mg/mL for a 10 mL/kg gavage volume. A batch sampling design was implemented where 3 samples were collected per mouse. Mice were divided into 3 groups for sample collection. Mice from group 1 were sampled at 0.125, 1, and 16 hr post-dose. Mice from group 2 were sampled at 0.25, 2, and 24 hr, and mice from group 3 were sampled at 0.5, 4, and 8 hr post-dose. Blood samples (~ 50 µL) were collected by retro-orbital eye bleed technique using Minivette POCT 50 µL capillary devices containing K3EDTA (Sarstedt AG, Germany). Terminal samples at the last time point were collected by cardiac puncture using a 1 mL syringe, and the blood placed in a Sarstedt Microvette K3EDTA 500 µL tube.

In the second erdafitinib PK study, the plasma and tumor PK were evaluated using 12.5 mg/kg erdafitinib 20% HPBCD solution formulation. Female athymic nude mice bearing rhabdomyosarcoma orthotopic xenografts in the quadriceps were sacrificed using an IACUC-approved method at 0.125, 1, 4, 8, 16 hr post-dose (3 mice per timepoint). Blood was collected by cardiac puncture, after which the carcass was perfused with PBS, the tumor extracted, rinsed, and placed in a microcentrifuge tube.

The third erdafitinib PK study, was a repeat of the first plasma-only study, using 12.5 mg/kg erdafitinib suspended in 1% hydroxyethylcellulose (MW 720,000), 0.25% Tween 80, and ~0.05% simethicone, 1.25 mg/kg for a 10 mL/kg gavage.

In all instances, blood samples were immediately centrifuged to plasma. Plasma and tumor samples were temporarily placed on dry ice until transfer to a deep freezer, and samples were stored at -80 °C until analysis.

Additionally, erdafitinib fraction unbound in mouse and human plasma (Fu,p,m and Fu,p,h), and patient derived rhabdomyosarcoma tumor homogenate (Fu,t) was determined using rapid equilibrium dialysis (RED, Pierce Biotechnology, ThermoFisher Scientific, Waltham, MA). Briefly, blank mouse plasma and tumor homogenates, diluted with PBS, were spiked with compounds in triplicate achieving final concentrations of 10 µM, placed in donor wells of RED apparatus, and permitted to equilibrate for 4-6 hours at 37 °C. Compounds were assayed in donor and receiver well samples using LC-MS, with the fraction unbound calculated as the ratio of concentration in receiver to donor adjusted for any dilution [26]. These experiments were conducted fully by SJCRH Chemical Biology and Therapeutics (CBT) Analytical Technologies Center (ATC) personnel under the direction of Lei Yang.

Frozen tumor samples were weighed in tared 15 mL Lysing Matrix D tubes (MP Biomedical, Santa Ana, CA) and diluted with a 5:1 volume of ultra-pure water. The tumor samples were then homogenized with a FastPrep-24 system (MP Biomedicals, Santa Ana, CA). The homogenization consisted of four 6.0 M/S vibratory cycles of 1 min each on the FastPrep-24 system. To prevent over-heating due to friction, samples were placed on wet ice for 5 min between each cycle. The homogenates were then stored at - 80°C until analysis.

Plasma and tumor samples were analyzed for erdafitinib (MCE, Lot # 17782, purity 97.6%) with a qualified liquid chromatography – tandem mass spectrometry (LC- MS/MS) assay. Plasma calibrators and quality controls were spiked with solutions, corrected for salt content, prepared in acetonitrile.

Plasma and tumor homogenate samples, 25 µL each, were protein precipitated with 100 µL of 50 ng/mL AZD4547 (LC- Labs, Lot # L11075B002) in acetonitrile as an internal standard. A 1 µL aliquot of the extracted supernatant was injected onto a Shimadzu LC-20ADXR high performance liquid chromatography system via a LEAP CTC PAL autosampler.

The LC separation was performed using a Phenomenex Kinetex (2.6 µm C18 100Å, 50 x 2.1 mm) column maintained at 50 °C with gradient elution at a flow rate of 0.4 mL/min. The binary mobile phase consisted of water-acetonitrile (9:1 v/v) + 0.1% formic acid in reservoir A and methanol-formic acid (100:0.1 v/v)) in reservoir B. The initial mobile phase consisted of 35% B with a linear increase to 100% B in 1.25 min. The column was then rinsed for 1 min at 100% B and then equilibrated at the initial conditions for 2.0 min for a total run time of 4.25 min. Under these conditions, the analyte and IS eluted at 1.27 and 1.18 min, respectively. Analyte and IS were detected with tandem mass spectrometry using a SCIEX API 5500 QTRAP in the positive ESI mode with the following mass transitions were monitored: 447.25 -> 388.20 for erdafitinib and 464.27 -> 217.20 for AZD4547.

The method qualification and bioanalytical runs all passed acceptance criteria for non- GLP assay performance. A linear model (1/X2 weighting) fit the calibrators across the 1 to 500 ng/mL range, with a correlation coefficient (R) of 0.9938. The lower limit of quantitation (LLOQ), defined as a peak area signal- to-noise ratio of 5 or greater verses a matrix blank with IS, was 1 ng/mL. Sample dilution integrity was confirmed. For the plasma matrix, the intra-run precision and accuracy was ≤ 4.07% CV and 88.1% to 108%, respectively. The resultant erdafitinib concentration-time (Ct) data were grouped by study, matrix, and time point, and manual imputation of data below the lower limit of quantitation (BLOQ) was as follows: IF at any time point ≥ 2/3rds of the Ct results were above the LLOQ, the BLOQ data were replaced with a value of ½ LLOQ, ELSE the entire time point's data were treated as missing.

Then, using Phoenix WinNonlin 6.4 (Certara USA, Inc., Princeton, NJ), Ct data summary statistics were generated, and the erdafitinib arithmetic mean Ct data for 1) each study and matrix, and for 2) plasma as an aggregate across studies (Study = Aggregate), was subjected to noncompartmental pharmacokinetic analysis (NCA).

The extravascular (Model 202) was applied, and area under the Ct curve (AUC) values were estimated using the "linear up log down" trapezoidal rule. The terminal phase was defined as the three time points at the end of the Ct profile, and the elimination rate constant (Ke) was estimated using an unweighted loglinear regression of the terminal phase. The terminal elimination half-life (T1/2) was estimated as 0.693/Ke, and the AUC from time 0 to infinity (AUCinf) was estimated as the AUC to the last time point (AUClast) + predicted Clast/Ke.

Other NCA parameters estimated included the observed maximum concentration (Cmax), time of Cmax (Tmax), concentration at the last observed time point (Clast), time of Clast (Tlast), apparent oral clearance (CL/F = Dose/AUCinf), and apparent terminal volume of distribution (Vz/F). The apparent partition coefficient of erdafitinib from the plasma to the tissue of interest (Kp,tissue) was estimated as the ratio of the AUCinf, tissue to AUCinf plasma when available.

To estimate a clinically relevant dosage (CRD) for mice, the resultant mouse plasma unbound AUCinf was compared with the reported human unbound plasma PK value at the Phase 1 single agent maximum tolerated dose of erdafitinib 10 mg PO [32]. All inferences were made under the assumption of time-independent, linear and dose- proportional PK in mice and humans.

*Tazemetostat*

The first tazemetostat PK study was a survival plasma PK evaluation using non-tumor bearing athymic nude mice. Tazemetostat was suspended in 1% hydroxyethylcellulose (MW 720,000), 0.25% Tween 80, and ~0.05% simethicone at 20 mg/mL and administered as a 10 mL/kg oral gavage for a 200 mg/kg dose. A batch sampling design was implemented where 3 samples were collected per mouse. Mice were divided into 3 groups for sample collection. Mice from group 1 were sampled at 0.125, 1, and 16 hr post-dose. Mice from group 2 were sampled at 0.25, 2, and 24 hr, and mice from group 3 were sampled at 0.5, 4, and 8 hr post-dose. Blood samples (~ 50 µL) were collected by retro-orbital eye bleed technique using Minivette POCT 50 µL capillary devices containing K3EDTA (Sarstedt AG, Germany). Terminal samples at the last time point were collected by cardiac puncture using a 1 mL syringe, and the blood placed in a Sarstedt Microvette K3EDTA 500 µL tube.

In the second tazemetostat PK study, the plasma and tumor PK were evaluated after a single oral dose of the tazemetostat 200 mg/kg suspension. Female athymic nude mice bearing rhabdoid orthotopic xenografts in the quadriceps were sacrificed using an IACUCapproved method at 0.125, 1, 4, 8, 16 hr post-dose (3 mice per timepoint). Blood was collected by cardiac puncture, after which the carcass was perfused with PBS, the tumor extracted, rinsed, and placed in a microcentrifuge tube.

In all instances, blood samples were immediately centrifuged to plasma. Plasma and tumor samples were temporarily placed on dry ice until transfer to a deep freezer, and samples were stored at -80 °C until analysis.

Additionally, tazemetostat fraction unbound in mouse and human plasma (Fu,p,m and Fu,p,h), and patient derived rhabdomyosarcoma tumor homogenate (Fu,t) was determined using rapid equilibrium dialysis (RED, Pierce Biotechnology, ThermoFisher Scientific, Waltham, MA). Briefly, blank mouse plasma and tumor homogenates, diluted with PBS, were spiked with compounds in triplicate achieving final concentrations of 10 µM, placed in donor wells of RED apparatus, and permitted to equilibrate for 4-6 hours at 37 °C. Compounds were assayed in donor and receiver well samples using LC-MS, with the fraction unbound calculated as the ratio of concentration in receiver to donor adjusted for any dilution [26]. These experiments were conducted fully by SJCRH Chemical Biology and Therapeutics (CBT) Analytical Technologies Center (ATC) personnel under the direction of Lei Yang.

Frozen tumor samples were weighed in tared 15 mL Lysing Matrix D tubes (MP Biomedical, Santa Ana, CA) and diluted with a 5:1 volume of ultra-pure water. The tumor samples were then homogenized with a FastPrep-24 system (MP Biomedicals, Santa Ana, CA). The homogenization consisted of four 6.0 M/S vibratory cycles of 1 min each on the FastPrep-24 system. To prevent over-heating due to friction, samples were placed on wet ice for 5 min between each cycle. The homogenates were then stored at - 80°C until analysis.

Plasma and tumor samples were analyzed for tazemetostat (ADOOQ, Lot # L12712B002, purity 99.7%) with a qualified liquid chromatography – tandem mass spectrometry (LC- MS/MS) assay. Plasma calibrators and quality controls were spiked with solutions, corrected for salt content, prepared in acetonitrile. Plasma and tumor homogenate samples, 25 µL each, were protein precipitated with

100 µL of 100 ng/mL GSK126 (ADOOQ, Lot # B006) in acetonitrile as an internal standard. A 2 µL aliquot of the extracted supernatant was injected onto a Shimadzu LC-20ADXR high performance liquid chromatography system via a LEAP CTC PAL autosampler.

The LC separation was performed using a Waters XBridge BEH C18 XP (2.5 µm 75 x 2.1 mm) column maintained at 50 °C with gradient elution at a flow rate of 0.6 mL/min. The binary mobile phase consisted of water: 20 mM ammonium acetate; formic acid (950:50:0.25 v/v/v) in reservoir A and acetonitrile: 20 mM ammonium acetate; formic acid (950:50:0.25 v/v/v) in reservoir B. The initial mobile phase consisted of 30% B with a linear increase to 100% B in 1.25 min. The column was then rinsed for 1 min at 100% B and then equilibrated at the initial conditions for 2.0 min for a total run time of 4.25 min. Under these conditions, the analyte and IS eluted at 0.94 and 1.08 min, respectively. Analyte and IS were detected with tandem mass spectrometry using a SCIEX API 5500 Q-TRAP in the positive ESI mode with the following mass transitions were monitored:
573.30 -> 486.30 for tazemetostat and 527.30 -> 392.20 for GSK126.

The method qualification and bioanalytical runs all passed P-PKSR's acceptance criteria for non-GLP assay performance. A linear model (1/X2 weighting) fit the calibrators across the 1 to 500 ng/mL range, with a correlation coefficient (R) of 0.9985. The lower limit of quantitation (LLOQ), defined as a peak area signal- to-noise ratio of 5 or greater verses a matrix blank with IS, was 1 ng/mL. Sample dilution integrity was confirmed. For the plasma matrix, the intra-run precision and accuracy was ≤ 5.81% CV and 93.6% to 99.3%, respectively.

The resultant tazemetostat concentration-time (Ct) data were grouped by study, matrix, and time point, and manual imputation of data below the lower limit of quantitation (BLOQ) was as follows: IF at any time point ≥ 2/3rds of the Ct results were above the LLOQ, the BLOQ data were replaced with a value of ½ LLOQ, ELSE the entire time point's data were treated as missing.

Then, using Phoenix WinNonlin 6.4 (Certara USA, Inc., Princeton, NJ), Ct data summary statistics were generated, and the tazemetostat arithmetic mean Ct data for 1) each study and matrix, and for 2) plasma as an aggregate across studies (Study = Aggregate), was subjected to noncompartmental pharmacokinetic analysis (NCA).

The extravascular (Model 202) was applied, and area under the Ct curve (AUC) values were estimated using the "linear up log down" trapezoidal rule. The terminal phase was defined as the three time points at the end of the Ct profile, and the elimination rate constant (Ke) was estimated using an unweighted loglinear regression of the terminal phase. The terminal elimination half-life (T1/2) was estimated as 0.693/Ke, and the AUC from time 0 to infinity (AUCinf) was estimated as the AUC to the last time point (AUClast) + predicted Clast/Ke.

Other NCA parameters estimated included the observed maximum concentration (Cmax), time of Cmax (Tmax), concentration at the last observed time point (Clast), time of Clast (Tlast), apparent oral clearance (CL/F = Dose/AUCinf), and apparent terminal volume of distribution (Vz/F). The apparent partition coefficient of tazemetostat from the plasma to the tissue of interest (Kp,tissue) was estimated as the ratio of the AUCinf, tissue to AUCinf plasma when available.

To estimate a clinically relevant dosage (CRD) for mice, the resultant mouse plasma unbound AUCinf was compared with the reported pediatric unbound plasma PK value at the recommended phase 2 dose (RP2D) of tazemetostat of 1200 mg/m2 PO BID [33]. All inferences were made under the assumption of time-independent, linear and dose- proportional PK in mice and humans.

*Ulixertinib*

The plasma pharmacokinetic (PK) profile of ERK 1/2 inhibitor ulixertinib hydrochloride was evaluate in female athymic nude mice (Charles River), approximately 8-12 weeks in age. Ulixertinib hydrochloride salt (SJ001001493-2, CHEMIETEK, Lot CT-VRT752, purity 99.87%) was suspended in 0.5% Methylcellulose (type 400 cPs) / 0.5% Tween 80 in ultrapure water (0.5% MCT), at a concentration of 1 mg/mL free base equivalents as a 10 mL/kg oral gavage, for a 10 mg/kg oral dose. Two survival blood samples were obtained from each mouse via retro-orbital plexus using 70 uL glass microhematocrit capillary tubes (Fisherbrand, Cat 22362574), and a third final sample by cardiac puncture, all using KEDTA as the anticoagulant. Samples were obtained at various times up to 24 hours post-dose, immediately processed to plasma, and stored at -80 °C until analysis. Remaining dosing solution was submitted for verification of potency, and chemical and physical stability during the study period.

Plasma samples were analyzed for ulixertinib with a qualified LC-MS/MS assay. Plasma calibrators and quality controls were spiked with solutions, corrected for salt content, prepared in methanol. Plasma samples, 10 µL each, were protein precipitated with 25 µL of 20 ng/mL LY3214996 (MCE, Lot # 33707, purity 99.95%) in methanol as an internal standard. A 2 µL aliquot of the extracted supernatant was injected onto a Shimadzu LC- 20ADXR high performance liquid chromatography system via a Shimadzu SIL-20AC XR autosampler. The LC separation was performed using a Phenomenex Syergi Hydro- RP (4 µm, 30 mm x 2 mm) column maintained at 40 °C with gradient elution at a flow rate of 0.25 mL/min. The binary mobile phase consisted of 0.1% formic acid in water- acetonitrile (90:10 v/v) in reservoir A and 0.1% formic acid in acetonitrile in reservoir B. The initial mobile phase consisted of 0% B with a linear increase to 100% B in 2 min. The column was then rinsed for 1 min at 100% B and then equilibrated at the initial conditions for 2.5 min for a total run time of 5 min. Under these conditions, the analyte and IS eluted at 2.85 and 2.59 min, respectively. Analyte and IS were detected with tandem mass spectrometry using a SCIEX API 4000 in the positive ESI mode with the following mass were transitions monitored: ulixertinib 433.20 -> 262.10, and LY3214996 454.20 -> 367.30.

The method qualification and bioanalytical runs all passed P-PKSR's acceptance criteria for non-GLP assay performance. A linear model (1/X2 weighting) fit the calibrators across the 1 to 500 ng/mL range, with a correlation coefficient (R) of ≥ 0.9991. The lower limit of quantitation (LLOQ), defined as a peak area signal-to-noise ratio of 5 or greater verses a matrix blank with IS, was 1 ng/mL for plasma and brain homogenate. The intra-run precision and accuracy was ≤ 5.88% CV and 85.9% to 106%, respectively.

Ulixertinib plasma Ct data were grouped by nominal time point, and the mean Ct values were subjected to noncompartmental analysis (NCA) using Phoenix WinNonlin 8.1 (Certara USA, Inc., Princeton, NJ). The extravascular model was applied, was applied, and area under the Ct curve (AUC) values were estimated using the "linear up log down" method. The terminal phase was defined as at least three time points at the end of the Ct profile, and the elimination rate constant (Kel) was estimated using an unweighted log- linear regression of the terminal phase. The terminal elimination half-life (T1/2) was estimated as 0.693/Kel, and the AUC from time 0 to infinity (AUCinf) was estimated as the AUC to the last time point (AUClast) + Clast (predicted)/Kel. Other parameters estimated included observed maximum concentration (Cmax), time of Cmax (Tmax), concentration at the last observed time point (Clast), time of Clast (Tlast), apparent clearance (CL/F = Dose/AUCinf), and apparent terminal volume of distribution (Vz/F).

*Liposomal Irinotecan*

The plasma pharmacokinetic (PK) profile of total (liposome encapsulated and non- encapsulated) irinotecan (IRN) and its active metabolite SN-38 were evaluated in female athymic nude mice (Charles River), approximately 12 weeks in age, in a mix of non- tumor bearing mice and mice bearing ES-8 Ewing sarcoma orthotopic xenografts across three separate studies. Onivyde (nal-IRI, nanoliposomal irinotecan, USP) was diluted with 5% dextrose in normal saline, to yield a dose of 5 mg/kg. A small study of tumor- bearing mice at a lower dose of nal-IRI 2.5 mg/kg IV was also conducted. In select mice across the studies, survival retro-orbital bleeds using Sarstedt 50 uL KEDTA POCT devices were obtained under isoflurane anesthesia, whereas terminal blood samples were obtained under IP Avertin (tribromoethanol) anesthesia. Blood samples were collected upon KEDTA, obtained at various times up to 168 hours postd-dose, immediately processed to plasma, and stored on dry ice until transfer to -80°C where they remained until analysis. Following terminal bleeds, animals were perfused with PBS to flush blood from the vasculature. Tissues were then extracted, rinsed with PBS as necessary, and then placed in appropriately labeled microcentrifuge tubes on dry ice. Tissue samples were then transferred to a -80°C freezer as soon as possible.

Frozen tumor samples were weighed in tared 15 mL Lysing Matrix D (MP Biomedical, Santa Ana, CA) tubes and diluted with a 5:1 volume of ultra-pure water. The tumor samples were then homogenized with a FastPrep-24 system (MP Biomedicals, Santa Ana, CA). The homogenization consisted of four 6.0 M/S vibratory cycles of 1 min each on the FastPrep-24 system. To prevent over-heating due to friction, samples were placed on wet ice for 5 min between each cycle. The homogenates were then stored at - 80°C until analysis.

Plasma and tumor homogenate samples were analyzed for irinotecan (SJ000312345-15, MCE) and SN38 (SJ000311679-8, TCI America) with a qualified LC MS/MS assay. Plasma calibrators and quality controls were spiked with solutions prepared in DMSO. Plasma samples, 25 µL each, were stabilized against carboxylesterase activity by the addition of 5 µL of 200 mM zinc sulfate and then protein precipitated with 100 µL of 10 ng/mL camptothecin (Cayman Chemical Co., Batch 0515272-12) in methanol as an internal standard. A 5 µL aliquot of the extracted supernatant was injected onto a Shimadzu LC-20ADXR high performance liquid chromatography system via a LEAP CTC PAL autosampler. The LC separation was performed using a Phenomenex Kinetex EVO C18 (2.6 µm, 50 mm x 2.1 mm) column maintained at 50 °C with gradient elution at a flow rate of 0.6 mL/min. The binary mobile phase consisted of water-acetonitrile-200 mM ammonium acetate pH 6.0 (90:10:10 v/v) in reservoir A and acetonitrile-water-200 mM ammonium acetate pH 6.0 (90:10:10 v/v) in reservoir B. The initial mobile phase consisted of 15% B with a linear increase to 60% B in 2.0 min. The column was then rinsed for 1.0 min at 100% B and then equilibrated at the initial conditions for 2.0 min for a total run time of 5 min. Under these conditions, irinotecan, IS and SN-38 eluted at 1.22, 1.43 and 1.48 min, respectively.

Analyte and IS were detected with tandem mass spectrometry using a SCIEX 5500 QTRAP in the positive ESI mode and the following mass transitions were monitored: Irinotecan 587.30 → 167.30, camptothecin (349.10 → 305.20) and SN-38 (393.10 → 305.20). The method qualification and bioanalytical runs all passed acceptance criteria for non-GLP assay performance. A linear model (1/X^2 weighting) fit the SN-38 calibrators across the 1 to 500 ng/mL range, with a correlation coefficient (R) of ≥ 0.9980 and 0.9992 for plasma and tumor, respectively. A linear model (1/X^2 weighting) fit the irinotecan calibrators across the 1 to 500 ng/mL range, with a correlation coefficient (R) of ≥ 0.9978 and 0.9972 for plasma and tumor, respectively. The lower limit of quantitation (LLOQ), defined as a peak area signal-to-noise ratio of 5 or greater verses a matrix blank with IS, was 1 ng/mL for both matrices, with a functional LLOQ of 6 ng/mL for tumor considering dilution. Sample dilution integrity was confirmed. The intra-run precision and accuracy for SN-38 in plasma was ≤ 11.6% CV and 93.2% to 107%, respectively. The intra-run precision and accuracy for irinotecan in plasma was ≤ 10.2% CV and 90.5% to 107%, respectively. For the tumor homogenate matrix, the intra-run precision and accuracy for SN-38 was ≤ 8.94% CV and 92.3% to 105%, respectively. The intra-run precision and accuracy for irinotecan in tumor homogenate was ≤ 11.5% CV and 93.1% to 104%, respectively.

The irinotecan and SN-38 concentration-time (Ct) data were grouped by matrix and nominal time point, and manual imputation of data below the lower limit of quantitation (BLOQ) was as follows: IF at any time point ≥ 2/3rds of the Ct results were above the LLOQ, the BLOQ data were replaced with a value of ½ LLOQ, ELSE the entire time point's data were treated as missing. Ct summary statistics including the arithmetic mean and standard deviation were then generated. The mean (+SD) Ct profiles were then plotted by analyte and matrix.

An approximate clinically relevant dose (CRD) of liposomal irinotecan for mice was estimated from plasma PK and exposure of total IRN to be 5 mg/kg IV. The approximate CRD was estimated as the mouse dose achieving a predicted mean plasma average concentration (Cavg) of IRN similar to humans with the adult nal-IRI FDA-approved dose. Dose proportional, linear, and time-invariant PK across species was assumed; however, evidence suggests this may not the case with nal-IRI [34]. Human and mouse plasma protein binding were assumed to be similar. Additional considerations influenced the final recommended mouse dose, including mouse dosing regimens prevalent in the literature and the tolerability of the compound in mice.

*MAP3K8 inhibitor (Cot Inhibitor-2, Compound 34)*
The plasma, brain, and subcutaneous melanoma xenograft tumor (SJMEL030083_X1) pharmacokinetic (PK) profile of Cot Inhibitor-2 (compound 34 in [35]) was evaluated in female athymic nude mice (Charles River) approximately 8-12 weeks in age. Cot Inhibitor-2 (SJ000988527-2, MedChemExpress, CAT# HY-32018, LOT# 05181) was suspended in 0.5% methylcellulose (type 400 cPs) / 2% Tween 80 / 0.5% L-(+)-Lactic acid, at 5 mg/mL for a 25 mg/kg free base equivalent dose as a 5 mL/kg oral gavage (formulation performed by CBT ATC staff). Terminal samples were collected over a 24-hour post-dose period by cardiac puncture using a 1 mL syringe, and the blood placed in a Sarstedt Microvette K3EDTA 500 µL tube, and immediately processed to plasma. The carcasses were perfused with PBS, brains and tumors extracted, rinsed, and placed in microcentrifuge tubes. All samples were immediately stored on dry ice and transferred to -80 °C until analysis. Remaining formulation, stored at 4° C protected from light, was submitted for verification of potency, and chemical and physical stability during the study period.

Brain and tumor samples were weighed in 4.5 mL and 15 mL Lysing matrix D (MP Biomedicals, Santa Ana, CA), diluted with a 1:5 volume of ultrapure water, and homogenized using a FastPrep-24 system (MP Biomedicals, Santa Ana, CA) for five cycles of 1 min vibration at 6.5 M/S speed, with 5 min in ice bath between each cycle to prevent over-heating. The homogenates were then stored at -80 °C for 24 hours until analysis.

Plasma and tissue homogenate samples were analyzed for Cot Inhibitor-2 (SJ000988527-2, MedChemExpress, CAT# HY-32018, LOT# 05181) using a qualified liquid chromatography – tandem mass spectrometry (LC-MS/MS) assay. Plasma calibrators and quality controls were spiked with solutions, corrected for salt content and purity as necessary, prepared in dimethyl sulfoxide. Plasma samples, 25 µL each, were protein precipitated with 75 µL of 10 ng/mL Cot Inhibitor-1 (MedChemExpress, CAT# HY-32015, LOT# 01596) in methanol as an internal standard (IS). A 5 µL aliquot of the extracted supernatant was injected onto a Shimadzu LC-20ADXR high performance liquid chromatography system via a LEAP CTC PAL autosampler. The LC separation was performed using a Phenomenex Kinetex EVO C18 (2.6 µm, 50 mm x 2.1 mm) column maintained at 50°C with gradient elution at a flow rate of 0.6 mL/min. The binary mobile phase consisted of water-acetonitrile-0.2 M ammonium acetate pH 6.0 (90:10:10 v/v) in reservoir A and methanol-water-0.2 M ammonium acetate pH 6.0 (90:10:10 v/v) in reservoir B. The initial mobile phase consisted of 55% B with a linear increase to 100% B in 1.25 min followed by a 2.25-minute hold at 100% B. The column was then equilibrated at the initial conditions for 2.0 min for a total run time of 5.5 min. Under these conditions, the analyte and IS eluted at 0.91 and 1.29 min, respectively.

Analyte and IS were detected with tandem mass spectrometry using a SCIEX QTRAP 5500 in the positive ESI mode and the following mass transitions were monitored: Cot Inhibitor-2 539.1 -> 112.1, Cot Inhibitor-1 553.2 -> 126.1. The method qualification and bioanalytical runs all passed acceptance criteria for non-GLP assay performance. A linear model (1/X2 weighting) fit the calibrators across the 1 to 500 ng/mL range, with a correlation coefficient (R) of ≥ 0.9989. The lower limit of quantitation (LLOQ), defined as a peak area signal-to-noise ratio of 5 or greater verses a matrix blank with IS, was 1 ng/mL for plasma and 6 ng/mL for brain and tumor tissues. Brain was quantitated using a brain homogenate calibration curve, while tumor was quantitated upon the plasma curve, given the rarity and unavailability of this matrix. Overall sample dilution integrity was confirmed. A significant matrix effect of ion or signal suppression was detected in control EDTA plasma from untreated experimental female athymic nude mice. Therefore, the use of this specific plasma for the preparation of all calibrators, QC's and matrix blanks was required. The intra-run precision and accuracy was ≤ 4.87% CV and 92.6% to 109%, respectively.

Plasma, brain and tumor concentration-time (Ct) data for Cot Inhibitor-2 were grouped by matrix and nominal time point. Manual imputation of data below the lower limit of quantitation (BLOQ) was as follows: if at any time point ≥ 2/3rds of the Ct results were above the LLOQ, the BLOQ data were replaced with a value of ½ LLOQ, or else the entire time point's data were treated as missing. Summary statistics were calculated, and the arithmetic mean Ct values were subjected to noncompartmental analysis (NCA) using Phoenix WinNonlin 8.1 (Certara USA, Inc., Princeton, NJ). The extravascular model was applied, and area under the Ct curve (AUC) values were estimated using the "linear up log down" method. The terminal phase was defined as at least three time points at the end of the Ct profile, and the elimination rate constant (Kel) was estimated using an unweighted log-linear regression of the terminal phase. The terminal elimination half-life (T1/2) was estimated as 0.693/Kel, and the AUC from time 0 to infinity (AUCinf) was estimated as the AUC to the last time point (AUClast) + Clast (predicted)/Kel. Additionally, an AUC from 0 to 8 hours (AUC0_8hr) was also estimated. Other parameters estimated included observed maximum concentration (Cmax), time of Cmax (Tmax), concentration at the last observed time point (Clast), time of Clast (Tlast), apparent clearance (CL/F = Dose/AUCinf), and apparent terminal volume of distribution (Vz/F), The apparent plasma-to-tissue partition coefficient (Kp0_8hrs) was estimated as the ratio of AUC0-8hr in tissue to AUC0-8hr in plasma. Finally, the average total concentration in the tumor (Cavg) was calculated as the tumor AUC from 0 to 24 hours divided by 24 hours.

Cot Inhibitor-2 fractions unbound in mouse plasma (Fu,p,m), and in mouse brain homogenate (Fu,b) were determined using rapid equilibrium dialysis (RED, 8K MWCO, Pierce Biotechnology, ThermoFisher Scientific, Waltham, MA). Briefly, blank plasma and brain homogenates were spiked with compounds and diluted with 100 mM PBS (6 replicates per group), achieving back-calculated final concentrations in blank plasma and brain tissue of 5 μM. The diluted samples were placed in donor wells of RED apparatus, and permitted to equilibrate for 6 hours (plasma) and 4 hours (brain) at 37 °C with 5% CO2. Compounds concentrations were assayed in donor and receiver well samples using the qualified LC-MS/MS assay, with the fraction unbound calculated as the ratio of concentration in receiver to donor, adjusted for any dilution [26]

Cot Inhibitor-2 Ct data demonstrated high variability between mice. Coefficients of variation ranged from 12.0% to 83.5%, with higher variability seen in tumor. All Cot Inhibitor-2 brain concentrations were BLOQ. The absorption rate of Cot Inhibitor-2 was rapid, with the Tmax occurring at 1 hour post-dose. After Cmax, plasma concentrations diminished in a nearly bi-exponential manner, with all plasma observations beyond 8 hours being BLOQ. The apparent terminal PK parameters (i.e. T1/2, CL/F, Vz/F) could not be estimated secondary to insufficient quantifiable data in the terminal phase. The formulation met specification (5.12 ± 0.371 mg/mL) and was stable over the 6-day usage period. The plasma protein binding of Cot Inhibitor-2 was high in mice, with an estimated fraction unbound of 0.00175 ± 0.000255. Brain homogenate binding appeared to be even higher than plasma. However, Cot Inhibitor-2 was unstable in brain tissue homogenate, precluding a reliable binding estimate, but suggesting possible extra-hepatic metabolism.

While plasma exposure was low, tumor penetration appeared to be slow (Tmax of 4 hours), relatively extensive (Kp0_8hrs of 6.66) and prolonged (T1/2 > 60 hours). The total tumor Cmax of 144 ng/mL and the Cavg of 26.3 ng/mL both fall short of the reported Tpl2 IC50 in human whole blood of ~162 ng/mL (300 nM) [35]. Moreover, assuming free fraction in tumor is similar to plasma, the unbound concentrations also fall well below the acellular IC50 for Tpl2 of ~0.863 ng/mL (1.6 nM) [35]. Thus, it is likely that insufficient tumor exposure is achieved under these studied conditions to sufficiently inhibit the target. In summary, the plasma exposure of Cot Inhibitor-2 in mice was low following a single oral dose of 25 mg/kg as a suspension. This low exposure could be due to an insufficiently enabled formulation and/or high systemic clearance of compound. Cot Inhibitor-2 does not appear to appreciably penetrate into the CNS. While tumor penetration was observed, it may not have been sufficient to adequately inhibit Cot / MAP3K8 / Tpl2 target.

*Olaparib, Trametinib, Palbociclib*

Pharmacokinetics for Olaparib, Trametinib, and Palbociclib were previously performed and the mouse equivalent dose calculated from those studies was used in preclinical studies here [2, 34]. The extended data reports for all preclinical PK studies that were performed can be found on the St. Jude Cloud (https://cstn.stjude.cloud/resources).

**Pharmacodynamics**

For PD study, mice were treated with two different doses (single or double) of each drug and perfused with PBS, 4 hours after dosage. Tumors were harvested and rinsed with PBS and kept fresh-frozen until use. The effects of drug treatments were determined by commercially available and validated direct/indirect sandwich ELISA assay kits for both total and phosphor proteins of Erk, TrkA, GFGR4, RB, 4EBP1, and ALK. These kits are ERK 1/2 (Total/Phospho) InstantOne ELISA assay Kit (85-86013; Invitrogen), Human Total TrkA: DuoSet IC ELISA kit (DYC175-2; R&D Systems), Human Phospho TrkA: DuoSet IC ELISA kit (DYC2578-2; R&D Systems), Human Total FGFR4: DuoSet IC ELISA kit (DYC685-2; R&D Systems), Human Phospho FGFR4: DuoSet IC ELISA kit (DYC5516-2; R&D Systems), FastScan Total Rb (23595; Cell Signaling), FastScan Phospho Rb (10754; Cell Signaling) and Human/Mouse Phospho-4E-BP1(T36) & Total 4E-BP1 ELISA (PEL-4EBP1-T36-T-1; RayBiotech), Total ALK and P-ALK (Y1604): PathScan Sandwich ELISA Kit (CST 7322CA and 7324CA, respectively). For the assessment of tazemetostat, immunoblotting against H3K27me3 and qRT-PCR for EZH2 were performed. Drugs, dosages and target proteins are listed below:

| Drug | Dose | Target |
|---|---|---|
| Selumetinib | 1.25 mg/kg | pErk |
| | 2.5 mg/kg | |
| Ensartinib | 100 mg/kg | pALK |
| | 200 mg/kg | |
| Larotrectinib | 30 mg/kg | pTrkA |
| | 60 mg/kg | |
| Erdafitinib | 12.5 mg/kg | pFGFR4 |
| | 25 mg/kg | |
| Palbociclib | 10 mg/kg | pRB |
| | 20 mg/kg | |
| LY3023414 | 15 mg/kg | p4EBP1 |
| | 30 mg/kg | |
| Tazemetostat | 225 mg/kg | EZH2 RT qPCR |
| | 450 mg/kg | H2K27Me3 WB |

The ELISA assay procedures were followed by the manufacturer's instructions. In brief, tumor tissue extracts, drugged or undrugged, were prepared with ice-cold 1X cell extraction buffer supplemented with protease and phosphatase inhibitors, with homogenization and sonication on ice and spun down, transferred cleared supernatant to new tubes and stored at -80 degree in single use aliquot. Total protein concentration was determined by BCA assay. Multiple dilutions of sample were made during the initial validation stages and in the tests to get a broad detection range. The Controls, internal controls provided in the kits and cell lysates positive controls prepared in the lab, and samples were assayed in duplicate or triplicate. PHERAstar microplate reader was used to measure the optical density (OD) at 450 nm after adding reaction stop solution. Most ELISA kits we used are for semi quantitative assays to measure relative OD changes between control and samples with treatment. Therefore, we analyzed the data by comparing the relative percentage changes of OD of phospho-protein/OD of total protein. Relative decrease was calculated by [OD (undrugged)-OD (drugged)]/ OD (undrugged).

*Western blot analysis MAP3K8 Inhibitor*
Athymic nude mice bearing melanoma tumors were treated with a single dose of MAP3K8 inhibitor (25 mg/kg and 50 mg/kg) and harvested after 4 hours. 10-15 mg of tumor were homogenized using a pestle in CelLytic MT Cell Lysis Reagent (SigmaC3228-50mL) supplemented with protease and phosphatase inhibitors (Thermo #A32959) and incubated on ice for 30 minutest to ensure complete lysis. Lysates were cleared by centrifugation at >17,000 x g for 10 minutes at 4°C. Following clearing, protein concentration was quantified using Pierce BCA Protein Assay (REF 23225). Samples were normalized to a concentration of 50 μg/sample in laemmli sample buffer and boiled for 10 minutes at 95°C. Samples were then run on a NuPAGE 4-12% Bis-Tris mini Protein gel (NP0322BOX) and transferred onto 0.2 μm Nitrocellulose blotting membrane (Amersham 10600011) for 2 hours at 250 mA using the BioRad Mini Trans-Blot system (Biorad #1703930). Following transfer, total protein staining was performed using the Revert 520 Total Protein Stain Kit (Licor 925-10016). Membranes were then blocked for 1 hour at room temperature in 5% BSA in TBS. Primary antibodies were diluted 1:1000 in 5% BSA in TBS and incubated on the membranes with gentle shaking overnight at 4°C. Antibodies used included ERK 1/2 (CST 9107S), and P-ERK 1/2 (T202/Y204) (CST 4370S) and β-actin (CST 3700S). Following primary antibody incubation, membranes were washed 3 times for 5 minutes in TBS with gentle shaking. Secondary antibodies (Licor #926-68071 and 926-32210) were diluted 1:10,000 in 5% non-fat milk and incubated on the membranes with gentle shaking for 1 hour at room temperature. Following secondary antibody incubation, the membranes were washed 3 times for 5 minutes in TBS with gentle shaking. Membranes were imaged using the Licor ClX and processed and analyzed using Image Studio.

**BaF3**

BaF3 cell lines that ectopically express druggable oncogenic fusions were generated as previously described [36]. The $BaF3^{ALK}$ line has an *EML4-ALK* fusion and the $BaF3^{NTRK}$ line has an *ETV6-NTRK3* fusion. Naïve BaF3 lines were supplemented with IL-3 (10μl/50mL) in RPMI and supplementation was stopped for lines that had oncogenic fusions.

*Lentiviral infection*
The BaF3 naïve, $BaF3^{ALK}$, and $BaF3^{NTRK}$ lines were plated at 300,000 cells per 2ml media (supplemented with polybrene and synperonic F108) in a 12-well dish, infected with up to 5 µl YFP-luc lentivirus, mixed, centrifuged at 300 x g for 90 min at 28$^{o}$C, and gently resuspended. Media was exchanged the following day to remove additives. YFP- luc infected cells were expanded and plated into 12-well plates. 2 µl of luciferin was added to the cells and measured after a 1 minute incubation on the IVIS Spectrum Imaging System to confirm bioluminescence *in vitro*. Cells were also expanded and implanted as $5x10^6$ and $10x10^6$ cell flanks into nude mice and were imaged on the IVIS 4 days later to confirm bioluminescence *in vivo*.

*In vitro drug testing*

The BaF3 naïve, $BaF3^{ALK}$, and $BaF3^{NTRK}$ lines were expanded for growth-curve determination over the course of 3 days, measured by Cell Titer Glo (Promega) luminescence on the Pherastar plate reader (BMG). $5x10^4$ cells per 0.5 ml media each were plated into 24-well plates and were dosed in triplicate with a 1:10 dilution series between 0-10,000 nM final concentration of ceritinib, ensartinib, or larotrectinib. Cells were measured after 3 days with Cell Titer Glo on the Phearstar plate reader to determine drug curves.

**Preclinical Testing**

*SJMEL031086_X4*

Melanoma orthotopic xenografts were created by injecting luciferase labeled SJMEL031086_X4 patient tumor cells, dissociated and passaged according to the MAST protocol, into recipient athymic nude mice using a subcutaneous injection technique. Mice were screened weekly by Xenogen and the bioluminescence was measured. Mice were enrolled in the study after achieving a target bioluminescence signal of $10^7$ photons/sec/$cm^2$ or a palpable tumor, and chemotherapy was started the following Monday.

Mice were randomized to the following treatment groups:

Control, Palbociclib, Trametinib, MAP3K8i, Trametinib+MAP3K8i, Selumetinib, Ulixertinib, LY3023414

The following doses and schedules were used for each drug:

| | |
|---|---|
| Palbociclib | 10 mg/kg oral gavage once daily on days 1-21 continuously |
| Trametinib | 0.1 mg/kg oral gavage once daily on days 1-21 continuously |
| MAP3K8i | 25 mg/kg oral gavage twice daily on days 1-21 continuously |
| Selumetinib | 1.25 mg/kg oral gavage twice daily on days 1-21 continuously |
| Ulixertinib | 50 mg/kg oral gavage twice daily on days 1-21 continuously |
| LY3023414 | 15 mg/kg oral gavage twice daily on days 1-21 continuously |

Mice received 4 courses of chemotherapy (3 weeks per course) and bioluminescence was monitored weekly and at the end of therapy. In addition to caliper measurements, thermal images were obtained for 3D tumor volume measurement weekly using the BioVolume (Fuel3D) system according to manufacture instructions. Disease response was classified according to bioluminescence signal. Mice with a signal of $10^5$ photons/sec/$cm^2$ or less (similar to background) were classified as complete response, $10^5$-$10^6$ photons/sec/$cm^2$ as partial response, $10^7$-$10^8$ photons/sec/$cm^2$ (similar to enrollment signal) as stable disease, and greater than $10^8$ photons/sec/$cm^2$ as progressive disease. Mice with tumor burden at any time greater than 20% of body weight were also classified as progressive disease. Mice were monitored daily while receiving chemotherapy.

*SJMEL030083_X1*

Melanoma orthotopic xenografts were created by injecting luciferase labeled SJMEL030083_X1 patient tumor cells, dissociated and passaged according to the MAST protocol, into recipient athymic nude mice using a subcutaneous injection technique. Mice were screened weekly by Xenogen and the bioluminescence was measured. Mice were enrolled in the study after achieving a target bioluminescence signal of $10^7$ photons/sec/$cm^2$ or a palpable tumor, and chemotherapy was started the following Monday.

Mice were randomized to the following treatment groups:

Control, Palbociclib, Trametinib, MAP3K8, Trametinib+MAP3K8i, Selumetinib, Ulixertinib, LY3023414

The following doses and schedules were used for each drug:

| | |
|---|---|
| Palbociclib | 10 mg/kg oral gavage once daily on days 1-21 continuously |
| Trametinib | 0.1 mg/kg oral gavage once daily on days 1-21 continuously |
| MAP3K8i | 25 mg/kg oral gavage twice daily on days 1-21 continuously |
| Selumetinib | 1.25 mg/kg oral gavage twice daily on days 1-21 continuously |
| Ulixertinib | 50 mg/kg oral gavage twice daily on days 1-21 continuously |
| LY3023414 | 15 mg/kg oral gavage twice daily on days 1-21 continuously |

Mice received 4 courses of chemotherapy (3 weeks per course) and bioluminescence was monitored weekly and at the end of therapy. In addition to caliper measurements, thermal images were obtained for 3D tumor volume measurement weekly using the BioVolume (Fuel3D) system according to manufacture instructions. Disease response was classified according to bioluminescence signal. Mice with a signal of $10^5$ photons/sec/$cm^2$ or less (similar to background) were classified as complete response, $10^5$-$10^6$ photons/sec/$cm^2$ as partial response, $10^7$-$10^8$ photons/sec/$cm^2$ (similar to enrollment signal) as stable disease, and greater than $10^8$ photons/sec/$cm^2$ as progressive disease. Mice with tumor burden at any time greater than 20% of body weight were also classified as progressive disease. Mice were monitored daily while receiving chemotherapy.

*SJDSRCT046155_X1*

Desmoplastic small round cell tumor (DSRCT) xenografts were created by injecting luciferase labeled SJDSRCT046155_X1 patient tumor cells, dissociated and passaged according to the MAST protocol, into recipient athymic nude mice using a subcutaneous injection technique. Mice were screened weekly by Xenogen and the bioluminescence was measured. Mice were enrolled in the study after achieving a target bioluminescence signal of $10^7$ photons/sec/$cm^2$ or a palpable tumor, and chemotherapy was started the following Monday.

Mice were randomized to the following treatment groups:

Control, Tazemetostat

The following doses and schedules were used for each drug:

| | |
|---|---|
| Tazemetostat | 225 mg/kg oral gavage twice daily on days 1-21 continuously |

Mice received 4 courses of chemotherapy (3 weeks per course) and bioluminescence was monitored weekly and at the end of therapy. Disease response was classified according to bioluminescence signal. Mice with a signal of $10^5$ photons/sec/cm$^2$ or less (similar to background) were classified as complete response, $10^5$-$10^6$ photons/sec/cm$^2$ as partial response, $10^7$-$10^8$ photons/sec/cm$^2$ (similar to enrollment signal) as stable disease, and greater than $10^8$ photons/sec/cm$^2$ as progressive disease. Mice with tumor burden at any time greater than 20% of body weight were also classified as progressive disease. Mice were monitored daily while receiving chemotherapy.

*SJRHB046156_X1*
Rhabdomyosarcoma orthotopic xenografts were created by injecting luciferase labeled SJRHB046156_X1 patient tumor cells, dissociated and passaged according to the MAST protocol, into recipient athymic nude mice using an intramuscular injection technique. Mice were screened weekly by Xenogen and the bioluminescence was measured. Mice were enrolled in the study after achieving a target bioluminescence signal of $10^7$ photons/sec/cm$^2$ or a palpable tumor, and chemotherapy was started the following Monday.

Mice were randomized to the following treatment groups:

Control, VCR+IRN, Selumetinib

The following doses and schedules were used for each drug:

| | |
|---|---|
| Vincristine (VCR) | 0.38 mg/kg IP on days 1, 8, and 15 |
| Irinotecan (IRN) | 1.25 mg/kg IP on days 1-5, 8-12 |
| Selumetinib | 1.25 mg/kg oral gavage twice daily on days 1-21 continuously |

Mice received 4 courses of chemotherapy (3 weeks per course) and bioluminescence was monitored weekly and at the end of therapy. Disease response was classified according to bioluminescence signal. Mice with a signal of $10^5$ photons/sec/cm$^2$ or less (similar to background) were classified as complete response, $10^5$-$10^6$ photons/sec/cm$^2$ as partial response, $10^7$-$10^8$ photons/sec/cm$^2$ (similar to enrollment signal) as stable disease, and greater than $10^8$ photons/sec/cm$^2$ as progressive disease. Mice with tumor burden at any time greater than 20% of body weight were also classified as progressive disease. Mice were monitored daily while receiving chemotherapy.

*SJRHB012405_X1*
Rhabdomyosarcoma orthotopic xenografts were created by injecting luciferase labeled SJRHB012405_X1 patient tumor cells, dissociated and passaged according to the MAST protocol, into recipient athymic nude mice using an intramuscular injection technique. Mice were screened weekly by Xenogen and the bioluminescence was measured. Mice were enrolled in the study after achieving a target bioluminescence signal of $10^7$ photons/sec/cm$^2$ or a palpable tumor, and chemotherapy was started the following Monday.

Mice were randomized to the following treatment groups: Control,

VCR+IRN, LY3023414

The following doses and schedules were used for each drug:

| | |
|---|---|
| Vincristine (VCR) | 0.38 mg/kg IP on days 1, 8, and 15 |
| Irinotecan (IRN) | 1.25 mg/kg IP on days 1-5, 8-12 |
| LY3023414 | 15 mg/kg oral gavage twice daily on days 1-21 continuously |

Mice received 4 courses of chemotherapy (3 weeks per course) and bioluminescence was monitored weekly and at the end of therapy. Disease response was classified according to bioluminescence signal. Mice with a signal of $10^5$ photons/sec/cm$^2$ or less (similar to background) were classified as complete response, $10^5$-$10^6$ photons/sec/cm$^2$ as partial response, $10^7$-$10^8$ photons/sec/cm$^2$ (similar to enrollment signal) as stable disease, and greater than $10^8$ photons/sec/cm$^2$ as progressive disease. Mice with tumor burden at any time greater than 20% of body weight were also classified as progressive disease. Mice were monitored daily while receiving chemotherapy.

*SJRHB011_X*

Rhabdomyosarcoma orthotopic xenografts were created by injecting luciferase labeled SJRHB011_X patient tumor cells, dissociated and passaged according to the MAST protocol, into recipient athymic nude mice using an intramuscular injection technique. Mice were screened weekly by Xenogen and the bioluminescence was measured. Mice were enrolled in the study after achieving a target bioluminescence signal of $10^7$ photons/sec/cm$^2$ or a palpable tumor, and chemotherapy was started the following Monday.

Mice were randomized to the following treatment groups:

Control, VCR+IRN, Selumetinib, Selumetinib+VCR+IRN, Erdafitinib, Erdafitinib+VCR+IRN

The following doses and schedules were used for each drug:

| | |
|---|---|
| Vincristine (VCR) | 0.38 mg/kg IP on days 1, 8, and 15 |
| Irinotecan (IRN) | 1.25 mg/kg IP on days 1-5, 8-12 |
| Selumetinib | 1.25 mg/kg oral gavage twice daily on days 1-21 continuously |
| Erdafitinib | 12.5 mg/kg oral gavage twice daily on days 1-21 continuously |

Mice received 4 courses of chemotherapy (3 weeks per course) and bioluminescence was monitored weekly and at the end of therapy. Disease response was classified according to bioluminescence signal. Mice with a signal of $10^5$ photons/sec/cm$^2$ or less (similar to background) were classified as complete response, $10^5$-$10^6$ photons/sec/cm$^2$ as partial response, $10^7$-$10^8$ photons/sec/cm$^2$ (similar to enrollment signal) as stable disease, and greater than $10^8$ photons/sec/cm$^2$ as progressive disease. Mice with tumor burden at any time greater than 20% of body weight were also classified as progressive disease. Mice were monitored daily while receiving chemotherapy.

*SJRHB012_Y*
Rhabdomyosarcoma orthotopic xenografts were created by injecting luciferase labeled SJRHB012_Y patient tumor cells, dissociated and passaged according to the MAST protocol, into recipient athymic nude mice using an intramuscular injection technique. Mice were screened weekly by Xenogen and the bioluminescence was measured. Mice were enrolled in the study after achieving a target bioluminescence signal of $10^7$ photons/sec/cm$^2$ or a palpable tumor, and chemotherapy was started the following Monday.

Mice were randomized to the following treatment groups:

Control, VCR+IRN, Selumetinib, Selumetinib+VCR+IRN

The following doses and schedules were used for each drug:

| | |
|---|---|
| Vincristine (VCR) | 0.38 mg/kg IP on days 1, 8, and 15 |
| Irinotecan (IRN) | 1.25 mg/kg IP on days 1-5, 8-12 |
| Selumetinib | 1.25 mg/kg oral gavage twice daily on days 1-21 continuously |

Mice received 4 courses of chemotherapy (3 weeks per course) and bioluminescence was monitored weekly and at the end of therapy. Disease response was classified according to bioluminescence signal. Mice with a signal of $10^5$ photons/sec/cm$^2$ or less (similar to background) were classified as complete response, $10^5$-$10^6$ photons/sec/cm$^2$ as partial response, $10^7$-$10^8$ photons/sec/cm$^2$ (similar to enrollment signal) as stable disease, and greater than $10^8$ photons/sec/cm$^2$ as progressive disease. Mice with tumor burden at any time greater than 20% of body weight were also classified as progressive disease. Mice were monitored daily while receiving chemotherapy.

*SJRHB012_Z*
Rhabdomyosarcoma orthotopic xenografts were created by injecting luciferase labeled SJRHB012_Z patient tumor cells, dissociated and passaged according to the MAST protocol, into recipient athymic nude mice using an intramuscular injection technique. Mice were screened weekly by Xenogen and the bioluminescence was measured. Mice were enrolled in the study after achieving a target bioluminescence signal of $10^7$ photons/sec/cm$^2$ or a palpable tumor, and chemotherapy was started the following Monday.

Mice were randomized to the following treatment groups:

Control, VCR+IRN, Selumetinib, Selumetinib+VCR+IRN, Erdafitinib, Ensartinib, Ensartinib*, Ensartinib+VCR+IRN

The following doses and schedules were used for each drug:

| | |
|---|---|
| Vincristine (VCR) | 0.38 mg/kg IP on days 1, 8, and 15 |
| Irinotecan (IRN) | 1.25 mg/kg IP on days 1-5, 8-12 |
| Selumetinib | 1.25 mg/kg oral gavage twice daily on days 1-21 continuously |
| Erdafitinib | 12.5 mg/kg oral gavage twice daily on days 1-21 continuously |
| Ensartinib | 100 mg/kg oral gavage twice daily on days 1-21 continuously |
| Ensartinib* | 50 mg/kg oral gavage twice daily on days 1-21 continuously |

*dose reduced due to toxicity

Mice received 4 courses of chemotherapy (3 weeks per course) and bioluminescence was monitored weekly and at the end of therapy. Disease response was classified according to bioluminescence signal. Mice with a signal of $10^5$ photons/sec/cm$^2$ or less (similar to background) were classified as complete response, $10^5$-$10^6$ photons/sec/cm$^2$ as partial response, $10^7$-$10^8$ photons/sec/cm$^2$ (similar to enrollment signal) as stable disease, and greater than $10^8$ photons/sec/cm$^2$ as progressive disease. Mice with tumor burden at any time greater than 20% of body weight were also classified as progressive disease. Mice were monitored daily while receiving chemotherapy.

*SJRHB013758_X1*

Rhabdomyosarcoma orthotopic xenografts were created by injecting luciferase labeled SJRHB013758_X1 patient tumor cells, dissociated and passaged according to the MAST protocol, into recipient athymic nude mice using an intramuscular injection technique.
Mice were screened weekly by Xenogen and the bioluminescence was measured. Mice were enrolled in the study after achieving a target bioluminescence signal of $10^7$ photons/sec/cm$^2$ or a palpable tumor, and chemotherapy was started the following Monday.

Mice were randomized to the following treatment groups:

Control, VAC, Olaparib

The following doses and schedules were used for each drug:

| | |
|---|---|
| Vincristine (V) | 0.38 mg/kg IP on days 1, 8, and 15 |
| Actinomycin D (A) | 0.5 mg/kg IP on day 1 |
| Cyclophosphamide (C) | 125 mg/kg IP on day 1 |
| Olaparib | 50 mg/kg oral gavage twice daily on days 1-21 continuously |

Mice received 4 courses of chemotherapy (3 weeks per course) and bioluminescence was monitored weekly and at the end of therapy. Disease response was classified according to bioluminescence signal. Mice with a signal of $10^5$ photons/sec/cm$^2$ or less (similar to background) were classified as complete response, $10^5$-$10^6$ photons/sec/cm$^2$ as partial response, $10^7$-$10^8$ photons/sec/cm$^2$ (similar to enrollment signal) as stable disease, and greater than $10^8$ photons/sec/cm$^2$ as progressive disease. Mice with tumor burden at any time greater than 20% of body weight were also classified as progressive disease. Mice were monitored daily while receiving chemotherapy.

*SJOS001108_X1*

Osteosarcoma orthotopic xenografts were created by injecting unlabeled SJOS001108_X1 patient tumor cells, dissociated and passaged according to the MAST protocol, into recipient athymic nude mice using a bone marrow injection technique. Mice were then observed weekly until a palpable tumor was detected and subsequently started chemotherapy the following Monday.

Mice were randomized to the following treatment groups:

Control, Gemcitabine+Docetaxel, LY3023414

The following doses and schedules were used for each drug:

| | |
|---|---|
| Gemcitabine | 30 mg/kg IP on days 2 and 9 |
| Docetaxel | 5 mg/kg IP on day 9 |
| LY3023414 | 15 mg/kg oral gavage twice daily on days 1-21 continuously |

Mice received 4 courses of chemotherapy (3 weeks per course). Caliper measurements of the tumors were performed the week prior to enrollment and weekly thereafter. Mice with tumor burden at any time greater than 20% of body weight were classified as progressive disease. Mice were monitored daily while receiving chemotherapy.

*SJOS046149_X1*
Osteosarcoma orthotopic xenografts were created by injecting unlabeled SJOS046149_X1 patient tumor cells, dissociated and passaged according to the MAST protocol, into recipient athymic nude mice using a bone marrow injection technique. Mice were then observed weekly until a palpable tumor was detected and subsequently started chemotherapy the following Monday.

Mice were randomized to the following treatment groups:

Gemcitabine+Docetaxel, Selumetinib

The following doses and schedules were used for each drug:

| | |
|---|---|
| Gemcitabine | 30 mg/kg IP on days 2 and 9 |
| Docetaxel | 5 mg/kg IP on day 9 |
| Selumetinib | 1.25 mg/kg oral gavage twice daily on days 1-21 continuously |

Mice received 4 courses of chemotherapy (3 weeks per course). Caliper measurements of the tumors were performed the week prior to enrollment and weekly thereafter. Mice with tumor burden at any time greater than 20% of body weight were classified as progressive disease. Mice were monitored daily while receiving chemotherapy.

*SJMRT015723_X1*
Rhabdoid tumor orthotopic xenografts were created by injecting luciferase labeled SJMRT015723_X1 patient tumor cells, dissociated and passaged according to MAST protocol, into recipient athymic nude mice in the peri-adrenal location. Mice were screened weekly by Xenogen and the bioluminescence was measured. Mice were enrolled in the study after achieving a target bioluminescence signal of $10^7$ photons/sec/$cm^2$ or a palpable tumor, and chemotherapy was started

the following Monday.

Mice were randomized to the following treatment groups:

Control, Tazemetostat

The following doses and schedules were used for each drug:

| | |
|---|---|
| Tazemetostat | 225 mg/kg oral gavage twice daily on days 1-21 |

Mice received 4 courses of chemotherapy (3 weeks per course) and bioluminescence was monitored weekly and at the end of therapy. Disease response was classified according to bioluminescence signal. Mice with a signal of $10^5$ photons/sec/cm$^2$ or less (similar to background) were classified as complete response, $10^5$-$10^6$ photons/sec/cm$^2$ as partial response, $10^7$-$10^8$ photons/sec/cm$^2$ (similar to enrollment signal) as stable disease, and greater than $10^8$ photons/sec/cm$^2$ as progressive disease. Mice with tumor burden at any time greater than 20% of body weight were also classified as progressive disease. Mice were monitored daily while receiving chemotherapy.

*SJNBL046148_X1*
Neuroblastoma orthotopic xenografts were created by injecting unlabeled SJNBL046148_X1 patient tumor cells, dissociated and passaged according to MAST protocol, into recipient athymic nude mice in the peri-adrenal location. Tumor measurement was performed weekly via ultrasound. Mice were enrolled once tumor reached volume >8mm$^3$ on ultrasound and chemotherapy was started the following Monday.

Mice were randomized to the following treatment groups:

Control, Selumetinib, Ensartinib

The following doses and schedules were used for each drug:

| | |
|---|---|
| Selumetinib | 1.25 mg/kg oral gavage twice daily on days 1-21 continuously |
| Ensartinib | 100 mg/kg oral gavage twice daily on days 1-21 continuously |

Mice received 4 courses of chemotherapy (3 weeks per course) and tumor volume was monitored weekly. Mice with tumor burden at any time greater than 20% of body weight were classified as progressive disease. Mice were monitored daily while receiving chemotherapy.

*ES-8*

ES-8 cell line cells were originally obtained from Chris Morton from SJCRH. Clinical information: obtained from 10 year old male from humerus tumor. Mutation status: Type II EWS-FLI1 translocation (EWSR1 exon_8 to FLI1 exon_8), STAG2 (-) by IHC, P53 (+) by IHC in ~70% of cells. STR verification was completed prior to injection of cells. EWS orthotopic xenografts were created by injecting luciferase labeled cells into recipient athymic nude mice using a bone marrow injection technique. Mice were screened weekly by Xenogen and the bioluminescence was measured. Mice were enrolled in the study after achieving a target bioluminescence signal of $10^7$ photons/sec/cm$^2$ or a palpable tumor, and chemotherapy was started the following Monday.

Mice were randomized to the following treatment groups:

Control, IRN+TMZ, TAL+IRN, L-IRN, TAL+L-IRN, TAL*+L-IRN+TMZ*

The following doses and schedules were used for each drug:

| | |
|---|---|
| Irinotecan (IRN) | 1.25 mg/kg IP on days 1-5, 8-12 |
| Talazoparib (TAL) | 0.125 mg/kg oral gavage twice daily on days 1-5, 8-12 |
| Talazoparib (TAL*) | 0.1 mg/kg oral gavage twice daily on days 1-5, 8-12 when given with L-IRN+TMZ |
| Temozolomide (TMZ) | 33 mg/kg oral gavage once daily on days 1-5 |
| Temozolomide (TMZ*) | 10 mg/kg oral gavage once daily on days 1-5 when given with TAL+L-IRN |
| Liposomal IRN (L-IRN) | 5 mg/kg tail vein given once on day 1 |

Mice received 4 courses of chemotherapy (3 weeks per course) and bioluminescence was monitored weekly and at the end of therapy. Disease response was classified according to bioluminescence signal. Mice with a signal of $10^5$ photons/sec/cm$^2$ or less (similar to background) were classified as complete response, $10^5$-$10^6$ photons/sec/cm$^2$ as partial response, $10^7$-$10^8$ photons/sec/cm$^2$ (similar to enrollment signal) as stable disease, and greater than $10^8$ photons/sec/cm$^2$ as progressive disease. Mice with tumor burden at any time greater than 20% of body weight were also classified as progressive disease. Mice were monitored daily while receiving chemotherapy.

*SJEWS030393_X1*
Ewing's sarcoma orthotopic xenografts were created by injecting luciferase labeled SJEWS030393_X1 patient tumor cells, dissociated and passaged according to the MAST protocol, into recipient athymic nude mice using a intrafemoral injection technique.
Mice were screened weekly by Xenogen and the bioluminescence was measured. Mice were enrolled in the study after achieving a target bioluminescence signal of $10^7$ photons/sec/cm$^2$ or a palpable tumor, and chemotherapy was started the following Monday.

Mice were randomized to the following treatment groups:

Control, Liposomal Irinotecan + Temozolomide, Irinotecan + Temozolomide, Talazoparib + Irinotecan, Talazoparib + Liposomal Irinotecan, Talazoparib + Temozolomide + Irinotecan, Talazoparib + Temozolomide + Liposomal Irinotecan

The following doses and schedules were used for each drug:

| | |
|---|---|
| Talazoparib | 0.1 mg/kg oral gavage twice daily on days 1-5 and 8-12 |
| Temozolomide | 10 mg/kg oral gavage once daily on days 1-5 |

| | |
|---|---|
| Irinotecan | 1.25 mg/kg intraperitoneal once daily on days 1-5 and 8-12 |
| Liposomal Irinotecan | 5 mg/kg intravenous once daily on days 1 and 8 |

Mice received 4 courses of chemotherapy (3 weeks per course) and bioluminescence was monitored weekly and at the end of therapy. Caliper measurements of the tumors were performed the week prior to enrollment and weekly thereafter. Disease response was classified according to bioluminescence signal. Mice with a signal of $10^5$ photons/sec/cm$^2$ or less (similar to background) were classified as complete response, $10^5$-$10^6$ photons/sec/cm$^2$ as partial response, $10^7$-$10^8$ photons/sec/cm$^2$ (similar to enrollment signal) as stable disease, and greater than $10^8$ photons/sec/cm$^2$ as progressive disease. Mice with tumor burden at any time greater than 20% of body weight were also classified as progressive disease. Mice were monitored daily while receiving chemotherapy.

*SJEWS063834_X2*

Ewing's sarcoma orthotopic xenografts were created by injecting luciferase labeled SJEWS063834_X2 patient tumor cells, dissociated and passaged according to the MAST protocol, into recipient athymic nude mice using a intrafemoral injection technique.

Mice were screened weekly by Xenogen and the bioluminescence was measured. Mice were enrolled in the study after achieving a target bioluminescence signal of $10^7$ photons/sec/cm$^2$ or a palpable tumor, and chemotherapy was started the following Monday.

Mice were randomized to the following treatment groups:

Control, Liposomal Irinotecan + Temozolomide, Irinotecan + Temozolomide, Talazoparib + Irinotecan, Talazoparib + Liposomal Irinotecan, Talazoparib + Temozolomide + Irinotecan, Talazoparib + Temozolomide + Liposomal Irinotecan

The following doses and schedules were used for each drug:

| | |
|---|---|
| Talazoparib | 0.1 mg/kg oral gavage twice daily on days 1-5 and 8-12 |
| Temozolomide | 10 mg/kg oral gavage once daily on days 1-5 |
| Irinotecan | 1.25 mg/kg intraperitoneal once daily on days 1-5 and 8-12 |
| Liposomal Irinotecan | 5 mg/kg intravenous once daily on days 1 and 8 |

Mice received 4 courses of chemotherapy (3 weeks per course) and bioluminescence was monitored weekly and at the end of therapy. Caliper measurements of the tumors were performed the week prior to enrollment and weekly thereafter. Disease response was classified according to bioluminescence signal. Mice with a signal of $10^5$ photons/sec/cm$^2$ or less (similar to background) were classified as complete response, $10^5$-$10^6$ photons/sec/cm$^2$ as partial response, $10^7$-$10^8$ photons/sec/cm$^2$ (similar to enrollment signal) as stable disease, and greater than $10^8$ photons/sec/cm$^2$ as progressive disease. Mice with tumor burden at any time greater than 20% of body weight were also classified as progressive disease. Mice were monitored daily while receiving chemotherapy.

*SJEWS063829_X1*

Ewing's sarcoma orthotopic xenografts were created by injecting luciferase labeled SJEWS063829_X1 patient tumor cells, dissociated and passaged according to the MAST protocol, into recipient athymic nude mice using a intrafemoral injection technique.

Mice were screened weekly by Xenogen and the bioluminescence was measured. Mice were enrolled in the study after achieving a target bioluminescence signal of $10^7$ photons/sec/cm$^2$ or a palpable tumor, and chemotherapy was started the following Monday.

Mice were randomized to the following treatment groups:

Control, Liposomal Irinotecan + Temozolomide, Irinotecan + Temozolomide, Talazoparib + Irinotecan, Talazoparib + Liposomal Irinotecan, Talazoparib + Temozolomide + Irinotecan, Talazoparib + Temozolomide + Liposomal Irinotecan

The following doses and schedules were used for each drug:

| | |
|---|---|
| Talazoparib | 0.1 mg/kg oral gavage twice daily on days 1-5 and 8-12 |
| Temozolomide | 10 mg/kg oral gavage once daily on days 1-5 |
| Irinotecan | 1.25 mg/kg intraperitoneal once daily on days 1-5 and 8-12 |
| Liposomal Irinotecan | 5 mg/kg intravenous once daily on days 1 and 8 |

Mice received 4 courses of chemotherapy (3 weeks per course) and bioluminescence was monitored weekly and at the end of therapy. Caliper measurements of the tumors were performed the week prior to enrollment and weekly thereafter. Disease response was classified according to bioluminescence signal. Mice with a signal of $10^5$ photons/sec/cm$^2$ or less (similar to background) were classified as complete response, $10^5$-$10^6$ photons/sec/cm$^2$ as partial response, $10^7$-$10^8$ photons/sec/cm$^2$ (similar to enrollment signal) as stable disease, and greater than $10^8$ photons/sec/cm$^2$ as progressive disease. Mice with tumor burden at any time greater than 20% of body weight were also classified as progressive disease. Mice were monitored daily while receiving chemotherapy.

*SJRHB000026_X1*

Two separately randomized studies using SJRHB000026_X1 RMS orthotopic xenografts were performed. Luciferase labeled cells from SJRHB000026_X1 were injected into recipient athymic nude mice using an intramuscular injection technique. Mice were screened weekly by Xenogen and the bioluminescence was measured. Mice were enrolled in the study after achieving a target bioluminescence signal of $10^7$ photons/sec/cm$^2$ or a palpable tumor, and chemotherapy was started the following Monday.

Study 1: Mice were randomized to the following treatment groups and dose schedules:

| |
|---|
| Control: vehicle/untreated |
| GDC-0084: GDC-0084 dosed at 5 mg/kg oral gavage once daily on days 1-21 |
| VCR + IRN[a]: Vincristine (VCR) dosed at 0.38 mg/kg once daily on days 1 and 8; Irinotecan (IRN) dosed at 3.125 mg/kg once daily on days 1-5 |
| VCR + L-IRN[a]: VCR dosed at 0.38 mg/kg once daily on days 1 and 8, Liposomal Irinotecan (L-IRN) dosed at 10 mg/kg once daily on day 1 |
| VCR+IRN+TMZ[a]: VCR dosed at 0.38 mg/kg once daily on days 1 and 8; IRN dosed at 3.125 mg/kg once daily on days 1-5; Temozolomide (TMZ) dosed at 33 mg/kg oral gavage once daily on days 1-5 |

| |
|---|
| VCR+IRN+TMZ[b]: VCR dosed at 0.38 mg/kg once daily on days 1 and 8; IRN dosed at 3.125 mg/kg once daily on days 1-5; TMZ dosed at 50 mg/kg oral gavage once daily on days 1-5 |
| VCR + L-IRN + TMZ: VCR dosed at 0.38 mg/kg once daily on days 1 and 8; L-IRN dosed at 10 mg/kg once daily on day 1; TMZ dosed at 50 mg/kg oral gavage once daily on days 1-5 |

Study 2: Mice were randomized to the following treatment groups and dose schedules:

| |
|---|
| Control: vehicle/untreated |
| L-IRN: L-IRN dosed at 5 mg/kg on day 1 |
| VCR+IRN[b]: VCR dosed at 0.38 mg/kg on days 1, 8, and 15; IRN dosed at 1.25 mg/kg daily on days 1-5, 8-12 |
| VCR+L-IRN[b]: VCR dosed at 0.38 mg/kg on days 1, 8, and 15; L-IRN dosed at 5 mg/kg on day 1 |
| VCR+L-IRN[c]: VCR dosed at 0.38 mg/kg on days 1, 8, and 15; L-IRN dosed at 5 mg/kg on days 1 and 8 |
| AZD1775+VCR+IRN: AZD1775 dosed at 48 mg/kg twice daily on days 1-5; VCR dosed at 0.38 mg/kg on days 1, 8, and 15; IRN dosed at 1.25 mg/kg daily on days 1-5, 8-12 |
| AZD1775+VCR+L-IRN: AZD1775 dosed at 48 mg/kg twice daily on days 1-5; VCR dosed at 0.38 mg/kg on days 1, 8, and 15; L-IRN dosed at 5 mg/kg on day 1 |

Mice in both studies received 4 courses of chemotherapy (3 weeks per course) and bioluminescence was monitored weekly and at the end of therapy. Disease response was classified according to bioluminescence signal. Mice with a signal of $10^5$ photons/sec/cm$^2$ or less (similar to background) were classified as complete response, $10^5$-$10^6$ photons/sec/cm$^2$ as partial response, $10^7$-$10^8$ photons/sec/cm$^2$ (similar to enrollment signal) as stable disease, and greater than $10^8$ photons/sec/cm$^2$ as progressive disease. Mice with tumor burden at any time greater than 20% of body weight were also classified as progressive disease. Mice were monitored daily while receiving chemotherapy.

*BaF3$^{ALK}$*

Study 1: Two cohorts of mice (n = 30 total) were injected with 1 x $10^6$ luciferase labeled BaF3- ALK cells using an intravenous (n = 15) or intramuscular (n = 15) injection technique. Mice were screened by Xenogen to confirm engraftment of labeled cells and enrolled in the study 10 days post injection.

Mice were randomized to the following treatment groups:

Control, Ensartinib, Larotrectinib

The following doses and schedules were used for each drug:

| | |
|---|---|
| Ensartinib | 100 mg/kg oral gavage once daily on days 1-21 continuously |

| | |
|---|---|
| Larotrectinib | 200 mg/kg oral gavage once daily on days 1-21 continuously |
| Control | No chemotherapy |

Mice received 2 courses of chemotherapy (3 weeks per course) and bioluminescence was monitored weekly and at the end of therapy. Disease response was classified according to bioluminescence signal. Mice with a signal of $10^5$ photons/sec/cm$^2$ or less (similar to background) were classified as complete response, $10^5$-$10^6$ photons/sec/cm$^2$ as partial response, $10^7$-$10^8$ photons/sec/cm$^2$ (similar to enrollment signal) as stable disease, and greater than $10^8$ photons/sec/cm$^2$ as progressive disease. Mice with tumor burden at any time greater than 20% of body weight were also classified as progressive disease. Mice were monitored daily while receiving chemotherapy and were harvested once tumor burden was greater than 20% of body weight or animal became moribund. At time of harvest, tumor tissue was flash frozen and fixed. Liver, spleen, and lymph nodes were also fixed for pathology.

Study 2: Mice were injected with $1.5 \times 10^6$ luciferase labeled BaF3-ALK cells using an intravenous injection technique and enrolled in the study 4 days post injection.

Mice were randomized to the following treatment groups:

Control, Ensartinib[a], Ensartinib[b], Ensartinib[c], Larotrectinib[a], Larotrectinib[b], Larotrectinib[c], Crizotinib[a], Crizotinib[b], Crizotinib[c]

The following doses and schedules were used for each drug:

| | |
|---|---|
| Ensartinib[a] | 25 mg/kg oral gavage twice daily on days 1-21 continuously |
| Ensartinib[b] | 50 mg/kg oral gavage twice daily on days 1-21 continuously |
| Ensartinib[c] | 100 mg/kg oral gavage twice daily on days 1-21 continuously |
| Larotrectinib[a] | 30 mg/kg oral gavage twice daily on days 1-21 continuously |
| Larotrectinib[b] | 60 mg/kg oral gavage twice daily on days 1-21 continuously |
| Larotrectinib[c] | 200 mg/kg oral gavage twice daily on days 1-21 continuously |
| Crizotinib[a] | 25 mg/kg oral gavage twice daily on days 1-21 continuously |
| Crizotinib[b] | 50 mg/kg oral gavage twice daily on days 1-21 continuously |
| Crizotinib[c] | 100 mg/kg oral gavage twice daily on days 1-21 continuously |

Mice received 1 course of chemotherapy (3 weeks per course) and bioluminescence was monitored weekly and at the end of therapy. Disease response was classified according to bioluminescence signal. Mice with a signal of $10^5$ photons/sec/cm$^2$ or less (similar to background) were classified as complete response, $10^5$-$10^6$ photons/sec/cm$^2$ as partial response, $10^7$-$10^8$ photons/sec/cm$^2$ (similar to enrollment signal) as stable disease, and greater than $10^8$ photons/sec/cm$^2$ as progressive disease. Mice with tumor burden at any time greater than 20% of body weight were also classified as progressive disease. Mice were monitored daily while receiving chemotherapy and were harvested once tumor burden was greater than 20% of body weight or animal became moribund. At time of harvest, tumor tissue was flash frozen and fixed. Liver, spleen, and lymph nodes were also fixed for pathology.

*$BaF3^{NTRK}$*

Study 1: Two cohorts of mice (n = 30 total) were injected with 1 x $10^6$ luciferase labeled BaF3-NTRK cells using an intravenous (n = 15) or intramuscular (n = 15) injection technique. Mice were screened by Xenogen to confirm engraftment of labeled cells and enrolled in the study 10 days post injection.

Mice were randomized to the following treatment groups:

Control, Ensartinib, Larotrectinib

The following doses and schedules were used for each drug:

| | |
|---|---|
| Ensartinib | 100 mg/kg oral gavage once daily on days 1-21 continuously |
| Larotrectinib | 200 mg/kg oral gavage once daily on days 1-21 continuously |
| Control | No chemotherapy |

Mice received 4 courses of chemotherapy (3 weeks per course) and bioluminescence was monitored weekly and at the end of therapy. Disease response was classified according to bioluminescence signal. Mice with a signal of $10^5$ photons/sec/$cm^2$ or less (similar to background) were classified as complete response, $10^5$-$10^6$ photons/sec/$cm^2$ as partial response, $10^7$-$10^8$ photons/sec/$cm^2$ (similar to enrollment signal) as stable disease, and greater than $10^8$ photons/sec/$cm^2$ as progressive disease. Mice with tumor burden at any time greater than 20% of body weight were also classified as progressive disease. Mice were monitored daily while receiving chemotherapy and were harvested once tumor burden was greater than 20% of body weight or became moribund. At time of harvest, tumor tissue was flash frozen and fixed. Liver, spleen, and lymph nodes were also fixed for pathology.

Study 2: Mice were injected with 1.5 x $10^6$ luciferase labeled BaF3-NTRK cells using an intravenous injection technique and enrolled in the study 4 days post injection.

Mice were randomized to the following treatment groups:

Control, Ensartinib[a], Ensartinib[b], Ensartinib[c], Larotrectinib[a], Larotrectinib[b], Larotrectinib[c], Crizotinib[a], Crizotinib[b], Crizotinib[c]

The following doses and schedules were used for each drug:

| | |
|---|---|
| Ensartinib[a] | 25 mg/kg oral gavage twice daily on days 1-21 continuously |
| Ensartinib[b] | 50 mg/kg oral gavage twice daily on days 1-21 continuously |
| Ensartinib[c] | 100 mg/kg oral gavage twice daily on days 1-21 continuously |
| Larotrectinib[a] | 30 mg/kg oral gavage twice daily on days 1-21 continuously |
| Larotrectinib[b] | 60 mg/kg oral gavage twice daily on days 1-21 continuously |
| Larotrectinib[c] | 200 mg/kg oral gavage twice daily on days 1-21 continuously |
| Crizotinib[a] | 25 mg/kg oral gavage twice daily on days 1-21 continuously |
| Crizotinib[b] | 50 mg/kg oral gavage twice daily on days 1-21 continuously |
| Crizotinib[c] | 100 mg/kg oral gavage twice daily on days 1-21 continuously |

Mice received 1 course of chemotherapy (3 weeks per course) and bioluminescence was monitored weekly and at the end of therapy. Disease response was classified according to bioluminescence signal. Mice with a signal of $10^5$ photons/sec/cm$^2$ or less (similar to background) were classified as complete response, $10^5$-$10^6$ photons/sec/cm$^2$ as partial response, $10^7$-$10^8$ photons/sec/cm$^2$ (similar to enrollment signal) as stable disease, and greater than $10^8$ photons/sec/cm$^2$ as progressive disease. Mice with tumor burden at any time greater than 20% of body weight were also classified as progressive disease. Mice were monitored daily while receiving chemotherapy and were harvested once tumor burden was greater than 20% of body weight or animal became moribund. At time of harvest, tumor tissue was flash frozen and fixed. Liver, spleen, and lymph nodes were also fixed for pathology.

*3T3- NTRK*

3T3-NTRK cells were received from Dr. Filemon Dela Cruz from Memorial Sloan Kettering Cancer Center. Mice were injected with 1.5 x $10^6$ luciferase labeled 3T3-NTRK cells using an intravenous injection technique and enrolled in the study 4 days post injection.

Mice were randomized to the following treatment groups:

Control, Ensartinib[a], Ensartinib[b], Ensartinib[c], Larotrectinib[a], Larotrectinib[b], Larotrectinib[c], Crizotinib[a], Crizotinib[b], Crizotinib[c]

The following doses and schedules were used for each drug:

| | |
|---|---|
| Ensartinib[a] | 25 mg/kg oral gavage twice daily on days 1-21 continuously |
| Ensartinib[b] | 50 mg/kg oral gavage twice daily on days 1-21 continuously |
| Ensartinib[c] | 100 mg/kg oral gavage twice daily on days 1-21 continuously |
| Larotrectinib[a] | 30 mg/kg oral gavage twice daily on days 1-21 continuously |
| Larotrectinib[b] | 60 mg/kg oral gavage twice daily on days 1-21 continuously |
| Larotrectinib[c] | 200 mg/kg oral gavage twice daily on days 1-21 continuously |
| Crizotinib[a] | 25 mg/kg oral gavage twice daily on days 1-21 continuously |
| Crizotinib[b] | 50 mg/kg oral gavage twice daily on days 1-21 continuously |
| Crizotinib[c] | 100 mg/kg oral gavage twice daily on days 1-21 continuously |

Mice received 1 course of chemotherapy (3 weeks per course) and bioluminescence was monitored weekly and at the end of therapy. Disease response was classified according to bioluminescence signal. Mice with a signal of $10^5$ photons/sec/cm$^2$ or less (similar to background) were classified as complete response, $10^5$-$10^6$ photons/sec/cm$^2$ as partial response, $10^7$-$10^8$ photons/sec/cm$^2$ (similar to enrollment signal) as stable disease, and greater than $10^8$ photons/sec/cm$^2$ as progressive disease. Mice with tumor burden at any time greater than 20% of body weight were also classified as progressive disease. Mice were monitored daily while receiving chemotherapy and were harvested once tumor burden was greater than 20% of body weight or animal became moribund. At time of harvest, tumor tissue was flash frozen and fixed. Liver, spleen, and lymph nodes were also fixed for pathology.

*SJIFS032941_X1*

Mice were injected with 1 x $10^6$ MAST 849 infantile fibrosarcoma cells using a subcutaneous injection technique. Mice were then observed weekly until visible tumor was detected in all mice and chemotherapy was started the following Monday.

Mice were randomized to the following treatment groups:

Control, Larotrectinib

The following doses and schedules were used for each drug:

| | |
|---|---|
| Larotrectinib | 30 mg/kg oral gavage twice daily on days 1-21 continuously |

Mice received 2 courses of chemotherapy (3 weeks per course). Caliper measurements of the tumors were performed weekly during and after completion of chemotherapy. Mice with tumor burden at any time greater than 20% of body weight were classified as progressive disease.

*Preclinical Statistics*
P-values for head-to-head survival comparisons of all treatment groups were calculated using a log-rank test for time to event using GraphPad Prism software.

**Xenogen Imaging and Quantification**

Mice were given intraperitoneal injections of Firefly D-Luciferin (Caliper Life Sciences 3 mg/mouse). Bioluminescent images were taken five minutes later using the IVIS® 200 imaging system. Anesthesia was administered throughout image acquisition (isoflurane 1.5% in $O_2$ delivered at 2 liters/min). The Living Image 4.3 software (Caliper Life Sciences) was used to generate a standard region of interest (ROI) encompassing the largest tumor at maximal bioluminescence signal. The identical ROI was used to determine the average radiance (photons/s/cm$^2$/sr) for all xenografts.
